# Sex-specific landscapes of crossover and non-crossover recombination in coppery titi monkeys (*Plecturocebus cupreus*)

**DOI:** 10.64898/2026.01.11.698870

**Authors:** Cyril J. Versoza, Karen L. Bales, Jeffrey D. Jensen, Susanne P. Pfeifer

## Abstract

Along with germline mutations, meiotic recombination plays a fundamental role in shaping genetic diversity and thus directly influences a species’ potential adaptive response to environmental change, amongst other features. Despite the recombination landscape being of central importance for a variety of questions in molecular evolution, the genome-wide distribution and frequency of recombination remains to be elucidated in many non-human primate species. Utilizing novel high-coverage genomic data from three multi-sibling families, we here provide the first estimates of the rates and patterns of crossover and non-crossover recombination in coppery titi monkeys (*Plecturocebus cupreus*) — a socially monogamous, pair-bonded primate that serves as an important model in behavioral research. Consistent with haplorrhines, crossover and non-crossover recombination in this platyrrhine are frequently localized at PRDM9-mediated hotspots, characterized by a 15-mer binding motif with substantial similarities to the degenerate 13-mer motif found in humans. The sex-averaged crossover rate in coppery titi monkeys is comparable with those of other primates; however, no significant difference in recombination rates was observed between the sexes, despite a pronounced maternal age effect in the species. Similarities also exist with regards to the sex-specific genomic distribution of non-crossover events, though the minimal conversion tract lengths of extended events was observed to be considerably longer in maternally-inherited non-crossovers. Taken together, these similarities and differences in the recombination landscape relative to other primates highlight the importance of incorporating species-specific rates and patterns in evolutionary models, and the resources provided here will thus serve to aid future studies in this important primate model system.

## INTRODUCTION

Along with germline mutations, meiotic recombination plays a fundamental role in generating and shaping observed genetic diversity (see the review of Johnston 2024). A requisite for the faithful segregation of homologous chromosomes during gametogenesis (Keeney 2001), recombination in primates is thought to primarily take place at PRDM9-mediated hotspots (Baudat et al. 2010; Myers et al. 2010; Parvanov et al. 2010). Localizing these hotspots by binding DNA at specific sequence motifs with its C-terminal zinc-finger domain (Ségurel et al. 2011), the histone methyltransferase PRDM9 trimethylates H3K4 and H3K36 (Powers et al. 2016), thereby creating an open chromatin environment permissive to the formation of DNA double-strand breaks (Neale and Keeney 2006). Programmed meiotic double-strand breaks are subsequently repaired using the homologous chromosome as a template (San Filippo et al. 2008), leading to either a reciprocal exchange of large segments of genetic material between the homologs (crossover; CO) or, more commonly, a non-reciprocal transfer of a short segment from a donor homolog (non-crossover; NCO) (Sun et al. 1989; and see the reviews of Wang et al. 2015; Lorenz and Mpaulo 2022).

Although a common determinant of recombination landscapes in primates, the location of PRDM9-mediated hotspots appears generally non-conserved between species (e.g., Auton et al. 2012; Stevison et al. 2016; and see Baker et al. 2017), as the rapid evolution of the PRDM9 zinc finger domain alters the recognition of, and binding specificity to, the sequence motif (Oliver et al. 2009; Myers et al. 2010; Hinch et al. 2011; Baudat et al. 2013; Baker et al. 2017). In many organisms, there also exists sexual dimorphism with regards to both the positioning and usage of recombination hotspots as well as the overall rate of recombination; for example, in humans, females generally exhibit higher rates of COs and lower rates of NCOs than males (though NCO tracts tend to be longer), whereas males display a more prominent elevation of recombination rates toward the telomeric ends of the chromosomes compared to the more uniformly distributed rates observed in females (Kong et al. 2002, 2010; Coop et al. 2008; Halldorsson et al. 2016, 2019; Palsson et al. 2025; and see Sardell and Kirkpatrick 2020).

Knowledge of the distribution and frequency of recombination is vitally important in many studies of molecular evolution and population genetics, including, for instance, in the inference of population history, the detection of genomic targets of natural selection, as well as genome-wide association and linkage studies (Johri et al. 2022). Consequently, species-specific recombination landscapes have been inferred in a variety of primates of evolutionary and biomedical interest over the past decades, ranging from the great apes (e.g., Kong et al. 2002, 2010; Auton et al. 2012; Stevison et al. 2016; Bhérer et al. 2017; Halldorsson et al. 2019), to baboons (Wall et al. 2022), rhesus macaques (Xue et al. 2016, 2020; Versoza et al. 2024; Terbot et al. 2025b), vervet monkeys (Pfeifer 2020), marmosets (Soni, Versoza et al. 2025b), and aye-ayes (Soni, Versoza, et al. 2025a; Terbot et al. 2025a; Versoza, Lloret-Villas et al. 2025; and see the review of Soni et al. 2025). Notably though, much of this previous work indirectly inferred sex-averaged recombination rates at the population-level from patterns of linkage disequilibrium (LD) along the genome, focusing by and large on the outcome of CO events. Conversely, despite occurring at much higher frequencies in the genomes of many organisms (Jeffreys and May 2004; Baudat and de Massy 2007; Cole et al. 2010; Comeron et al. 2012; Li et al. 2019; Palsson et al. 2025), NCO events remain relatively understudied, leaving our picture of recombination landscapes incomplete.

To address this gap in our knowledge, and to circumvent (potentially problematic) simplifying assumptions regarding the demographic history, effective population size, and mutation rate necessary for the application of LD-based approaches, a limited number of studies have recently begun to directly investigate CO and NCO events based on whole genome data from pedigreed individuals or from single-molecule sequences of individual sperm (or testis) samples (Coop et al. 2008; Williams et al. 2015; Halldorsson et al. 2016; Wall et al. 2022; Porsborg et al. 2024; Schweiger et al. 2024; Versoza et al. 2024; Palsson et al. 2025; Versoza, Lloret-Villas et al. 2025). However, in contrast to CO events which are relatively straightforward to identify, the detection of NCO events remains challenging (see the discussion in Li et al. 2019). Specifically, NCO detection is complicated by their short tract size (e.g., spanning ∼50-1,000 bp in humans, with recent empirical estimates suggesting that the majority of NCOs are likely associated with PRDM9-mediated recombination and exhibit a mean conversion tract length of ∼40-100 bp); a minor proportion, likely the result of a distinct process independent of PRDM9, exhibits longer mean conversion tract lengths (Jeffreys and May 2004; Odenthal-Hesse et al. 2014; Williams et al. 2015; Halldorsson et al. 2016; Hardarson et al. 2023; Porsborg et al. 2024; Schweiger et al. 2024; Palsson et al. 2025). Recent statistical inference of the underlying length distribution estimated an even lower overall mean length of ∼20 bp (Masaki and Browning 2025). Importantly, given the necessity of a heterozygous marker intersecting with the affected segment of genetic material, one would further anticipate that a greater proportion of events will be rendered undetectable as species-level heterozygosity is reduced (Charmouh et al. 2025). For levels of heterozygosity commonly observed in many primate genomes for example, NCO tracts frequently include only a single phase-informative marker (as is the case in >90% of NCOs observed in humans; Schweiger et al. 2024), thus care must be taken to reliably distinguish between genuine NCO events and sequencing / genotyping errors. Complicating detection even further are complex NCO events containing both converted and non-converted markers (Webb et al. 2008). Despite these challenges, pedigree-based approaches remain quintessential in improving our understanding of sex-specific CO and NCO events. Yet, aside from humans and the handful of non-human primates in which these approaches have been applied (Williams et al. 2015; Halldorsson et al. 2016, 2019; Wall et al. 2022; Porsborg et al. 2024; Schweiger et al. 2024; Versoza et al. 2024; Palsson et al. 2025), detailed knowledge of the rates and patterns of recombination remains lacking in many species, in part due to limited sample availability.

In order to better illuminate the nature of these landscapes across the primate clade, we here consider the coppery titi monkey (*Plecturocebus cupreus*), a socially monogamous species characterized by small pair-bonded family groups native to the Amazon forests of Brazil and Peru (Kinzey 1997). Although fewer than 10% of mammals display social monogamy and pair-bonding in the wild (Kleiman 1977; Lukas and Clutton-Brock 2013), the prevalence of these traits in humans (Gavrilets 2012) and their importance to healthy aging (House et al. 1988) has prompted the development of coppery titi monkeys as an important model system for the study of social behavior (e.g., Bales et al. 2007, 2017, 2021). To facilitate this research, a colony of coppery titi monkeys was established at the California National Primate Research Center (CNPRC) in the 1970s (Lorenz and Mason 1971). Yet, genome-wide association studies aimed at identifying genetic variants underlying these phenotypes of interest are currently hampered by a lack of both population genomic data and knowledge of the species-specific recombination landscape. Taking advantage of the recently released chromosome-level genome assembly (Pfeifer et al. 2024), together with novel high-coverage genomic data from 17 individuals belonging to three multi-sibling families, we here provide the first recombination rate estimates for the species and characterize the sex-specific spatial distribution of CO and NCO events along the genome. In addition to providing novel insights into the rates and patterns of recombination, the resulting genetic map will thus also serve as a valuable resource for future studies in this important behavioral model system.

## MATERIALS AND METHODS

### Ethics statement

This study was performed in compliance with all regulations regarding the care and use of captive primates, including the NIH Guidelines for the Care and Use of Animals and the American Society of Primatologists’ Guidelines for the Ethical Treatment of Nonhuman Primates. Procedures were approved by the UC-Davis Institutional Animal Care and Use Committee (protocol 22523).

### Samples and sequencing

Peripheral blood samples were collected from 17 coppery titi monkeys (*Plecturocebus cupreus*) belonging to three multi-sibling families housed at the CNPRC (Supplementary Figure 1; and see Supplementary Table 1 for sample information). Genomic DNA was extracted from whole blood samples using the PAXgene Blood DNA System or the QIAamp DNA Mini Kit (Qiagen, Hilden, Germany), the DNA concentration of each sample was measured using an Invitrogen Qubit Fluorometer (Thermo Fisher Scientific, Waltham, MA, US), and sample integrity and purity were assessed using Agarose Gel Electrophoresis. PCR-free libraries were prepared for all samples, checked with Qubit and real-time PCR for quantification and bioanalyzer for size distribution detection, and paired-end sequenced (2 × 150 bp) on an Illumina NovaSeq 6000 (Illumina, San Diego, CA, USA).

### Whole genome alignment

Raw reads were preprocessed using fastp v.0.24.0 (Chen et al. 2018) which detects and trims both sequencing adapters in paired-end data (*--detect_adapter_for_pe*) and polyG tails common in Illumina NovaSeq data (default minimum length: 10 bp). Additionally, fastp automatically performs quality and length filtering, removing any reads that contain more than 40% of bases with a Phred quality score < 15, more than 5 bases without call information (Ns), or an overall length of less than 15 bp. Quality-controlled reads were aligned to the coppery titi monkey reference genome (GenBank accession number: GCA_040437455.1; Pfeifer et al. 2024) using the GPU-accelerated NVIDIA Parabricks v.4.4.0-1 implementation of the Burrows-Wheeler Aligner (Li 2013), marking shorter split hits as secondary (*-M* option). Depth of coverage for each sample is provided in Supplementary Table 1.

### Variant discovery

Prior to calling variants, duplicates were marked using the NVIDIA Parabricks v.4.4.0-1 implementation of the Genome Analysis Toolkit (GATK; van der Auwera and O’Connor 2020) to eliminate spurious read coverage support (Pfeifer 2017). Given the lack of polymorphism data for the species, a first round of variant calling was performed on the high-quality, duplicate-marked mappings (*--minimum-mapping-quality 40*) of each sample, using the GATK *HaplotypeCaller* without PCR correction (*-pcr-indel-model NONE*), following the developers’ recommendations. Resulting genomic variant call files were then merged (*CombineGVCF*) to allow for joint genotyping across samples (*GenotypeGVCF*). In order to create a high-confidence dataset, annotations informative for bootstrapping were plotted for single nucleotide polymorphisms (SNPs) to determine appropriate filtering thresholds that control the transition-transversion ratio (for details, see Auton et al. 2012); subsequently any SNPs supported by less than half, or more than twice, a sample’s mean autosomal read depth (*DP*), exhibiting a genotype quality (*GQ*) less than 60, a quality-by-depth (*QD*) less than 10, a mapping quality rank sum test (*MQRankSum*) score less than –12.5, or a read position rank sum (*ReadPosRankSum*) score less than –8.0, or showing any signs of strand bias as determined by a Fisher’s exact test (*FS* > 5) or symmetric odds ratio (*SOR* > 1.5), were removed from the dataset using BCFtools *filter* v.1.14 (Danecek et al. 2021). GATK’s machine-learning-based base quality score recalibration procedure (*BaseRecalibrator* and *ApplyBQSR*) was then used to model systematic errors present in the sequencing data and to adjust the base quality scores of the original, duplicate-marked mappings to more accurately reflect the probability of a base call being incorrect. Subsequently, a second round of variant calling was performed on the recalibrated mappings as described above. To further improve accuracy, biallelic SNPs that contained genotype information for all individuals in the pedigrees were re-genotyped using Graphtyper v.2.7.2 (Eggertsson et al. 2017), a graph-based genotyping approach shown to improve variant detection, particularly in complex genomic regions. Due to the lower read depth on the sex chromosomes (X and Y), the re-genotyped dataset was limited to autosomal sites that passed all built-in sample and record level filters.

As the detection of CO and NCO events is sensitive to genotyping errors, the dataset was restricted to regions of the reference genome in which the 150bp-long sequencing reads could be confidently mapped. To this end, an accessibility mask with a stringency of 1 was generated using the *SNPable* workflow (https://lh3lh3.users.sourceforge.net/snpable.shtml), and any regions not deemed confidently mappable were excluded from downstream analyses. Additionally, the variant dataset was limited to sites consistent with the patterns of Mendelian inheritance located farther than 10 bp from the nearest insertion/deletion, with a depth of coverage of at least half, but no more than twice, the average autosomal coverage for each individual. Lastly, to ensure that heterozygous calls were of high quality, genotypes exhibiting an allelic imbalance (*p*-value > 0.01 in a two-sided exact binomial test) were filtered out following Prentout et al. 2025. Details of the variant dataset are provided in Supplementary Table 2.

### Detection of recombination events

Recombination events were detected by tracing gamete transmission across the multiple-offspring families (including a total of 11 maternal meioses and 11 paternal meioses; Supplementary Figure 1). To this end, phase-informative markers — that is, sites at which only one of the parents exhibits a heterozygous genotype — observed in a family were first used to distinguish between maternally and paternally inherited haplotypes in each offspring (see Supplementary Table 3 for the number, Supplementary Table 4 for the per-chromosome density, and Supplementary Figures 2-4 for the distribution of phase-informative markers per family). This assignment allowed for the subsequent detection of recombination events through the comparison of phase-informative markers inherited by each sibling of a multi-offspring family in order to identify switches in the parental haplotype phase (Coop et al. 2008; and see Supplementary Figure 20 in Versoza, Lloret-Villas et al. 2025 for a schematic representation of this approach). Following the methodology outlined in Prentout et al. 2025, recombination events were separated into COs (a single switch in haplotype phase) and NCOs (two phase changes occurring in close proximity and surrounded by the same parental haplotype). To guard against genotyping errors, putative CO events located within 10 phase-informative markers in an individual meiosis were grouped, keeping only COs with a single switch in haplotype, and CO events within 10 phase-informative markers from the chromosomal ends were removed. Subsequently, all putative CO events were manually inspected and those forming tight clusters (defined here as ≥ 3 COs within 10 Mb, as per Wall et al. 2022) were removed (Supplementary Table 5). In a similar vein, as putative NCO events tend to be short, frequently involving only a single phase-informative marker, such markers were required to be supported by high-confidence read mappings (defined here as read mappings with an average depth of coverage of > 20 and an average mapping quality of > 40 across individuals) and located more than 10 phase-informative markers away from the chromosomal ends; additionally, NCO tracts were required to be enclosed by ≥ 10 markers with consistent haplotype phase on either side (as per Prentout et al. 2025). Moreover, putative NCO events with a switch in haplotype phase in the same genomic location in any other meiosis, as well as those coinciding with another NCO in the surrounding 1 kb region, were excluded to avoid spurious events caused by genome assembly errors. Finally, putative NCO events overlapping structural variants known to segregate in the population (Versoza et al. 2026) were excluded from downstream analyses. A summary of the final recombination events is provided in Supplementary Table 6, with details of the detected CO and NCO events given in Supplementary Tables 7 and 8, respectively.

### Annotation of recombination events

CO events with a high resolution (< 10 kb) and NCO events with a short tract length (< 10 kb) were annotated using ANNOVAR v.2020-06-08 (Wang et al. 2010) to determine overlaps with genomic features. In addition, overlaps with CpG islands were categorized based on the annotations obtained from the reference genome (Pfeifer et al. 2024) via the *newcpgreport* function implemented in the EMBOSS suite (Rice et al. 2020; Madeira et al. 2022).

### Identification of a putative PRDM9 sequence and binding motifs

To identify a putative PRDM9 sequence in coppery titi monkeys, Liftoff v.1.6.3 (Shumate and Salzberg 2021) was used to project the human PRDM9 sequence from UniProt (entry: Q9NQV7; UniProt Consortium 2025) onto the coppery titi monkey assembly. As the highly repetitive nature and rapid evolution of PRDM9 makes the gene prone to mis-assembly, previously generated long-read data was aligned against the current reference genome assembly (Pfeifer et al. 2024) with minimap2 v.2.24 (using the recommended Oxford Nanopore pre-set ‘*-ax* map-ont’; Li 2018), the 2-Mb region encompassing the putative PRDM9 sequence identified by LiftOff was extracted using SAMtools v.1.16 (Danecek et al. 2021), and subsequently re-assembled using Flye v.2.9.1 (Kolmogorov et al. 2019). The resulting contig was then mapped back to the human PRDM9 sequence using GeneWise v.2.4.1 (Birney et al. 2004) to predict gene structure. To ensure that the extracted sequence was of high quality, it was translated into amino acids using the ExPASy platform (Gasteiger et al. 2003) and manually inspected using InterPro (Blum et al. 2025). Additionally, a BLASTX (Altschul et al. 1990) search was performed against the non-redundant protein sequences (nr) database in primates (taxid: 9443) to confirm PRDM9 identity and to manually inspect for both the presence and completeness of the KRAB, SSXRD, PR/SET, and zinc-finger domains based on identified conserved domains (Supplementary Figure 5).

To help determine whether the identified PRDM9 was likely functional, patterns of gene expression of a previously collected testis sample were compared to those of heart and adrenal glands (Pfeifer et al. 2024), with the latter two serving as negative controls as PRDM9 is expressed during meiosis. To this end, RNAseq data from these tissues were first trimmed using Trimmomatic v.0.39 (Bolger et al. 2014) in paired-end mode (with the ‘*LEADING*: 3’, *TRAILING*: 3’, and ‘*MINLEN*: 36’ flags enabled) before mapping the reads to the coppery titi monkey reference genome (Pfeifer et al. 2024) containing the putative PRDM9 contig using STAR v.2.7 (Dobin et al. 2013). Presence or absence of gene expression was assessed based on properly paired reads that mapped uniquely to the putative PRDM9 contig.

High-resolution CO events were used to identify putative PRDM9 binding motifs. In brief, following the approach described in Wijnker et al. (2013), each high-resolution CO event was expanded by 500 bp on either side to capture the entire interval likely involved in the double-strand break formation and repair. Based on these breakpoint intervals, a *de novo* motif discovery was then performed using MEME v.5.5.7 (Bailey & Elkan 1994) under the ZOOPS model with a significance threshold of 1e-05, assuming that sequence motifs exhibit a length between 5 and 15 nucleotides. To mitigate sequence composition bias during this search, a background model was generated from randomly chosen genomic regions that matched the breakpoint intervals in overall size and GC-content. Given the high repetitiveness of PRDM9 motifs in humans and other primates (e.g., Altemose et al. 2017), a permutation test was carried out using MOODS v.1.9.4.1 (Korhonen et al. 2009), comparing the frequencies of the discovered motif between the breakpoint intervals and 1,000 randomly sampled non-breakpoint intervals (*p*-value cutoff for matches: 0.05). The statistical significance of the observed enrichment in breakpoint intervals was evaluated using a Fisher’s exact test in R v.4.2.2 (R Core Team 2022).

### Estimating the power to detect NCO events

In order to estimate the power to detect NCO events present in the population genomic data analyzed in this study, 2,000 NCO events were simulated based on the tract length distributions observed in humans (Palsson et al. 2025), including 1,000 short tracts with a mean length of 113 bp and 1,000 long tracts with a mean length of 8.2 kb. For each of these two categories, power was then estimated by dividing the number of NCO events that included at least one phase-informative marker (and were hence detectable) by the number of simulated events (i.e., detectable and non-detectable).

## RESULTS AND DISCUSSION

To gain insights into the sex-specific landscapes of CO and NCO recombination in coppery titi monkeys, 9.3 million autosomal variants were called from high-coverage (∼50×) genomic data of 17 individuals belonging to three multi-sibling families housed at the CNPRC (Supplementary Figure 1; and see Supplementary Tables 1 and 2 for sample information and details of the variant dataset, respectively). Of these, an average of 4.2 million variants per family were informative with regards to the parent-of-origin of the haplotypes inherited by the offspring (Supplementary Table 3; and see Supplementary Table 4 and Supplementary Figures 2-4 for the density and distribution of the phase-informative markers across chromosomes in each family, respectively) and could thus be used to detect COs, consisting of a single switch in haplotype phase, and NCOs, consisting of short segments with two switches in haplotype phase in close proximity to one another (see “Materials and Methods” for more information).

### Sex-specific landscape of CO recombination in coppery titi monkeys

In the 22 meioses of the three multi-sibling coppery titi monkey families (Supplementary Figure 1), 579 putative autosomal CO events were detected. Out of these events, 40 were removed during manual curation as they formed tight clusters (≥ 3 COs within 10 Mb) within single meioses (Supplementary Table 5) — an observation more likely to be a technical artifact resulting from local assembly errors than biologically genuine, as CO interference reduces the likelihood of several COs in close proximity on the same chromosome (Muller 1916; and see the review of Otto and Payseur 2019). As an expected result of such CO interference, of the remaining 539 COs, CO events occurring in the same meiosis were more distant than expected based on the distribution of COs in different meioses (Supplementary Figure 6).

The 265 maternally-inherited and 274 paternally-inherited COs could be resolved to a median interval resolution of 8.1 kb and 10.9 kb, respectively (Supplementary Figure 7). The genome-wide distribution of these COs is depicted in Figure 1a (genomic locations are provided in Supplementary Table 7; and see Supplementary Figures 8-29 for the haplotype blocks observed on each chromosome in each meiosis). At the broad-scale, a pronounced enrichment of COs near telomeres was observed in males (Figure 1b), similar to other mammals (Lenormand and Dutheil 2005; Sardell and Kirkpatrick 2020). At the fine-scale, COs were highly localized in both intergenic and intronic regions (Supplementary Figure 30) as expected for PRDM9-mediated recombination (Coop et al. 2008); however, males and females displayed significant differences with regards to their overall genomic distribution of CO events (×^2^ = 7602.49, df = 4, *p*-value < 0.0001) driven by an enrichment of COs in intergenic regions in females and in gene-associated regions (downstream, exonic, intronic, and upstream) in males. Providing further evidence of a functional PRDM9, COs were harbored within CpG-islands no more frequently than expected by chance (Supplementary Figure 31) and COs were depleted around genes (Supplementary Figure 32), as expected given that PRDM9 tends to prevent meiotic recombination in close proximity to transcription start sites (Myers et al. 2005; Coop et al. 2008).

**Figure 1.**
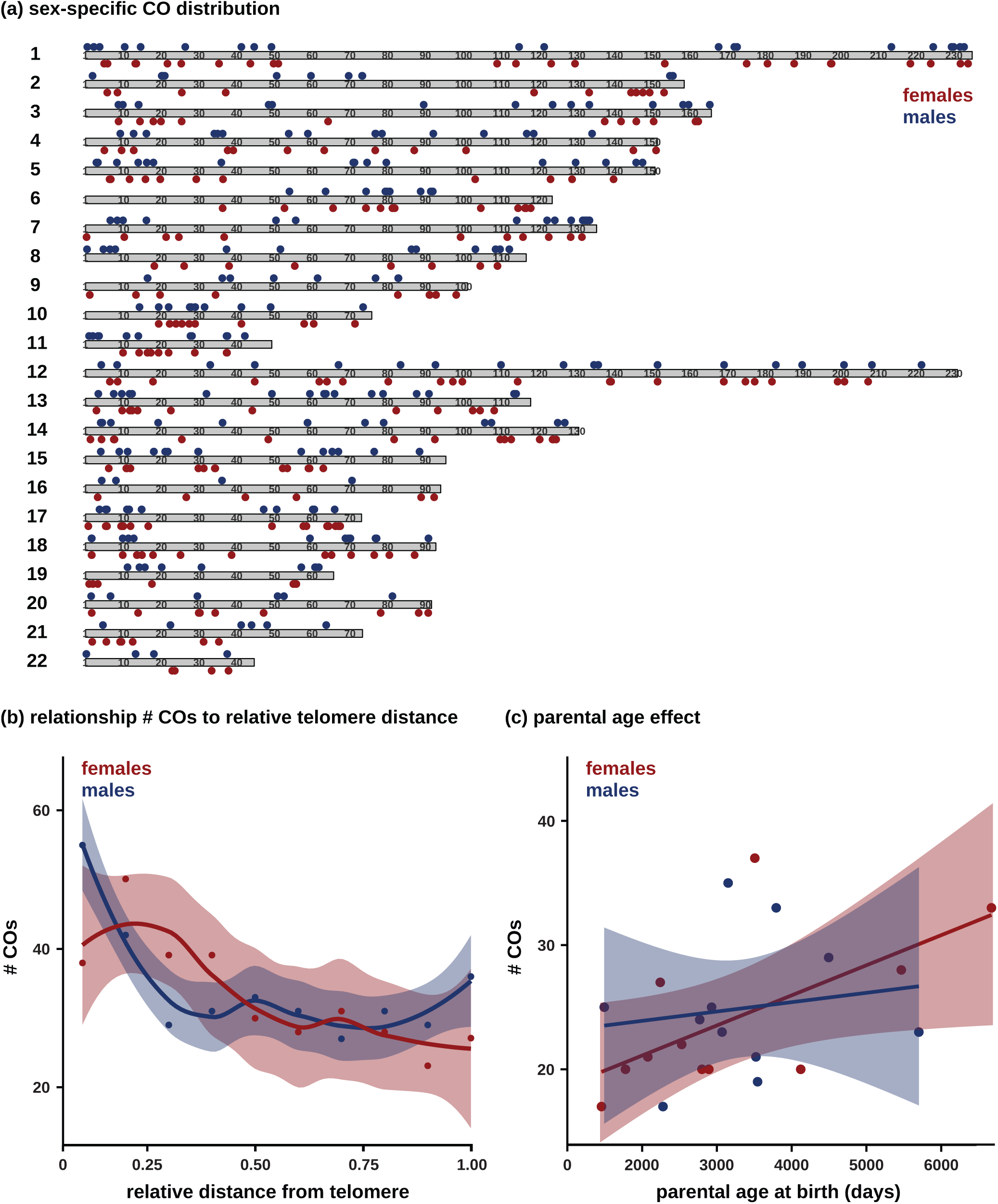
CO recombination in coppery titi monkeys. (a) Sex-specific CO distribution per autosome (chromosomes 1–22), with maternally-inherited COs shown in red and paternally-inherited COs shown in blue. (b) Relationship between the sex-specific number of CO events and relative distance to the nearest telomere. (c) Relationship between the sex-specific number of CO events and parental age at birth.

On average, 23 COs were observed per meiosis (Supplementary Table 6), with the average number of COs per chromosome significantly correlated with autosomal length (*r* = 0.78; *p*-value = 1.88 × 10^-5^; Supplementary Figure 33). Most primates studied to date exhibit higher recombination rates in females (see the review of Sardell and Kirkpatrick 2020); however, a similar number of COs were observed in both sexes in the three coppery titi monkey families, with an average of 1.10 COs and 1.13 COs per autosome per meiosis in the 11 maternal and 11 paternal meioses, respectively (two-tailed binomial test: *p*-value = 0.698; Supplementary Table 6), consistent with one obligatory CO per chromosome arm (Pardo-Manuel de Villena and Sapienza 2001). Due to these similarities in genome-wide CO rates (on average 1.00 cM/Mb and 1.04 cM/Mb in females and males, respectively), sex-specific autosomal genetic maps were also alike (2,409 cM vs 2,491 cM; Table 1; and see Supplementary Figure 34 for the cumulative genetic distance per chromosome in males and females). The estimated sex-averaged autosomal genetic map in coppery titi monkeys (2450 cM) was observed to be about 30% shorter than the sex-averaged autosomal genetic map in humans (Kong et al. 2002), despite the same karyotype (2*n* = 46; Dumas et al. 2005). Although the map length reported here is likely an underestimate due to the small sample size of this study (relative to existing studies in humans), it is noteworthy that similar differences from humans have been observed in other non-human primates (Rogers et al. 2000, 2006; Cox et al. 2006; Jasinska et al. 2007; Wall et al. 2022; Versoza et al. 2024; Versoza, Lloret-Villas et al. 2025; and see the review of Coop and Przeworski 2007).

**Table 1.** Sex-specific genetic linkage maps (male / female).

| **chromosome** | **length (Mb)** | **# CO events** | **cM** | **cM/Mb** |
| --- | --- | --- | --- | --- |
| 1 | 235.10 | 23 / 24 | 209.1 / 218.2 | 0.89 / 0.93 |
| 2 | 158.74 | 13 / 11 | 118.2 / 100.0 | 0.74 / 0.63 |
| 3 | 165.94 | 15 / 12 | 136.4 / 136.4 | 0.82 / 0.66 |
| 4 | 151.62 | 16 / 15 | 145.5 / 136.4 | 0.96 / 0.90 |
| 5 | 150.99 | 16 / 11 | 145.5 / 100.0 | 0.96 / 0.66 |
| 6 | 123.74 | 9 / 14 | 81.8 / 127.3 | 0.66 / 1.03 |
| 7 | 135.51 | 14 / 11 | 127.3 / 100.0 | 0.94 / 0.74 |
| 8 | 116.83 | 12 / 8 | 109.1 / 72.7 | 0.93 / 0.62 |
| 9 | 101.23 | 9 / 10 | 81.8 / 90.9 | 0.81 / 0.90 |
| 10 | 75.88 | 13 / 10 | 118.2 / 90.9 | 1.56 / 1.20 |
| 11 | 49.37 | 13 / 10 | 118.2 / 90.9 | 2.39 / 1.84 |
| 12 | 231.18 | 19 / 24 | 172.7 / 218.2 | 0.75 / 0.94 |
| 13 | 118.03 | 20 / 12 | 181.8 / 109.1 | 1.54 / 0.92 |
| 14 | 130.78 | 13 / 15 | 118.2 / 136.4 | 0.90 / 1.04 |
| 15 | 95.53 | 16 / 13 | 145.5 / 118.2 | 1.52 / 1.24 |
| 16 | 94.17 | 4 / 6 | 36.4 / 54.5 | 0.39 / 0.58 |
| 17 | 73.18 | 12 / 17 | 109.1 / 154.5 | 1.49 / 2.11 |
| 18 | 92.88 | 12 / 15 | 109.1 / 136.4 | 1.17 / 1.47 |
| 19 | 65.77 | 9 / 7 | 81.8 / 63.6 | 1.24 / 0.97 |
| 20 | 91.75 | 6 / 9 | 54.5 / 81.8 | 0.59 / 0.89 |
| 21 | 73.43 | 6 / 7 | 54.5 / 63.6 | 0.74 / 0.87 |
| 22 | 44.72 | 4 / 4 | 36.4 / 36.4 | 0.81 / 0.81 |
| **autosomal**<br>**(sex-averaged)** | **2,576.39** | **274 / 265**<br>**(269.5)** | **2,491 / 2,409**<br>**(2,450)** | **1.04 / 1.00**<br>**(1.02)** |

Notably, the number of CO events in females was significantly positively correlated with maternal age (*r* = 0.62; *p*-value = 0.043; Figure 1c), with an estimated 0.7 additional COs per year. A maternal age effect on recombination has also previously been observed in humans (Kong et al. 2004; Coop et al. 2008; Martin et al. 2015; Halldorsson et al. 2019; though see Hussin et al. 2011 and Porubsky et al. 2025) and has been suggested to be potentially driven by selection as a larger number of COs along homologous chromosomes may increase the stability of the bivalent (Robinson et al. 1998) and thus serve to counteract maternal age-related nondisjunction which frequently leads to aneuploidy and fetal loss (see the review of Hassold et al. 2000). In further concordance with humans, no statistically significant paternal age effect was observed for coppery titi monkeys (*r* = 0.15; *p*-value = 0.657).

### Identification of a putative PRDM9 sequence and binding motifs

In primates, as in many other sexually reproducing organisms, recombination hotspots are generally localized by the histone methyltransferase PRDM9 (Baudat et al. 2010; Myers et al. 2010; Parvanov et al. 2010). Although genes in the coppery titi monkey genome were previously characterized (Pfeifer et al. 2024), no annotation was available for PRDM9. As the highly repetitive nature and rapid evolution of PRDM9 makes the gene prone to mis-assembly, previously generated long-read data (Pfeifer et al. 2024) was thus used together with information from the human PRDM9 sequence to *de novo* assemble a putative PRDM9 sequence in coppery titi monkeys. A BLASTX (Altschul et al. 1990) search provided support for a high sequence similarity of the reassembled contig with PRDM9 sequences of other primates, including the central SSXRD and PR/SET domains as well as three C-terminal zinc fingers (Supplementary Figure 5); however, no N-terminal KRAB domain could be identified. The KRAB domain is an integral component of PRDM9 (Hohenauer and Moore 2012), necessary for double-strand repair and synapsis during meiosis, with previous research demonstrating that a domain truncation leads to a loss of PRDM9 function (Imai et al. 2017). Although certain strepsirrhines appear to be lacking the KRAB domain (Baker et al. 2017), given that the observed CO patterns are in general agreement with PRDM9-mediated recombination, the contig thus likely represents a partial / truncated PRDM9 ortholog, with the lack of a KRAB domain reflecting either an incomplete assembly or difficulties in the domain recognition / gene prediction due to its rapid evolution (Imbeault et al. 2017). Additional support for a functional PRDM9 was provided by gene expression observed in a previously collected testis sample, but not in the samples of heart and adrenal glands.

Analyses of high-resolution CO breakpoints identified a putative 15-mer PRDM9 binding motif, CCTGCCTCAGCCTCC, which shows considerable similarities to the degenerate 13(-15)-mer motif of the common PRDM9 A allele in humans, (N)CCNCCNTNNCCNC(N) (Myers et al. 2008; Baudat et al. 2010; Pratto et al. 2014; Patel et al. 2016; Altemose et al. 2017), including the well-defined spacing of multiple CC dinucleotides and generally high GC-content (Supplementary Figure 35). Similar to humans, in which the degenerate 13-mer motif is predicted to recruit COs to ∼40% of hotspots (Myers et al. 2008), the 15-mer motif identified in coppery titi monkeys is harbored within 43% of the CO breakpoint regions — a significant enrichment compared to the genomic background (Fisher’s exact test, *p*-value = 1.10 × 10^-35^).

### Sex-specific landscape of NCO recombination in coppery titi monkeys

A total of 287 putative autosomal NCO events were detected in the 22 meioses of the three coppery titi monkey families (Supplementary Figure 1). Out of these events, 103 (35.9%) were removed as they failed at least one of the stringent filter criteria applied to circumvent genotyping errors. Specifically, 7 events were removed due to low-confidence read mappings (with 5 and 2 events based on read mappings with an average depth of coverage of ≤ 20 and an average mapping quality of ≤ 40 across individuals, respectively), 4 events were removed as they were located within 10 phase-informative markers from the end of a chromosome, 58 events were removed as they were harbored within regions characterized by frequent haplotype changes (with < 10 markers showing a consistent haplotype phase on either side of the event), 29 events were removed because they coincided with another putative NCO in the surrounding 1 kb region, 3 events were removed due to a change of phase within the same region in another meiosis, and 2 events were removed because they overlapped with structural variation previously identified in the population (Versoza et al. 2026).

Of the final 184 NCOs (Supplementary Table 8), the vast majority (163, or 88.6%) were supported by a single marker, thus resulting in minimal conversion tract lengths of 1 bp, in agreement with previous observations in humans (92.8%; Schweiger et al. 2024). Among the 21 NCOs supported by more than one marker, 17 exhibited a mean minimal NCO conversion tract length of 55 bp (range: 2–350 bp), similar to those previously observed in other primates (Wall et al. 2022; Versoza et al. 2024; Charmouh et al. 2025; Porsborg et al. 2025; Versoza, Lloret-Villas et al. 2025). The remaining 4 NCOs exhibited minimal conversion tract lengths > 10 kb, with the two largest events being particularly poorly resolved. Previous research has demonstrated that such NCOs detected from short-read data frequently represent technical artefacts (Wall et al. 2022), thus the analyses presented here focused on the 180 NCO events with a tract length shorter than 10 kb.

The genome-wide distribution of the final 97 maternally-inherited and 83 paternally-inherited NCOs is depicted in Figure 2a (genomic locations are provided in Supplementary Table 8; and see Supplementary Figures 9-30 for the NCOs observed on each chromosome in each meiosis). In agreement with humans, where NCOs — like COs — are often concentrated at PRDM9-mediated hotspots (Williams et al. 2015; Halldorsson et al. 2016), the majority (52.8%) of short (10 kb) regions centered around NCOs contained the putative 15-mer PRDM9 binding motif. Analogous to COs, NCOs were localized in intergenic and, to a lesser extent, intronic regions (Supplementary Figure 30) but with a more similar overall genomic distribution between the sexes (×^2^ = 6.8332, df = 3, *p*-value = 0.077). Sex-specific differences exist, however, with regards to the minimal conversion tract lengths of extended events which, consistent with previous observations in humans (Palsson et al. 2025), were considerably longer in maternally-inherited NCOs than in paternally-inherited NCOs (72.8 bp vs 30.7 bp; Supplementary Table 8). Independent of sex, the overall distribution of minimal conversion tract lengths (Figure 2b) mimics those observed in other species (e.g., Williams et al. 2015; Halldorsson et al. 2016; Wall et al. 2022; Versoza et al. 2024; Palsson et al. 2025; Versoza, Lloret-Villas et al. 2025). Notably though, in contrast to the GC-biased gene conversion observed in these species (see the review of Duret and Galtier 2009), no transmission bias toward strong alleles (C or G) was detected in the dataset (45.1%).

**Figure 2.**
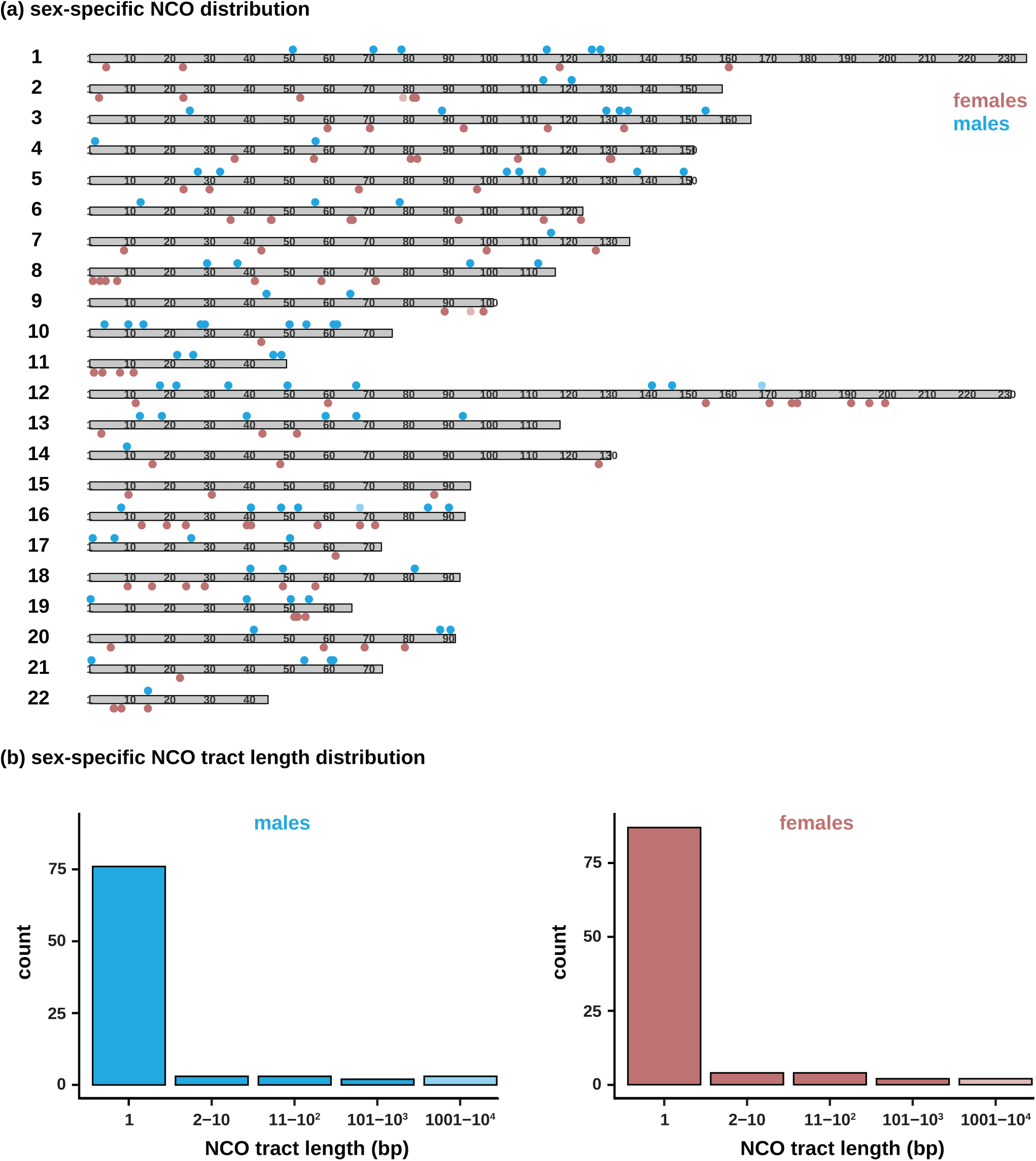
NCO recombination in coppery titi monkeys. (a) Sex-specific NCO distribution per autosome (chromosomes 1–22), with maternally-inherited NCOs shown in red and paternally-inherited NCOs shown in blue (NCO events > 10 kb are indicated by shading). (b) Sex-specific minimal NCO conversion tract length distributions.

Per meiosis, between 0 and 16 NCOs were identified, with an average of 8.8 and 7.6 NCOs in females and males, respectively, resulting in a combined sex-averaged rate of 0.37 NCOs per autosome (Supplementary Table 7). This estimate is considerably lower than the average number of NCOs observed in both humans (∼2–3 NCOs per chromosome and meiosis; Palsson et al. 2025) and rhesus macaques (∼1 NCO per chromosome and meiosis; Versoza et al. 2024) and likely represents an underestimate due to both the stringent filtering criteria applied and the status of the coppery titi monkey genome assembly compared to the near-complete telomere-to-telomere assemblies available for these biomedically-relevant species. For comparison, a recent study in aye-ayes, which relied on an assembly with a similar contiguity to that of coppery titi monkeys (930 contigs with a contig N50 of 80.4 Mb in aye-ayes [Versoza and Pfeifer 2024] vs 2,013 contigs with a contig N50 of 17 Mb in coppery titi monkeys [Pfeifer et al. 2024]), observed only a slightly higher rate of NCOs (∼0.7 NCOs per chromosome and meiosis; Versoza, Lloret-Villas et al. 2025). Therefore, in order to estimate the power to detect NCO events present in this study, 2,000 NCO events were simulated based on the tract length distributions observed in humans (Palsson et al. 2025). In concordance with recent work demonstrating a low power to detect short conversion tracts (∼2–4% in humans; Palsson et al. 2025), only 7.3% of short tracts were identifiable in coppery titi monkeys. Power was considerably higher (90.1%) for extended conversion tracts due to the high marker density; nevertheless, as most NCO events tend to be short, it is expected that the genuine number of bases affected by NCOs will be much larger.

Taken together, in addition to providing the first insights into the sex-specific landscapes of CO and NCO recombination in coppery titi monkeys, the resources generated in this study will also enable future evolutionary studies, including those aiming to understand the selective pressures shaping the genome of this important socially monogamous, pair-bonded behavioral primate model.

## ACKNOWLEDGEMENTS

DNA extraction, library preparation, and Illumina sequencing were conducted at the DNA Technologies and Expression Analysis Core at the UC Davis Genome Center (supported by NIH Shared Instrumentation Grant 1S10OD010786-01) and Novogene (Sacramento, CA, USA). Computations were performed on the Sol supercomputer at Arizona State University (Jennewein et al. 2023).

## FUNDING

This work was supported by the National Institute of General Medical Sciences of the National Institutes of Health under Award Number R35GM151008 to SPP and the California National Primate Research Center Pilot Program (NIH P51OD011107). CJV was supported by the National Science Foundation CAREER Award DEB-2045343 to SPP. KLB was supported by the Eunice Kennedy Shriver National Institute of Child Health and Human Development and the National Institute of Mental Health of the National Institutes of Health under Award Numbers R01HD092055 and MH125411, and by the Good Nature Institute. JDJ was supported by National Institutes of Health Award Number R35GM139383. The content is solely the responsibility of the authors and does not necessarily represent the official views of the funders.

## CONFLICT OF INTEREST

None declared.

## Notes

### Competing Interest Statement

The authors have declared no competing interest.

