## Supplementary Materials for "Sex-specific landscapes of crossover and non-crossover recombination in coppery titi monkeys (*Plecturocebus cupreus*)"

**Supplementary Table 1**. Sample information of the 17 individuals included in this study.

| ID | sex | coverage |
|---|---|---|
| 1 | M | 56.7 |
| 2 | F | 59.2 |
| 3 | M | 62.8 |
| 4 | M | 61.8 |
| 5 | M | 48.9 |
| 6 | F | 47.2 |
| 7 | M | 39.6 |
| 8 | F | 42.2 |
| 9 | F | 42.3 |
| 10 | M | 43.6 |
| 11 | F | 41.0 |
| 12 | F | 41.1 |
| 13 | M | 52.0 |
| 14 | F | 57.7 |
| 15 | M | 61.8 |
| 16 | M | 55.7 |
| 17 | M | 50.1 |

**Supplementary Table 2**. The variant dataset of this study contains 9,295,782 biallelic, single nucleotide polymorphisms (SNPs) with a transition-transversion ratio (Ts/Tv) of 2.62 in the autosomal genome.

| chromosome | length | # SNPs | Ts/Tv |
|---|---|---|---|
| 1 | 235,103,535 | 895,295 | 2.63 |
| 2 | 158,738,102 | 585,343 | 2.69 |
| 3 | 165,937,890 | 627,363 | 2.56 |
| 4 | 151,623,469 | 521,090 | 2.64 |
| 5 | 150,993,727 | 574,223 | 2.59 |
| 6 | 123,740,199 | 404,240 | 2.70 |
| 7 | 135,512,294 | 495,610 | 2.53 |
| 8 | 116,834,941 | 442,439 | 2.47 |
| 9 | 101,234,152 | 355,436 | 2.57 |
| 10 | 75,878,286 | 257,697 | 2.81 |
| 11 | 49,373,254 | 184,101 | 2.66 |
| 12 | 231,176,762 | 803,489 | 2.63 |
| 13 | 118,029,634 | 422,640 | 2.75 |
| 14 | 130,776,705 | 432,099 | 2.67 |
| 15 | 95,533,141 | 314,100 | 2.58 |
| 16 | 94,172,006 | 338,357 | 2.57 |
| 17 | 73,180,778 | 268,760 | 2.82 |
| 18 | 92,880,398 | 373,625 | 2.53 |
| 19 | 65,768,754 | 267,352 | 2.91 |
| 20 | 91,752,637 | 323,248 | 2.44 |
| 21 | 73,432,007 | 257,062 | 2.54 |
| 22 | 44,717,611 | 152,213 | 2.51 |
| ∑ or ∅ | **2,576,390,282** | **9,295,782** | **2.62** |

**Supplementary Table 3**. The number of phase-informative markers used in the detection of CO and NCO events in each family.

| **family**<br>*sire – dam (# offspring)* | **# phase-informative markers** | |
|---|---|---|
| | **maternally** | **paternally** |
| 1 – 2 (4) | 2,196,378 | 2,021,681 |
| 7 – 8 (4) | 2,168,166 | 2,065,096 |
| 13 – 14 (3) | 2,187,802 | 2,058,592 |

**Supplementary Table 4**. The density of phase-informative markers per autosome in each family.

| | marker density in chromosome (per kb) | | | | | | | | | | |
|---|---|---|---|---|---|---|---|---|---|---|---|
| **family** *sire–dam* | **1** | **2** | **3** | **4** | **5** | **6** | **7** | **8** | **9** | **10** | **11** |
| 1–2 | 0.87 | 0.79 | 0.81 | 0.74 | 0.84 | 0.76 | 0.85 | 0.86 | 0.79 | 0.84 | 0.81 |
| 7–8 | 0.88 | 0.84 | 0.89 | 0.90 | 0.87 | 0.71 | 0.85 | 0.91 | 0.83 | 0.78 | 0.82 |
| 13–14 | 0.86 | 0.88 | 0.91 | 0.72 | 0.90 | 0.68 | 0.94 | 0.87 | 0.71 | 0.62 | 0.79 |

| | marker density in chromosome (per kb) | | | | | | | | | | |
|---|---|---|---|---|---|---|---|---|---|---|---|
| **family** *sire–dam* | **12** | **13** | **14** | **15** | **16** | **17** | **18** | **19** | **20** | **21** | **22** |
| 1–2 | 0.78 | 0.79 | 0.76 | 0.71 | 0.91 | 0.73 | 0.92 | 0.95 | 0.85 | 0.87 | 0.87 |
| 7–8 | 0.81 | 0.68 | 0.68 | 0.57 | 0.79 | 0.85 | 0.93 | 0.91 | 0.92 | 0.74 | 0.80 |
| 13–14 | 0.80 | 0.78 | 0.83 | 0.93 | 0.74 | 0.87 | 0.83 | 0.94 | 0.95 | 0.82 | 0.60 |

**Supplementary Table 5**. Putative CO regions that were removed due to a clustering of ≥ 3 COs within 10 Mb in a single meiosis. Individual chromosomes are indicated by alternating shading.

| chromosome | region |
|---|---|
| 3 | 163,812,646 – 165,403,268 |
| 4 | 81,373,142 – 83,554,401 |
| 5 | 133,835,955 – 137,340,230 |
| 9 | 82,924,225 – 88,649,457 |
| 9 | 91,118,637 – 92,357,314 |
| 9 | 95,496,589 – 98,179,980 |
| 11 | 37,373,127 – 47,889,873 |
| 14 | 120,354,543 – 124,829,118 |
| 21 | 22,592,890 – 24,218,548 |
| 22 | 21,243,581 – 29,330,567 |

**Supplementary Table 6**. Summary statistics of the observed CO and NCO events. NCOs with a tract length ≥ 10 kb are shown in brackets.

| meiosis | # CO events | median resolution (bp) | # of NCO events | median tract length (bp) |
|---|---|---|---|---|
| 1–3 | 17 | 30,254 | 0 (1) | 0 |
| 1–4 | 25 | 20,673 | 3 | 1 |
| 1–5 | 19 | 15,663 | 3 | 1 |
| 1–6 | 25 | 29,455 | 9 | 1 |
| 2–3 | 27 | 8,879 | 6 | 1 |
| 2–4 | 20 | 11,150 | 7 | 1 |
| 2–5 | 37 | 13,747 | 5 | 1 |
| 2–6 | 17 | 15,947 | 8 | 1 |
| 7–9 | 24 | 6,423 | 9 (1) | 1 |
| 7–10 | 23 | 5,391 | 9 | 1 |
| 7–11 | 21 | 4,457 | 16 | 1 |
| 7–12 | 33 | 4,207 | 9 | 1 |
| 8–9 | 20 | 3,252 | 13 | 1 |
| 8–10 | 21 | 3,554 | 6 | 1 |
| 8–11 | 22 | 3,678 | 16 | 1 |
| 8–12 | 20 | 6,316 | 4 | 1 |
| 13–15 | 35 | 7,878 | 8 | 1 |
| 13–16 | 29 | 12,300 | 11 | 1 |
| 13–17 | 23 | 12,406 | 6 | 1 |
| 14–15 | 20 | 10,477 | 13 (1) | 1 |
| 14–16 | 28 | 8,007 | 6 | 1 |
| 14–17 | 33 | 10,469 | 13 (1) | 1 |

**Supplementary Table 7**. CO events detected in the 22 meioses of this study. Individual meioses are indicated in alternating shading.

| meiosis | chromosome | start | end | resolution |
|---|---|---|---|---|
| 1–3 | 1 | 10,359,005 | 10,397,928 | 38,923 |
| 1–3 | 1 | 232,929,261 | 232,941,488 | 12,227 |
| 1–3 | 2 | 20,501,366 | 20,537,906 | 36,540 |
| 1–3 | 2 | 155,571,482 | 155,574,951 | 3,469 |
| 1–3 | 7 | 131,829,699 | 131,839,207 | 9,508 |
| 1–3 | 7 | 132,404,304 | 132,504,407 | 100,103 |
| 1–3 | 9 | 49,885,693 | 49,921,328 | 35,635 |
| 1–3 | 10 | 19,399,165 | 19,404,296 | 5,131 |
| 1–3 | 12 | 4,157,251 | 4,187,505 | 30,254 |
| 1–3 | 12 | 126,713,462 | 126,755,545 | 42,083 |
| 1–3 | 12 | 151,663,900 | 151,671,029 | 7,129 |
| 1–3 | 13 | 65,937,523 | 66,097,472 | 159,949 |
| 1–3 | 16 | 8,051,311 | 8,059,158 | 7,847 |
| 1–3 | 16 | 70,598,416 | 70,689,919 | 91,503 |
| 1–3 | 17 | 47,220,076 | 47,233,276 | 13,200 |
| 1–3 | 18 | 76,850,695 | 76,912,066 | 61,371 |
| 1–3 | 21 | 22,493,163 | 22,516,550 | 23,387 |
| 1–4 | 1 | 439,415 | 493,056 | 53,641 |
| 1–4 | 1 | 49,278,217 | 49,303,131 | 24,914 |
| 1–4 | 1 | 114,936,547 | 114,967,079 | 30,532 |
| 1–4 | 1 | 213,655,444 | 213,670,753 | 15,309 |
| 1–4 | 3 | 123,813,705 | 123,868,069 | 54,364 |
| 1–4 | 3 | 159,875,190 | 159,915,853 | 40,663 |
| 1–4 | 5 | 74,628,834 | 74,652,093 | 23,259 |
| 1–4 | 6 | 79,951,706 | 79,958,180 | 6,474 |
| 1–4 | 8 | 7,872,085 | 7,891,476 | 19,391 |
| 1–4 | 8 | 103,356,623 | 103,363,039 | 6,416 |
| 1–4 | 10 | 27,617,238 | 27,627,771 | 10,533 |
| 1–4 | 11 | 10,853,392 | 10,910,175 | 56,783 |
| 1–4 | 13 | 9,490,151 | 9,712,228 | 222,077 |
| 1–4 | 13 | 11,650,795 | 11,671,468 | 20,673 |
| 1–4 | 13 | 75,844,979 | 75,880,240 | 35,261 |
| 1–4 | 14 | 4,023,110 | 4,032,738 | 9,628 |
| 1–4 | 14 | 105,830,025 | 105,872,121 | 42,096 |
| 1–4 | 15 | 8,919,093 | 8,942,803 | 23,710 |
| 1–4 | 15 | 65,404,622 | 65,407,041 | 2,419 |
| 1–4 | 17 | 10,959,066 | 10,975,912 | 16,846 |
| 1–4 | 17 | 50,668,268 | 50,679,971 | 11,703 |
| 1–4 | 19 | 15,729,023 | 15,731,028 | 2,005 |
| 1–4 | 19 | 20,184,099 | 20,195,931 | 11,832 |
| 1–4 | 20 | 29,650,748 | 29,652,178 | 1,430 |
| 1–4 | 21 | 63,785,801 | 63,832,464 | 46,663 |
| 1–5 | 2 | 20,192,952 | 20,206,547 | 13,595 |
| 1–5 | 3 | 150,413,547 | 150,414,090 | 543 |
| 1–5 | 4 | 58,872,800 | 59,008,392 | 135,592 |
| 1–5 | 4 | 92,200,639 | 92,209,835 | 9,196 |
| 1–5 | 5 | 35,993,486 | 36,009,382 | 15,896 |
| 1–5 | 7 | 16,100,797 | 16,165,555 | 64,758 |
| 1–5 | 8 | 37,371,067 | 37,376,821 | 5,754 |
| 1–5 | 9 | 82,787,359 | 83,006,721 | 219,362 |
| 1–5 | 10 | 28,050,149 | 28,054,346 | 4,197 |
| 1–5 | 10 | 41,319,555 | 41,338,138 | 18,583 |
| 1–5 | 10 | 73,581,797 | 73,597,460 | 15,663 |
| 1–5 | 11 | 939,839 | 940,467 | 628 |
| 1–5 | 12 | 8,356,222 | 8,372,500 | 16,278 |

| | | | | |
|---|---|---|---|---|
| 1–5 | 13 | 59,454,671 | 59,511,050 | 56,379 |
| 1–5 | 13 | 113,957,975 | 113,969,382 | 11,407 |
| 1–5 | 15 | 63,016,523 | 63,040,868 | 24,345 |
| 1–5 | 19 | 14,267,139 | 14,286,604 | 19,465 |
| 1–5 | 19 | 61,074,268 | 61,077,545 | 3,277 |
| 1–5 | 20 | 81,359,498 | 81,364,476 | 4,978 |
| 1–6 | 1 | 2,180,315 | 2,190,725 | 10,410 |
| 1–6 | 2 | 20,998,867 | 21,028,322 | 29,455 |
| 1–6 | 2 | 73,358,781 | 73,436,059 | 77,278 |
| 1–6 | 3 | 49,390,055 | 49,558,935 | 168,880 |
| 1–6 | 4 | 78,597,935 | 78,598,921 | 986 |
| 1–6 | 5 | 17,965,049 | 18,022,125 | 57,076 |
| 1–6 | 5 | 70,988,772 | 70,999,604 | 10,832 |
| 1–6 | 5 | 137,808,982 | 138,156,315 | 347,333 |
| 1–6 | 6 | 91,985,164 | 92,155,188 | 170,024 |
| 1–6 | 7 | 50,483,513 | 50,497,924 | 14,411 |
| 1–6 | 8 | 385,718 | 385,939 | 221 |
| 1–6 | 8 | 87,667,571 | 87,669,725 | 2,154 |
| 1–6 | 10 | 28,999,610 | 29,094,577 | 94,967 |
| 1–6 | 11 | 1,913,264 | 1,926,411 | 13,147 |
| 1–6 | 12 | 110,132,740 | 110,187,566 | 54,826 |
| 1–6 | 12 | 208,500,370 | 208,513,926 | 13,556 |
| 1–6 | 13 | 3,309,543 | 3,458,460 | 148,917 |
| 1–6 | 13 | 12,235,800 | 12,428,561 | 192,761 |
| 1–6 | 13 | 63,715,018 | 63,740,001 | 24,983 |
| 1–6 | 14 | 19,225,876 | 19,260,262 | 34,386 |
| 1–6 | 15 | 4,027,600 | 4,044,340 | 16,740 |
| 1–6 | 15 | 13,785,090 | 29,849,404 | 16,064,314 |
| 1–6 | 18 | 90,913,591 | 90,922,064 | 8,473 |
| 1–6 | 19 | 57,187,451 | 57,221,822 | 34,371 |
| 1–6 | 22 | 207,048 | 211,009 | 3,961 |
| 2–3 | 1 | 21,701,017 | 21,721,615 | 20,598 |
| 2–3 | 1 | 175,281,980 | 175,290,046 | 8,066 |
| 2–3 | 3 | 64,301,417 | 64,314,912 | 13,495 |
| 2–3 | 3 | 150,629,753 | 150,633,701 | 3,948 |
| 2–3 | 5 | 15,880,139 | 15,884,922 | 4,783 |
| 2–3 | 6 | 81,358,135 | 81,378,955 | 20,820 |
| 2–3 | 6 | 116,504,302 | 116,513,181 | 8,879 |
| 2–3 | 8 | 18,221,714 | 18,242,517 | 20,803 |
| 2–3 | 9 | 92,882,973 | 92,883,958 | 985 |
| 2–3 | 11 | 16,403,431 | 16,429,005 | 25,574 |
| 2–3 | 12 | 63,986,627 | 64,006,213 | 19,586 |
| 2–3 | 12 | 199,404,335 | 199,412,580 | 8,245 |
| 2–3 | 13 | 2,885,523 | 2,887,144 | 1,621 |
| 2–3 | 13 | 12,246,630 | 12,378,939 | 132,309 |
| 2–3 | 13 | 104,614,962 | 104,616,033 | 1,071 |
| 2–3 | 14 | 7,480,322 | 7,503,237 | 22,915 |
| 2–3 | 14 | 109,942,786 | 109,943,769 | 983 |
| 2–3 | 16 | 25,691,354 | 27,721,676 | 2,030,322 |
| 2–3 | 17 | 10,021,899 | 10,023,926 | 2,027 |
| 2–3 | 17 | 67,516,064 | 67,530,715 | 14,651 |
| 2–3 | 18 | 14,937,066 | 14,976,530 | 39,464 |
| 2–3 | 18 | 63,498,382 | 63,541,571 | 43,189 |
| 2–3 | 19 | 17,603,789 | 17,603,898 | 109 |
| 2–3 | 19 | 55,880,980 | 55,882,982 | 2,002 |
| 2–3 | 20 | 30,310,736 | 30,348,967 | 38,231 |
| 2–3 | 21 | 1,765,730 | 1,767,072 | 1,342 |
| 2–3 | 22 | 33,426,424 | 33,434,069 | 7,645 |
| 2–4 | 1 | 123,455,758 | 123,458,999 | 3,241 |

| | | | | |
|---|---|---|---|---|
| 2–4 | 1 | 197,780,583 | 197,832,730 | 52,147 |
| 2–4 | 3 | 162,448,631 | 162,453,509 | 4,878 |
| 2–4 | 5 | 6,367,221 | 6,370,924 | 3,703 |
| 2–4 | 6 | 74,358,755 | 74,389,173 | 30,418 |
| 2–4 | 6 | 116,778,269 | 116,798,056 | 19,787 |
| 2–4 | 7 | 10,265,245 | 10,271,402 | 6,157 |
| 2–4 | 8 | 38,029,813 | 38,082,713 | 52,900 |
| 2–4 | 10 | 19,394,298 | 19,398,323 | 4,025 |
| 2–4 | 10 | 71,432,120 | 71,449,964 | 17,844 |
| 2–4 | 11 | 28,966,144 | 28,968,766 | 2,622 |
| 2–4 | 12 | 94,153,721 | 94,157,297 | 3,576 |
| 2–4 | 13 | 13,788,365 | 13,798,468 | 10,103 |
| 2–4 | 15 | 11,894,886 | 11,899,868 | 4,982 |
| 2–4 | 15 | 29,818,791 | 29,997,419 | 178,628 |
| 2–4 | 15 | 34,389,178 | 34,401,375 | 12,197 |
| 2–4 | 17 | 64,415,192 | 64,504,286 | 89,094 |
| 2–4 | 18 | 17,840,832 | 17,924,950 | 84,118 |
| 2–4 | 18 | 65,219,089 | 65,228,948 | 9,859 |
| 2–4 | 20 | 34,368,157 | 34,381,806 | 13,649 |
| 2–5 | 1 | 35,382,890 | 35,395,156 | 12,266 |
| 2–5 | 1 | 114,105,442 | 114,126,950 | 21,508 |
| 2–5 | 1 | 218,693,960 | 218,711,645 | 17,685 |
| 2–5 | 1 | 233,989,020 | 233,999,063 | 10,043 |
| 2–5 | 2 | 5,726,814 | 5,780,675 | 53,861 |
| 2–5 | 2 | 118,921,676 | 118,929,860 | 8,184 |
| 2–5 | 4 | 4,984,529 | 4,991,845 | 7,316 |
| 2–5 | 4 | 12,746,324 | 12,786,525 | 40,201 |
| 2–5 | 4 | 76,802,483 | 76,826,323 | 23,840 |
| 2–5 | 4 | 145,232,855 | 145,262,717 | 29,862 |
| 2–5 | 4 | 151,204,445 | 151,331,419 | 126,974 |
| 2–5 | 5 | 36,444,639 | 36,446,514 | 1,875 |
| 2–5 | 7 | 36,751,031 | 36,754,148 | 3,117 |
| 2–5 | 7 | 115,970,691 | 115,998,546 | 27,855 |
| 2–5 | 8 | 55,453,054 | 55,492,412 | 39,358 |
| 2–5 | 8 | 109,206,556 | 109,230,669 | 24,113 |
| 2–5 | 9 | 91,241,898 | 91,248,755 | 6,857 |
| 2–5 | 10 | 27,468,962 | 27,477,554 | 8,592 |
| 2–5 | 10 | 60,491,856 | 60,496,415 | 4,559 |
| 2–5 | 11 | 17,309,215 | 17,312,454 | 3,239 |
| 2–5 | 12 | 6,438,209 | 6,443,930 | 5,721 |
| 2–5 | 12 | 68,210,859 | 68,221,832 | 10,973 |
| 2–5 | 12 | 174,830,243 | 175,104,647 | 274,404 |
| 2–5 | 13 | 82,369,777 | 82,374,491 | 4,714 |
| 2–5 | 13 | 108,348,032 | 108,355,716 | 7,684 |
| 2–5 | 14 | 7,657,096 | 7,676,453 | 19,357 |
| 2–5 | 14 | 81,679,076 | 81,952,248 | 273,172 |
| 2–5 | 15 | 31,323,949 | 31,378,337 | 54,388 |
| 2–5 | 15 | 34,266,571 | 34,277,595 | 11,024 |
| 2–5 | 17 | 16,623,729 | 16,626,292 | 2,563 |
| 2–5 | 17 | 58,538,369 | 58,591,722 | 53,353 |
| 2–5 | 19 | 936,770 | 963,237 | 26,467 |
| 2–5 | 19 | 55,097,009 | 55,100,540 | 3,531 |
| 2–5 | 20 | 47,209,207 | 47,222,954 | 13,747 |
| 2–5 | 22 | 22,982,293 | 22,986,919 | 4,626 |
| 2–5 | 22 | 23,611,789 | 23,703,229 | 91,440 |
| 2–5 | 22 | 37,806,646 | 37,991,671 | 185,025 |
| 2–6 | 1 | 49,833,355 | 49,889,385 | 56,030 |
| 2–6 | 2 | 153,385,824 | 153,395,309 | 9,485 |
| 2–6 | 3 | 17,964,420 | 17,984,387 | 19,967 |

| | | | | |
|---|---|---|---|---|
| 2–6 | 6 | 52,737,385 | 52,743,629 | 6,244 |
| 2–6 | 6 | 65,621,030 | 65,636,625 | 15,595 |
| 2–6 | 7 | 128,622,149 | 128,626,903 | 4,754 |
| 2–6 | 8 | 26,133,400 | 26,135,354 | 1,954 |
| 2–6 | 9 | 34,460,255 | 34,464,771 | 4,516 |
| 2–6 | 9 | 92,940,439 | 92,945,509 | 5,070 |
| 2–6 | 11 | 37,382,047 | 37,628,578 | 246,531 |
| 2–6 | 12 | 97,374,290 | 97,392,421 | 18,131 |
| 2–6 | 14 | 25,514,214 | 25,536,992 | 22,778 |
| 2–6 | 17 | 727,233 | 743,180 | 15,947 |
| 2–6 | 17 | 5,264,720 | 5,425,876 | 161,156 |
| 2–6 | 17 | 66,280,722 | 66,287,552 | 6,830 |
| 2–6 | 20 | 29,987,447 | 30,017,996 | 30,549 |
| 2–6 | 21 | 12,487,838 | 12,504,100 | 16,262 |
| 7–9 | 1 | 14,598,582 | 14,606,428 | 7,846 |
| 7–9 | 3 | 128,746,771 | 128,759,007 | 12,236 |
| 7–9 | 4 | 16,132,225 | 16,133,169 | 944 |
| 7–9 | 7 | 128,764,534 | 128,765,327 | 793 |
| 7–9 | 8 | 112,355,189 | 112,359,811 | 4,622 |
| 7–9 | 9 | 16,413,078 | 16,428,907 | 15,829 |
| 7–9 | 10 | 41,318,751 | 41,331,048 | 12,297 |
| 7–9 | 10 | 49,075,862 | 49,080,862 | 5,000 |
| 7–9 | 11 | 10,854,662 | 10,863,203 | 8,541 |
| 7–9 | 11 | 27,780,833 | 27,858,254 | 77,421 |
| 7–9 | 12 | 169,128,783 | 169,469,071 | 340,288 |
| 7–9 | 13 | 87,781,890 | 87,796,412 | 14,522 |
| 7–9 | 14 | 24,003,100 | 48,657,336 | 24,654,236 |
| 7–9 | 14 | 58,806,322 | 58,863,798 | 57,476 |
| 7–9 | 14 | 58,876,283 | 58,878,237 | 1,954 |
| 7–9 | 14 | 127,058,675 | 127,063,285 | 4,610 |
| 7–9 | 15 | 18,074,762 | 18,076,201 | 1,439 |
| 7–9 | 17 | 14,890,294 | 14,891,257 | 963 |
| 7–9 | 18 | 1,570,797 | 1,580,294 | 9,497 |
| 7–9 | 18 | 1,649,357 | 1,650,085 | 728 |
| 7–9 | 19 | 61,843,828 | 61,846,747 | 2,919 |
| 7–9 | 20 | 1,478,562 | 1,479,245 | 683 |
| 7–9 | 20 | 52,603,519 | 52,605,999 | 2,480 |
| 7–9 | 22 | 37,128,405 | 37,991,671 | 863,266 |
| 7–10 | 1 | 26,422,932 | 26,438,157 | 15,225 |
| 7–10 | 1 | 172,010,727 | 172,022,557 | 11,830 |
| 7–10 | 1 | 172,744,277 | 172,756,222 | 11,945 |
| 7–10 | 2 | 50,670,318 | 50,684,168 | 13,850 |
| 7–10 | 3 | 8,707,088 | 8,708,713 | 1,625 |
| 7–10 | 4 | 36,352,659 | 36,358,219 | 5,560 |
| 7–10 | 5 | 129,940,994 | 129,942,407 | 1,413 |
| 7–10 | 6 | 54,050,959 | 54,054,462 | 3,503 |
| 7–10 | 7 | 8,403,595 | 8,405,710 | 2,115 |
| 7–10 | 7 | 114,305,190 | 114,336,684 | 31,494 |
| 7–10 | 8 | 108,690,450 | 108,743,300 | 52,850 |
| 7–10 | 9 | 76,824,763 | 76,947,277 | 122,514 |
| 7–10 | 10 | 14,273,553 | 14,329,612 | 56,059 |
| 7–10 | 10 | 31,555,692 | 31,557,910 | 2,218 |
| 7–10 | 11 | 28,213,837 | 28,216,957 | 3,120 |
| 7–10 | 12 | 221,653,207 | 221,655,346 | 2,139 |
| 7–10 | 13 | 49,415,160 | 49,420,245 | 5,085 |
| 7–10 | 14 | 6,704,518 | 6,709,030 | 4,512 |
| 7–10 | 15 | 11,152,107 | 11,152,671 | 564 |
| 7–10 | 17 | 3,696,449 | 3,701,840 | 5,391 |
| 7–10 | 17 | 5,256,877 | 5,347,234 | 90,357 |

| | | | | |
|---|---|---|---|---|
| 7–10 | 18 | 68,902,318 | 68,912,485 | 10,167 |
| 7–10 | 20 | 50,919,295 | 50,922,899 | 3,604 |
| 7–11 | 1 | 3,730,160 | 3,731,185 | 1,025 |
| 7–11 | 1 | 231,866,018 | 231,872,355 | 6,337 |
| 7–11 | 2 | 154,935,228 | 154,938,682 | 3,454 |
| 7–11 | 3 | 14,095,106 | 14,098,324 | 3,218 |
| 7–11 | 3 | 133,577,827 | 133,583,856 | 6,029 |
| 7–11 | 4 | 53,866,398 | 53,867,034 | 636 |
| 7–11 | 5 | 3,279,653 | 3,282,459 | 2,806 |
| 7–11 | 6 | 74,363,623 | 74,388,744 | 25,121 |
| 7–11 | 6 | 91,521,774 | 91,540,079 | 18,305 |
| 7–11 | 7 | 133,000,643 | 133,005,407 | 4,764 |
| 7–11 | 8 | 4,729,265 | 4,734,791 | 5,526 |
| 7–11 | 9 | 16,460,858 | 16,463,275 | 2,417 |
| 7–11 | 12 | 189,989,835 | 190,016,804 | 26,969 |
| 7–11 | 13 | 91,055,028 | 91,058,018 | 2,990 |
| 7–11 | 14 | 74,100,306 | 74,105,156 | 4,850 |
| 7–11 | 14 | 125,196,519 | 125,197,269 | 750 |
| 7–11 | 15 | 21,108,077 | 21,116,292 | 8,215 |
| 7–11 | 16 | 36,179,458 | 36,182,748 | 3,290 |
| 7–11 | 17 | 60,247,450 | 60,248,994 | 1,544 |
| 7–11 | 21 | 4,589,234 | 4,630,053 | 40,819 |
| 7–11 | 21 | 48,141,904 | 48,146,361 | 4,457 |
| 7–12 | 1 | 2,195,714 | 2,197,336 | 1,622 |
| 7–12 | 1 | 41,304,574 | 41,315,743 | 11,169 |
| 7–12 | 1 | 224,759,096 | 224,760,372 | 1,276 |
| 7–12 | 2 | 1,852,458 | 1,853,655 | 1,197 |
| 7–12 | 2 | 69,765,818 | 69,772,125 | 6,307 |
| 7–12 | 2 | 155,688,929 | 155,689,854 | 925 |
| 7–12 | 3 | 9,888,624 | 9,893,649 | 5,025 |
| 7–12 | 3 | 89,653,827 | 89,656,213 | 2,386 |
| 7–12 | 3 | 165,515,839 | 165,516,848 | 1,009 |
| 7–12 | 4 | 9,208,315 | 9,216,328 | 8,013 |
| 7–12 | 4 | 12,701,365 | 12,781,730 | 80,365 |
| 7–12 | 5 | 16,227,746 | 16,229,462 | 1,716 |
| 7–12 | 6 | 63,640,536 | 63,643,164 | 2,628 |
| 7–12 | 7 | 6,555,049 | 6,561,149 | 6,100 |
| 7–12 | 7 | 55,718,330 | 55,722,290 | 3,960 |
| 7–12 | 8 | 51,654,131 | 51,684,200 | 30,069 |
| 7–12 | 9 | 36,247,248 | 36,249,050 | 1,802 |
| 7–12 | 10 | 22,019,895 | 22,034,518 | 14,623 |
| 7–12 | 10 | 29,074,919 | 29,094,305 | 19,386 |
| 7–12 | 11 | 3,310,300 | 3,311,202 | 902 |
| 7–12 | 12 | 183,028,055 | 183,042,529 | 14,474 |
| 7–12 | 13 | 63,636,689 | 63,638,685 | 1,996 |
| 7–12 | 13 | 113,600,196 | 113,606,259 | 6,063 |
| 7–12 | 14 | 79,038,994 | 79,040,145 | 1,151 |
| 7–12 | 15 | 4,031,292 | 4,035,499 | 4,207 |
| 7–12 | 15 | 29,818,791 | 29,862,160 | 43,369 |
| 7–12 | 15 | 76,533,478 | 76,536,031 | 2,553 |
| 7–12 | 16 | 4,280,295 | 4,284,619 | 4,324 |
| 7–12 | 18 | 9,854,222 | 9,861,871 | 7,649 |
| 7–12 | 18 | 12,772,209 | 12,773,356 | 1,147 |
| 7–12 | 18 | 69,735,583 | 69,742,279 | 6,696 |
| 7–12 | 19 | 30,725,421 | 30,730,891 | 5,470 |
| 7–12 | 21 | 41,263,111 | 41,264,785 | 1,674 |
| 8–9 | 1 | 5,825,873 | 5,831,601 | 5,728 |
| 8–9 | 2 | 37,156,076 | 37,160,735 | 4,659 |
| 8–9 | 3 | 141,976,430 | 141,980,676 | 4,246 |

| | | | | |
|---|---|---|---|---|
| 8–9 | 4 | 9,533,624 | 9,571,249 | 37,625 |
| 8–9 | 4 | 12,780,191 | 12,786,147 | 5,956 |
| 8–9 | 4 | 53,553,031 | 53,555,391 | 2,360 |
| 8–9 | 5 | 129,000,436 | 129,002,607 | 2,171 |
| 8–9 | 7 | 266,780 | 267,110 | 330 |
| 8–9 | 7 | 131,606,501 | 131,609,989 | 3,488 |
| 8–9 | 9 | 1,116,625 | 1,117,480 | 855 |
| 8–9 | 12 | 169,123,278 | 169,374,263 | 250,985 |
| 8–9 | 12 | 207,499,166 | 207,500,457 | 1,291 |
| 8–9 | 13 | 9,688,398 | 9,725,822 | 37,424 |
| 8–9 | 13 | 11,724,819 | 11,730,880 | 6,061 |
| 8–9 | 14 | 4,228,697 | 4,231,014 | 2,317 |
| 8–9 | 17 | 57,754,322 | 57,759,717 | 5,395 |
| 8–9 | 18 | 80,547,025 | 80,547,400 | 375 |
| 8–9 | 19 | 1,936,747 | 1,937,341 | 594 |
| 8–9 | 20 | 78,245,033 | 78,246,918 | 1,885 |
| 8–9 | 21 | 5,511,123 | 5,514,139 | 3,016 |
| 8–10 | 1 | 4,923,136 | 4,937,126 | 13,990 |
| 8–10 | 1 | 153,584,859 | 153,585,949 | 1,090 |
| 8–10 | 1 | 231,977,911 | 231,978,477 | 566 |
| 8–10 | 2 | 25,515,752 | 25,517,429 | 1,677 |
| 8–10 | 2 | 133,524,244 | 133,527,592 | 3,348 |
| 8–10 | 3 | 161,724,552 | 161,728,267 | 3,715 |
| 8–10 | 4 | 39,164,635 | 39,171,630 | 6,995 |
| 8–10 | 4 | 76,800,285 | 76,818,849 | 18,564 |
| 8–10 | 4 | 100,883,864 | 100,887,693 | 3,829 |
| 8–10 | 5 | 19,811,123 | 19,812,258 | 1,135 |
| 8–10 | 6 | 81,971,929 | 81,972,948 | 1,019 |
| 8–10 | 11 | 14,182,445 | 14,214,049 | 31,604 |
| 8–10 | 11 | 37,382,047 | 37,441,849 | 59,802 |
| 8–10 | 12 | 8,494,221 | 8,494,867 | 646 |
| 8–10 | 14 | 79,020,560 | 106,214,398 | 27,193,838 |
| 8–10 | 15 | 59,095,466 | 59,099,020 | 3,554 |
| 8–10 | 17 | 9,419,042 | 9,422,149 | 3,107 |
| 8–10 | 17 | 49,431,848 | 49,434,712 | 2,864 |
| 8–10 | 18 | 25,178,421 | 25,182,582 | 4,161 |
| 8–10 | 18 | 87,247,235 | 87,250,163 | 2,928 |
| 8–10 | 21 | 9,165,494 | 9,173,780 | 8,286 |
| 8–11 | 1 | 129,779,958 | 129,781,236 | 1,278 |
| 8–11 | 1 | 187,882,436 | 187,883,723 | 1,287 |
| 8–11 | 3 | 14,428,629 | 14,429,838 | 1,209 |
| 8–11 | 5 | 123,279,967 | 123,280,573 | 606 |
| 8–11 | 7 | 111,812,848 | 111,817,531 | 4,683 |
| 8–11 | 8 | 104,656,369 | 104,662,511 | 6,142 |
| 8–11 | 10 | 22,293,639 | 22,295,957 | 2,318 |
| 8–11 | 12 | 17,789,420 | 17,961,760 | 172,340 |
| 8–11 | 12 | 80,262,436 | 80,263,375 | 939 |
| 8–11 | 12 | 139,001,174 | 139,011,938 | 10,764 |
| 8–11 | 12 | 151,668,661 | 151,673,010 | 4,349 |
| 8–11 | 12 | 181,997,515 | 182,001,873 | 4,358 |
| 8–11 | 13 | 44,210,331 | 44,212,902 | 2,571 |
| 8–11 | 14 | 1,291,726 | 1,296,051 | 4,325 |
| 8–11 | 14 | 112,853,373 | 112,859,252 | 5,879 |
| 8–11 | 15 | 10,951,301 | 10,955,389 | 4,088 |
| 8–11 | 16 | 88,979,549 | 88,981,995 | 2,446 |
| 8–11 | 17 | 5,675,357 | 5,680,928 | 5,571 |
| 8–11 | 18 | 1,571,589 | 1,595,507 | 23,918 |
| 8–11 | 18 | 1,649,569 | 1,652,837 | 3,268 |
| 8–11 | 18 | 70,408,655 | 70,409,830 | 1,175 |

| | | | | |
|---|---|---|---|---|
| 8–11 | 20 | 1,630,843 | 1,631,112 | 269 |
| 8–12 | 1 | 13,452,458 | 13,454,768 | 2,310 |
| 8–12 | 1 | 109,124,982 | 109,127,009 | 2,027 |
| 8–12 | 1 | 224,091,047 | 224,094,317 | 3,270 |
| 8–12 | 2 | 149,569,200 | 149,578,511 | 9,311 |
| 8–12 | 3 | 8,743,388 | 8,759,606 | 16,218 |
| 8–12 | 5 | 6,684,905 | 6,689,283 | 4,378 |
| 8–12 | 6 | 104,783,587 | 104,818,264 | 34,677 |
| 8–12 | 6 | 114,569,551 | 114,789,593 | 220,042 |
| 8–12 | 6 | 118,015,325 | 118,055,461 | 40,136 |
| 8–12 | 7 | 21,356,059 | 21,358,521 | 2,462 |
| 8–12 | 8 | 91,831,289 | 91,842,369 | 11,080 |
| 8–12 | 9 | 13,359,038 | 13,369,346 | 10,308 |
| 8–12 | 10 | 23,971,224 | 23,972,941 | 1,717 |
| 8–12 | 11 | 9,942,102 | 9,946,255 | 4,153 |
| 8–12 | 13 | 12,242,913 | 32,990,154 | 20,747,241 |
| 8–12 | 13 | 82,292,703 | 104,491,498 | 22,198,795 |
| 8–12 | 15 | 52,276,624 | 52,284,192 | 7,568 |
| 8–12 | 17 | 5,519,962 | 5,520,926 | 964 |
| 8–12 | 18 | 38,718,697 | 38,721,506 | 2,809 |
| 8–12 | 21 | 9,520,740 | 9,525,804 | 5,064 |
| 13–15 | 1 | 44,715,721 | 44,719,104 | 3,383 |
| 13–15 | 1 | 167,846,919 | 167,860,081 | 13,162 |
| 13–15 | 1 | 230,163,749 | 230,167,910 | 4,161 |
| 13–15 | 3 | 48,539,843 | 48,544,987 | 5,144 |
| 13–15 | 3 | 113,984,962 | 114,003,493 | 18,531 |
| 13–15 | 3 | 158,408,478 | 158,432,027 | 23,549 |
| 13–15 | 4 | 35,012,483 | 35,018,623 | 6,140 |
| 13–15 | 4 | 76,946,979 | 76,953,113 | 6,134 |
| 13–15 | 4 | 116,944,944 | 117,011,241 | 66,297 |
| 13–15 | 4 | 118,809,003 | 118,824,978 | 15,975 |
| 13–15 | 4 | 134,228,480 | 134,377,682 | 149,202 |
| 13–15 | 5 | 13,956,807 | 13,960,870 | 4,063 |
| 13–15 | 5 | 79,744,786 | 79,751,019 | 6,233 |
| 13–15 | 6 | 79,498,659 | 79,502,435 | 3,776 |
| 13–15 | 7 | 133,609,582 | 133,616,218 | 6,636 |
| 13–15 | 9 | 61,519,712 | 61,533,789 | 14,077 |
| 13–15 | 11 | 13,993,706 | 14,000,289 | 6,583 |
| 13–15 | 11 | 37,348,325 | 37,423,923 | 75,598 |
| 13–15 | 12 | 44,863,517 | 44,878,682 | 15,165 |
| 13–15 | 12 | 92,771,999 | 92,773,511 | 1,512 |
| 13–15 | 12 | 134,812,531 | 134,819,674 | 7,143 |
| 13–15 | 13 | 32,053,200 | 32,061,078 | 7,878 |
| 13–15 | 13 | 114,157,640 | 114,169,606 | 11,966 |
| 13–15 | 14 | 4,437,007 | 4,438,958 | 1,951 |
| 13–15 | 14 | 107,514,876 | 107,736,882 | 222,006 |
| 13–15 | 15 | 29,857,111 | 29,994,760 | 137,649 |
| 13–15 | 15 | 57,178,417 | 57,188,094 | 9,677 |
| 13–15 | 15 | 66,993,745 | 67,095,102 | 101,357 |
| 13–15 | 15 | 88,551,030 | 88,556,034 | 5,004 |
| 13–15 | 17 | 14,846,570 | 14,849,415 | 2,845 |
| 13–15 | 17 | 60,658,181 | 60,664,575 | 6,394 |
| 13–15 | 18 | 59,500,239 | 59,500,446 | 207 |
| 13–15 | 19 | 11,124,720 | 11,136,737 | 12,017 |
| 13–15 | 19 | 60,798,733 | 60,882,771 | 84,038 |
| 13–15 | 21 | 43,998,020 | 44,030,936 | 32,916 |
| 13–16 | 1 | 121,601,087 | 121,602,193 | 1,106 |
| 13–16 | 1 | 229,606,390 | 229,618,690 | 12,300 |
| 13–16 | 2 | 73,302,295 | 73,329,810 | 27,515 |

| | | | | |
|---|---|---|---|---|
| 13–16 | 2 | 155,743,949 | 155,756,074 | 12,125 |
| 13–16 | 5 | 8,300,890 | 8,311,852 | 10,962 |
| 13–16 | 5 | 71,309,798 | 71,311,748 | 1,950 |
| 13–16 | 5 | 147,674,406 | 147,717,184 | 42,778 |
| 13–16 | 6 | 80,656,541 | 80,659,142 | 2,601 |
| 13–16 | 7 | 9,930,264 | 9,936,348 | 6,084 |
| 13–16 | 7 | 124,407,742 | 124,435,745 | 28,003 |
| 13–16 | 8 | 6,472,472 | 6,477,599 | 5,127 |
| 13–16 | 8 | 86,422,641 | 86,427,104 | 4,463 |
| 13–16 | 9 | 38,368,307 | 38,368,401 | 94 |
| 13–16 | 9 | 82,924,225 | 83,014,740 | 90,515 |
| 13–16 | 10 | 41,239,762 | 41,375,148 | 135,386 |
| 13–16 | 11 | 10,850,295 | 10,927,183 | 76,888 |
| 13–16 | 11 | 37,607,087 | 37,623,445 | 16,358 |
| 13–16 | 11 | 42,020,807 | 42,449,912 | 429,105 |
| 13–16 | 12 | 67,059,479 | 67,084,467 | 24,988 |
| 13–16 | 12 | 151,623,806 | 151,678,465 | 54,659 |
| 13–16 | 12 | 201,093,617 | 201,098,522 | 4,905 |
| 13–16 | 13 | 7,461,515 | 7,476,458 | 14,943 |
| 13–16 | 13 | 12,235,983 | 12,376,981 | 140,998 |
| 13–16 | 17 | 11,565,200 | 11,571,687 | 6,487 |
| 13–16 | 17 | 66,051,934 | 66,057,053 | 5,119 |
| 13–16 | 18 | 1,617,620 | 21,129,474 | 19,511,854 |
| 13–16 | 18 | 77,163,008 | 77,166,328 | 3,320 |
| 13–16 | 20 | 6,609,794 | 6,630,336 | 20,542 |
| 13–16 | 22 | 13,258,905 | 13,270,510 | 11,605 |
| 13–17 | 1 | 172,013,829 | 172,022,712 | 8,883 |
| 13–17 | 1 | 172,741,577 | 172,759,137 | 17,560 |
| 13–17 | 2 | 59,755,639 | 59,781,654 | 26,015 |
| 13–17 | 3 | 14,009,240 | 14,051,969 | 42,729 |
| 13–17 | 4 | 34,096,258 | 34,193,119 | 96,861 |
| 13–17 | 4 | 76,802,483 | 76,824,019 | 21,536 |
| 13–17 | 4 | 105,616,287 | 105,623,529 | 7,242 |
| 13–17 | 5 | 2,877,292 | 2,878,398 | 1,106 |
| 13–17 | 5 | 121,187,151 | 121,195,990 | 8,839 |
| 13–17 | 5 | 145,980,594 | 145,993,000 | 12,406 |
| 13–17 | 6 | 88,818,180 | 88,875,673 | 57,493 |
| 13–17 | 7 | 122,370,505 | 122,380,840 | 10,335 |
| 13–17 | 8 | 109,907,010 | 109,913,894 | 6,884 |
| 13–17 | 11 | 3,563,905 | 3,592,671 | 28,766 |
| 13–17 | 12 | 33,059,317 | 33,064,465 | 5,148 |
| 13–17 | 12 | 75,213,424 | 91,837,540 | 16,624,116 |
| 13–17 | 12 | 135,936,452 | 135,937,796 | 1,344 |
| 13–17 | 13 | 63,226,185 | 63,252,724 | 26,539 |
| 13–17 | 13 | 78,857,989 | 78,902,592 | 44,603 |
| 13–17 | 15 | 18,128,534 | 18,129,367 | 833 |
| 13–17 | 17 | 5,638,507 | 5,656,952 | 18,445 |
| 13–17 | 18 | 70,216,749 | 70,228,212 | 11,463 |
| 13–17 | 22 | 18,111,262 | 18,123,200 | 11,938 |
| 14–15 | 1 | 25,370,085 | 25,378,552 | 8,467 |
| 14–15 | 1 | 197,585,568 | 197,616,099 | 30,531 |
| 14–15 | 2 | 8,456,691 | 8,458,309 | 1,618 |
| 14–15 | 2 | 144,629,809 | 144,631,907 | 2,098 |
| 14–15 | 3 | 19,993,741 | 20,011,798 | 18,057 |
| 14–15 | 4 | 37,613,864 | 37,749,542 | 135,678 |
| 14–15 | 4 | 87,143,413 | 87,154,383 | 10,970 |
| 14–15 | 5 | 11,663,653 | 11,667,017 | 3,364 |
| 14–15 | 5 | 139,995,037 | 140,005,489 | 10,452 |
| 14–15 | 11 | 19,358,079 | 19,368,530 | 10,451 |

| | | | | |
|---|---|---|---|---|
| 14–15 | 14 | 48,439,012 | 48,453,207 | 14,195 |
| 14–15 | 14 | 124,050,257 | 124,063,977 | 13,720 |
| 14–15 | 15 | 10,789,096 | 10,824,524 | 35,428 |
| 14–15 | 15 | 59,395,818 | 59,406,319 | 10,501 |
| 14–15 | 16 | 55,911,540 | 55,916,486 | 4,946 |
| 14–15 | 17 | 9,709,889 | 9,739,767 | 29,878 |
| 14–15 | 18 | 13,604,969 | 13,605,075 | 106 |
| 14–15 | 20 | 13,929,433 | 13,931,658 | 2,225 |
| 14–15 | 20 | 90,849,602 | 90,859,242 | 9,640 |
| 14–15 | 21 | 35,333,015 | 35,355,379 | 22,364 |
| 14–16 | 1 | 43,694,315 | 43,709,110 | 14,795 |
| 14–16 | 2 | 147,763,738 | 147,766,802 | 3,064 |
| 14–16 | 3 | 137,625,741 | 137,680,193 | 54,452 |
| 14–16 | 4 | 61,071,987 | 65,493,613 | 4,421,626 |
| 14–16 | 5 | 29,344,905 | 29,361,278 | 16,373 |
| 14–16 | 6 | 36,338,661 | 36,347,744 | 9,083 |
| 14–16 | 6 | 65,614,501 | 65,633,742 | 19,241 |
| 14–16 | 7 | 99,425,496 | 99,445,307 | 19,811 |
| 14–16 | 9 | 19,712,186 | 19,784,191 | 72,005 |
| 14–16 | 10 | 25,529,422 | 25,553,446 | 24,024 |
| 14–16 | 10 | 29,007,204 | 29,098,364 | 91,160 |
| 14–16 | 11 | 37,368,428 | 37,432,943 | 64,515 |
| 14–16 | 12 | 44,852,803 | 44,858,690 | 5,887 |
| 14–16 | 12 | 114,568,713 | 114,573,362 | 4,649 |
| 14–16 | 12 | 139,349,218 | 139,350,548 | 1,330 |
| 14–16 | 12 | 151,668,776 | 151,671,029 | 2,253 |
| 14–16 | 12 | 201,218,641 | 201,220,554 | 1,913 |
| 14–16 | 13 | 102,649,340 | 102,652,630 | 3,290 |
| 14–16 | 14 | 120,374,572 | 120,468,868 | 94,296 |
| 14–16 | 14 | 124,700,148 | 124,837,031 | 136,883 |
| 14–16 | 15 | 53,488,161 | 53,495,531 | 7,370 |
| 14–16 | 16 | 3,216,768 | 3,216,937 | 169 |
| 14–16 | 16 | 92,429,510 | 92,429,851 | 341 |
| 14–16 | 17 | 11,920,897 | 11,923,484 | 2,587 |
| 14–16 | 17 | 64,027,583 | 64,036,226 | 8,643 |
| 14–16 | 18 | 13,662,475 | 13,663,319 | 844 |
| 14–16 | 19 | 3,235,525 | 3,236,245 | 720 |
| 14–16 | 19 | 55,643,688 | 55,645,116 | 1,428 |
| 14–17 | 1 | 13,179,217 | 13,192,758 | 13,541 |
| 14–17 | 1 | 51,095,186 | 51,117,025 | 21,839 |
| 14–17 | 1 | 180,806,508 | 180,814,256 | 7,748 |
| 14–17 | 2 | 146,000,909 | 146,002,301 | 1,392 |
| 14–17 | 3 | 25,454,828 | 25,501,931 | 47,103 |
| 14–17 | 3 | 146,027,752 | 146,032,560 | 4,808 |
| 14–17 | 4 | 76,799,712 | 76,822,792 | 23,080 |
| 14–17 | 5 | 103,302,003 | 103,312,472 | 10,469 |
| 14–17 | 6 | 74,268,982 | 74,370,138 | 101,156 |
| 14–17 | 6 | 78,207,103 | 78,216,928 | 9,825 |
| 14–17 | 7 | 24,735,233 | 24,741,037 | 5,804 |
| 14–17 | 7 | 122,817,744 | 122,823,623 | 5,879 |
| 14–17 | 8 | 80,960,129 | 80,967,123 | 6,994 |
| 14–17 | 9 | 82,624,922 | 82,991,921 | 366,999 |
| 14–17 | 9 | 91,115,821 | 91,239,633 | 123,812 |
| 14–17 | 9 | 98,276,610 | 98,287,590 | 10,980 |
| 14–17 | 10 | 41,318,667 | 41,338,706 | 20,039 |
| 14–17 | 10 | 57,961,205 | 57,969,300 | 8,095 |
| 14–17 | 11 | 22,017,800 | 22,027,182 | 9,382 |
| 14–17 | 12 | 61,965,134 | 61,978,465 | 13,331 |
| 14–17 | 12 | 99,910,096 | 99,913,603 | 3,507 |

| | | | | |
|---|---|---|---|---|
| 14–17 | 12 | 169,122,248 | 169,289,853 | 167,605 |
| 14–17 | 12 | 177,421,142 | 177,423,356 | 2,214 |
| 14–17 | 14 | 111,086,245 | 111,104,218 | 17,973 |
| 14–17 | 14 | 123,823,346 | 123,825,731 | 2,385 |
| 14–17 | 15 | 6,143,677 | 6,149,269 | 5,592 |
| 14–17 | 15 | 63,040,940 | 63,041,958 | 1,018 |
| 14–17 | 16 | 42,341,452 | 42,401,409 | 59,957 |
| 14–17 | 17 | 67,019,052 | 67,023,949 | 4,897 |
| 14–17 | 18 | 9,854,222 | 9,860,847 | 6,625 |
| 14–17 | 18 | 76,534,674 | 76,547,449 | 12,775 |
| 14–17 | 20 | 88,322,987 | 88,337,271 | 14,284 |
| 14–17 | 21 | 31,287,647 | 31,314,697 | 27,050 |

**Supplementary Table 8**. NCO events detected in the 22 meioses of this study. Individual meioses are indicated by alternating shading.

| meiosis | chromosome | start | end | # markers | tract length (bp) | REF alleles | ALT alleles |
|---|---|---|---|---|---|---|---|
| 1–3 | 16 | 67,759,704 | 67,770,370 | 9 | 10,667 | A,C,G,G,G,A,G,G,T | G,T,T,T,A,G,A,A,C |
| 1–4 | 5 | 27,099,378 | 27,099,378 | 1 | 1 | G | A |
| 1–4 | 13 | 66,884,821 | 66,884,821 | 1 | 1 | G | A |
| 1–4 | 16 | 40,421,191 | 40,421,191 | 1 | 1 | C | G |
| 1–5 | 12 | 21,709,900 | 21,709,900 | 1 | 1 | C | T |
| 1–5 | 20 | 87,917,254 | 87,917,254 | 1 | 1 | A | G |
| 1–5 | 21 | 61,103,156 | 61,103,156 | 1 | 1 | C | T |
| 1–6 | 4 | 1,291,632 | 1,291,632 | 1 | 1 | C | T |
| 1–6 | 5 | 113,530,330 | 113,530,330 | 1 | 1 | G | A |
| 1–6 | 5 | 137,387,371 | 137,387,371 | 1 | 1 | C | T |
| 1–6 | 5 | 149,075,267 | 149,075,267 | 1 | 1 | A | G |
| 1–6 | 9 | 65,371,165 | 65,371,165 | 1 | 1 | A | T |
| 1–6 | 10 | 28,863,467 | 28,863,467 | 1 | 1 | C | A |
| 1–6 | 12 | 49,614,053 | 49,614,053 | 1 | 1 | G | A |
| 1–6 | 13 | 59,167,798 | 59,167,798 | 1 | 1 | T | C |
| 1–6 | 18 | 40,297,953 | 40,297,953 | 1 | 1 | A | G |
| 2–3 | 4 | 80,523,567 | 80,523,567 | 1 | 1 | A | G |
| 2–3 | 6 | 45,593,300 | 45,593,300 | 1 | 1 | A | C |
| 2–3 | 8 | 2,522,023 | 2,522,023 | 1 | 1 | G | T |
| 2–3 | 12 | 11,494,726 | 11,494,726 | 1 | 1 | C | T |
| 2–3 | 13 | 43,313,207 | 43,313,207 | 1 | 1 | G | T |
| 2–3 | 13 | 51,984,454 | 51,984,454 | 1 | 1 | C | T |
| 2–4 | 3 | 134,094,740 | 134,094,740 | 1 | 1 | T | C |
| 2–4 | 11 | 3,138,514 | 3,138,514 | 1 | 1 | C | T |
| 2–4 | 15 | 30,609,330 | 30,609,330 | 1 | 1 | T | C |
| 2–4 | 16 | 71,601,679 | 71,601,679 | 1 | 1 | G | A |
| 2–4 | 18 | 15,601,590 | 15,601,590 | 1 | 1 | C | T |
| 2–4 | 18 | 24,179,479 | 24,179,479 | 1 | 1 | C | T |
| 2–4 | 22 | 6,039,057 | 6,039,057 | 1 | 1 | T | C |
| 2–5 | 2 | 81,822,540 | 81,822,540 | 1 | 1 | T | C |
| 2–5 | 2 | 81,822,540 | 81,822,540 | 1 | 1 | T | C |
| 2–5 | 4 | 36,341,579 | 36,341,579 | 1 | 1 | G | A |
| 2–5 | 4 | 36,341,579 | 36,341,579 | 1 | 1 | G | A |
| 2–5 | 4 | 130,531,469 | 130,531,469 | 1 | 1 | G | A |
| 2–5 | 4 | 130,531,469 | 130,531,469 | 1 | 1 | G | A |
| 2–5 | 5 | 97,175,726 | 97,175,726 | 1 | 1 | A | C |
| 2–5 | 5 | 97,175,726 | 97,175,726 | 1 | 1 | A | C |

| | | | | | | | |
|---|---|---|---|---|---|---|---|
| 2–5 | 6 | 45,436,838 | 45,436,838 | 1 | 1 | A | C |
| 2–5 | 6 | 45,436,838 | 45,436,838 | 1 | 1 | A | C |
| 2–6 | 4 | 130,910,511 | 130,910,511 | 1 | 1 | C | T |
| 2–6 | 7 | 99,601,782 | 99,601,782 | 1 | 1 | G | C |
| 2–6 | 9 | 89,058,189 | 89,058,189 | 1 | 1 | C | T |
| 2–6 | 12 | 59,778,040 | 59,778,040 | 1 | 1 | T | C |
| 2–6 | 15 | 86,437,202 | 86,437,203 | 2 | 2 | T,G | C,C |
| 2–6 | 16 | 67,800,688 | 67,800,688 | 1 | 1 | T | C |
| 2–6 | 18 | 9,469,651 | 9,469,651 | 1 | 1 | G | A |
| 2–6 | 22 | 7,905,430 | 7,905,430 | 1 | 1 | C | T |
| 7–9 | 2 | 120,926,264 | 120,926,264 | 1 | 1 | G | A |
| 7–9 | 10 | 3,675,421 | 3,675,421 | 1 | 1 | A | T |
| 7–9 | 10 | 13,431,127 | 13,431,127 | 1 | 1 | A | G |
| 7–9 | 10 | 61,158,915 | 61,158,966 | 2 | 52 | G,G | A,T |
| 7–9 | 10 | 62,056,545 | 62,056,545 | 1 | 1 | T | C |
| 7–9 | 12 | 146,149,238 | 146,149,238 | 1 | 1 | T | C |
| 7–9 | 12 | 168,669,630 | 168,710,265 | 2 | 40,636 | G,T | A,C |
| 7–9 | 16 | 52,264,274 | 52,264,274 | 1 | 1 | A | G |
| 7–9 | 16 | 84,872,395 | 84,872,395 | 1 | 1 | A | G |
| 7–9 | 19 | 191,705 | 191,710 | 2 | 6 | G,A | T,G |
| 7–10 | 2 | 113,819,615 | 113,819,615 | 1 | 1 | G | A |
| 7–10 | 6 | 56,561,963 | 56,561,964 | 2 | 2 | T,T | G,G |
| 7–10 | 12 | 34,753,613 | 34,753,613 | 1 | 1 | T | C |
| 7–10 | 13 | 18,065,493 | 18,065,493 | 1 | 1 | C | T |
| 7–10 | 13 | 39,364,793 | 39,364,793 | 1 | 1 | G | A |
| 7–10 | 16 | 47,992,791 | 47,992,791 | 1 | 1 | T | C |
| 7–10 | 18 | 48,453,998 | 48,453,998 | 1 | 1 | T | C |
| 7–10 | 19 | 50,420,140 | 50,420,140 | 1 | 1 | A | T |
| 7–10 | 19 | 54,980,592 | 54,980,592 | 1 | 1 | C | T |
| 7–11 | 1 | 71,166,163 | 71,166,163 | 1 | 1 | C | T |
| 7–11 | 1 | 114,698,091 | 114,698,091 | 1 | 1 | C | A |
| 7–11 | 1 | 128,165,981 | 128,165,981 | 1 | 1 | T | C |
| 7–11 | 3 | 88,390,168 | 88,390,168 | 1 | 1 | T | A |
| 7–11 | 4 | 56,658,754 | 56,658,754 | 1 | 1 | G | A |
| 7–11 | 5 | 32,686,194 | 32,686,194 | 1 | 1 | A | G |
| 7–11 | 5 | 104,664,696 | 104,664,696 | 1 | 1 | G | A |
| 7–11 | 8 | 95,454,272 | 95,454,272 | 1 | 1 | C | A |
| 7–11 | 10 | 54,354,251 | 54,354,251 | 1 | 1 | T | C |
| 7–11 | 11 | 21,909,977 | 21,909,977 | 1 | 1 | C | T |

| | | | | | | | |
|---|---|---|---|---|---|---|---|
| 7–11 | 12 | 66,851,882 | 66,851,882 | 1 | 1 | A | G |
| 7–11 | 12 | 141,079,294 | 141,079,294 | 1 | 1 | A | T |
| 7–11 | 13 | 12,548,341 | 12,548,341 | 1 | 1 | C | T |
| 7–11 | 16 | 7,850,568 | 7,850,568 | 1 | 1 | C | T |
| 7–11 | 16 | 90,128,074 | 90,128,074 | 1 | 1 | G | A |
| 7–11 | 17 | 25,437,701 | 25,437,701 | 1 | 1 | C | G |
| 7–12 | 1 | 50,934,775 | 50,934,775 | 1 | 1 | A | G |
| 7–12 | 1 | 78,197,745 | 78,197,745 | 1 | 1 | A | G |
| 7–12 | 5 | 107,781,405 | 107,781,405 | 1 | 1 | T | C |
| 7–12 | 6 | 12,722,834 | 12,722,834 | 1 | 1 | A | G |
| 7–12 | 7 | 115,750,910 | 115,750,910 | 1 | 1 | G | A |
| 7–12 | 12 | 17,586,789 | 17,586,789 | 1 | 1 | T | C |
| 7–12 | 17 | 50,237,578 | 50,237,578 | 1 | 1 | T | C |
| 7–12 | 20 | 41,156,375 | 41,156,375 | 1 | 1 | G | A |
| 7–12 | 22 | 14,595,061 | 14,595,061 | 1 | 1 | C | T |
| 8–9 | 4 | 56,235,740 | 56,235,740 | 1 | 1 | C | T |
| 8–9 | 5 | 23,497,759 | 23,497,759 | 1 | 1 | C | T |
| 8–9 | 5 | 30,041,365 | 30,041,365 | 1 | 1 | C | T |
| 8–9 | 5 | 67,518,353 | 67,518,353 | 1 | 1 | A | G |
| 8–9 | 6 | 65,409,747 | 65,409,990 | 2 | 244 | G,G | T,A |
| 8–9 | 8 | 745,287 | 745,287 | 1 | 1 | T | C |
| 8–9 | 8 | 3,964,279 | 3,964,279 | 1 | 1 | A | G |
| 8–9 | 8 | 58,110,489 | 58,110,489 | 1 | 1 | C | T |
| 8–9 | 16 | 24,082,982 | 24,082,982 | 1 | 1 | G | T |
| 8–9 | 18 | 48,456,435 | 48,456,529 | 2 | 95 | C,C | T,A |
| 8–9 | 18 | 56,606,281 | 56,606,281 | 1 | 1 | T | A |
| 8–9 | 20 | 79,053,137 | 79,053,137 | 1 | 1 | A | G |
| 8–9 | 22 | 5,994,104 | 5,994,104 | 1 | 1 | T | C |
| 8–10 | 2 | 52,796,977 | 52,796,977 | 1 | 1 | T | C |
| 8–10 | 2 | 81,156,195 | 81,156,195 | 1 | 1 | T | C |
| 8–10 | 3 | 70,293,515 | 70,293,515 | 1 | 1 | G | T |
| 8–10 | 4 | 82,176,734 | 82,176,736 | 2 | 3 | A,G | T,C |
| 8–10 | 8 | 71,597,994 | 71,597,994 | 1 | 1 | C | T |
| 8–10 | 11 | 1,019,220 | 1,019,220 | 1 | 1 | G | A |
| 8–11 | 1 | 4,083,928 | 4,083,928 | 1 | 1 | C | T |
| 8–11 | 3 | 59,652,228 | 59,652,228 | 1 | 1 | A | T |
| 8–11 | 3 | 114,945,079 | 114,945,079 | 1 | 1 | T | C |
| 8–11 | 4 | 107,442,634 | 107,442,634 | 1 | 1 | T | C |
| 8–11 | 6 | 65,967,506 | 65,967,506 | 1 | 1 | T | C |

| | | | | | | | |
|---|---|---|---|---|---|---|---|
| 8–11 | 12 | 170,591,861 | 170,591,861 | 1 | 1 | C | T |
| 8–11 | 12 | 176,188,745 | 176,188,745 | 1 | 1 | C | T |
| 8–11 | 12 | 191,083,153 | 191,083,153 | 1 | 1 | T | C |
| 8–11 | 12 | 199,623,479 | 199,623,479 | 1 | 1 | C | T |
| 8–11 | 14 | 15,719,160 | 15,719,160 | 1 | 1 | G | C |
| 8–11 | 15 | 9,693,142 | 9,693,142 | 1 | 1 | T | C |
| 8–11 | 16 | 57,155,592 | 57,155,592 | 1 | 1 | A | G |
| 8–11 | 18 | 28,840,832 | 28,840,832 | 1 | 1 | A | G |
| 8–11 | 19 | 52,127,857 | 52,127,857 | 1 | 1 | G | A |
| 8–11 | 19 | 54,133,071 | 54,133,071 | 1 | 1 | C | T |
| 8–11 | 21 | 22,618,937 | 22,618,937 | 1 | 1 | G | C |
| 8–12 | 7 | 8,569,610 | 8,569,610 | 1 | 1 | G | T |
| 8–12 | 7 | 43,031,210 | 43,031,210 | 1 | 1 | A | C |
| 8–12 | 8 | 6,834,961 | 6,834,961 | 1 | 1 | T | C |
| 8–12 | 20 | 68,955,322 | 68,955,332 | 2 | 11 | G,A | A,G |
| 13–15 | 3 | 129,645,599 | 129,645,599 | 1 | 1 | C | T |
| 13–15 | 8 | 112,529,513 | 112,529,513 | 1 | 1 | T | G |
| 13–15 | 10 | 27,815,219 | 27,815,219 | 1 | 1 | A | G |
| 13–15 | 11 | 25,940,381 | 25,940,381 | 1 | 1 | G | A |
| 13–15 | 11 | 46,035,315 | 46,035,315 | 1 | 1 | C | T |
| 13–15 | 11 | 48,083,623 | 48,083,623 | 1 | 1 | C | T |
| 13–15 | 13 | 93,622,683 | 93,622,683 | 1 | 1 | C | G |
| 13–15 | 20 | 90,517,344 | 90,517,344 | 1 | 1 | T | C |
| 13–16 | 3 | 25,089,271 | 25,089,271 | 1 | 1 | A | G |
| 13–16 | 3 | 133,001,747 | 133,001,747 | 1 | 1 | A | T |
| 13–16 | 3 | 154,541,371 | 154,541,403 | 3 | 33 | C,C,A | T,T,G |
| 13–16 | 6 | 77,767,231 | 77,767,231 | 1 | 1 | C | T |
| 13–16 | 8 | 29,414,510 | 29,414,510 | 1 | 1 | C | T |
| 13–16 | 9 | 44,322,965 | 44,322,965 | 1 | 1 | A | C |
| 13–16 | 10 | 9,661,662 | 9,661,662 | 1 | 1 | G | A |
| 13–16 | 10 | 50,127,334 | 50,127,334 | 1 | 1 | C | T |
| 13–16 | 17 | 706,841 | 707,191 | 4 | 350 | C,A,A,T | T,G,G,C |
| 13–16 | 18 | 81,547,240 | 81,547,240 | 1 | 1 | A | G |
| 13–16 | 21 | 391,877 | 391,987 | 2 | 111 | C,T | A,G |
| 13–17 | 3 | 135,054,018 | 135,054,028 | 2 | 11 | C,A | T,C |
| 13–17 | 8 | 37,053,942 | 37,053,942 | 1 | 1 | G | A |
| 13–17 | 14 | 9,294,937 | 9,294,937 | 1 | 1 | A | T |
| 13–17 | 17 | 6,185,751 | 6,185,751 | 1 | 1 | T | C |
| 13–17 | 19 | 39,376,799 | 39,376,799 | 1 | 1 | C | T |

| | | | | | | | |
|---|---|---|---|---|---|---|---|
| 13–17 | 21 | 53,825,838 | 53,825,838 | 1 | 1 | A | G |
| 14–15 | 1 | 23,343,196 | 23,343,196 | 1 | 1 | G | A |
| 14–15 | 2 | 2,301,866 | 2,301,874 | 2 | 9 | A,A | G,G |
| 14–15 | 2 | 78,610,397 | 78,630,505 | 2 | 20,109 | G,T | A,C |
| 14–15 | 6 | 92,562,985 | 92,562,985 | 1 | 1 | T | G |
| 14–15 | 7 | 127,012,832 | 127,012,832 | 1 | 1 | G | T |
| 14–15 | 8 | 41,419,376 | 41,419,376 | 1 | 1 | T | G |
| 14–15 | 8 | 71,791,500 | 71,791,500 | 1 | 1 | A | G |
| 14–15 | 11 | 7,564,947 | 7,564,947 | 1 | 1 | T | G |
| 14–15 | 12 | 154,610,198 | 154,610,198 | 1 | 1 | G | A |
| 14–15 | 12 | 177,542,447 | 177,542,447 | 1 | 1 | C | G |
| 14–15 | 12 | 195,659,476 | 195,659,476 | 1 | 1 | T | C |
| 14–15 | 13 | 2,876,125 | 2,876,125 | 1 | 1 | T | C |
| 14–15 | 19 | 51,325,070 | 51,325,070 | 1 | 1 | G | C |
| 14–15 | 20 | 5,230,149 | 5,230,153 | 2 | 5 | C,C | T,T |
| 14–16 | 6 | 123,243,333 | 123,243,333 | 1 | 1 | A | T |
| 14–16 | 11 | 10,981,572 | 10,981,572 | 1 | 1 | C | A |
| 14–16 | 16 | 19,319,138 | 19,319,368 | 8 | 231 | C,C,T,G,T,A,T,G | T,T,C,A,G,C,A,T |
| 14–16 | 16 | 39,374,336 | 39,374,336 | 1 | 1 | C | T |
| 14–16 | 16 | 40,467,461 | 40,467,461 | 1 | 1 | C | G |
| 14–16 | 17 | 61,690,647 | 61,690,647 | 1 | 1 | A | G |
| 14–17 | 1 | 117,918,370 | 117,918,370 | 1 | 1 | G | A |
| 14–17 | 1 | 160,386,962 | 160,386,962 | 1 | 1 | G | T |
| 14–17 | 2 | 23,478,120 | 23,478,120 | 1 | 1 | T | C |
| 14–17 | 3 | 93,833,617 | 93,833,617 | 1 | 1 | C | T |
| 14–17 | 6 | 35,321,175 | 35,321,175 | 1 | 1 | G | A |
| 14–17 | 6 | 113,933,380 | 113,933,449 | 2 | 70 | G,A | A,G |
| 14–17 | 9 | 95,554,884 | 95,571,494 | 23 | 16,611 | A,G,A,C,C,T,G,C,T,C,G,C,<br>G,G,G,C,G,G,G,G,A,G,C | T,A,G,G,G,G,T,A,C,T,A,T,<br>C,A,A,T,A,A,A,A,G,A,T |
| 14–17 | 9 | 98,797,581 | 98,797,638 | 2 | 58 | G,C | A,T |
| 14–17 | 10 | 43,011,886 | 43,011,886 | 1 | 1 | G | T |
| 14–17 | 14 | 47,774,749 | 47,774,749 | 1 | 1 | G | A |
| 14–17 | 14 | 127,738,669 | 127,738,669 | 1 | 1 | A | G |
| 14–17 | 16 | 12,999,791 | 12,999,791 | 1 | 1 | C | T |
| 14–17 | 20 | 58,710,079 | 58,710,079 | 1 | 1 | C | T |
| 14–17 | 22 | 14,556,506 | 14,556,506 | 1 | 1 | G | A |

**Supplementary Figure 1**.

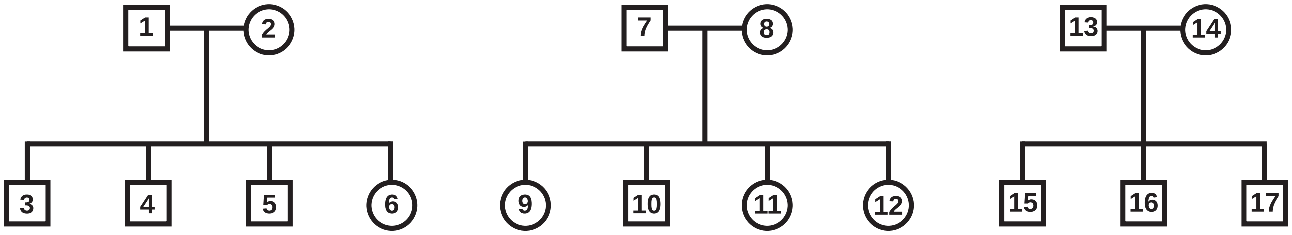


Pedigree structure of the 17 individuals included in the study.

**Supplementary Figure 2**.

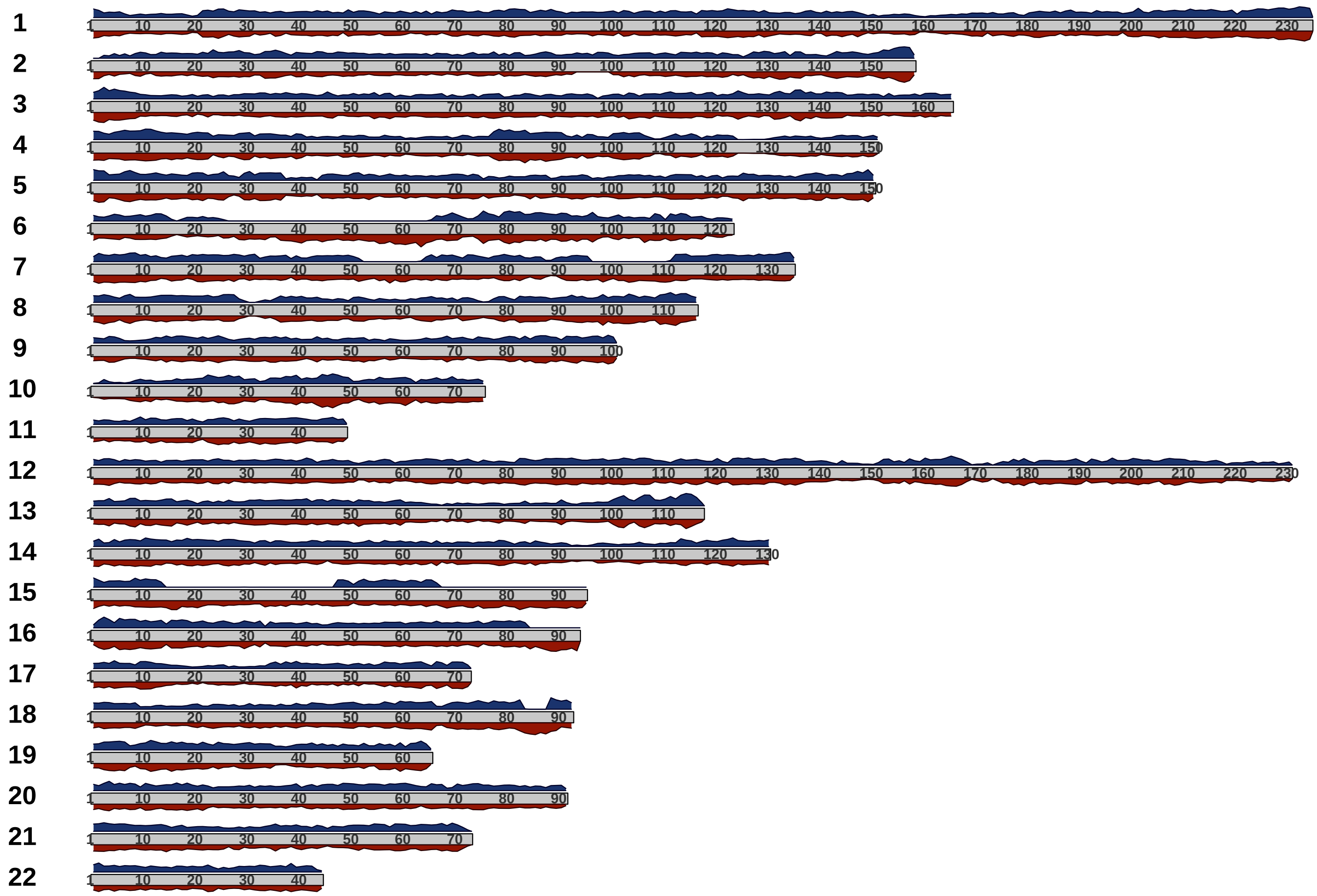


Distribution of phase-informative markers across the autosomes (chromosomes 1-22) in family 1–2, with peak height being proportional to marker density.

**Supplementary Figure 3**.

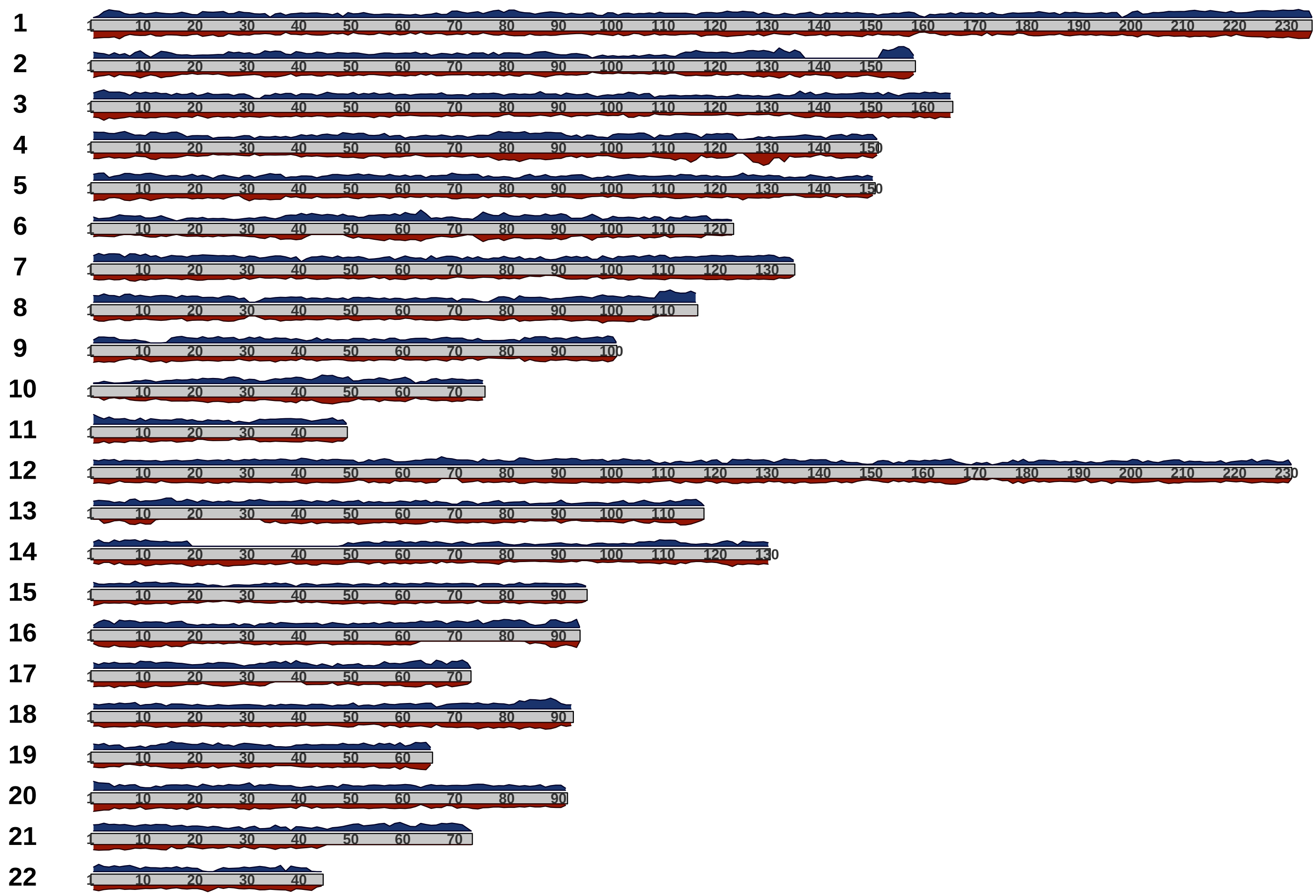

Distribution of phase-informative markers across the autosomes (chromosomes 1-22) in family 7–8, with peak height being proportional to marker density.

**Supplementary Figure 4**.

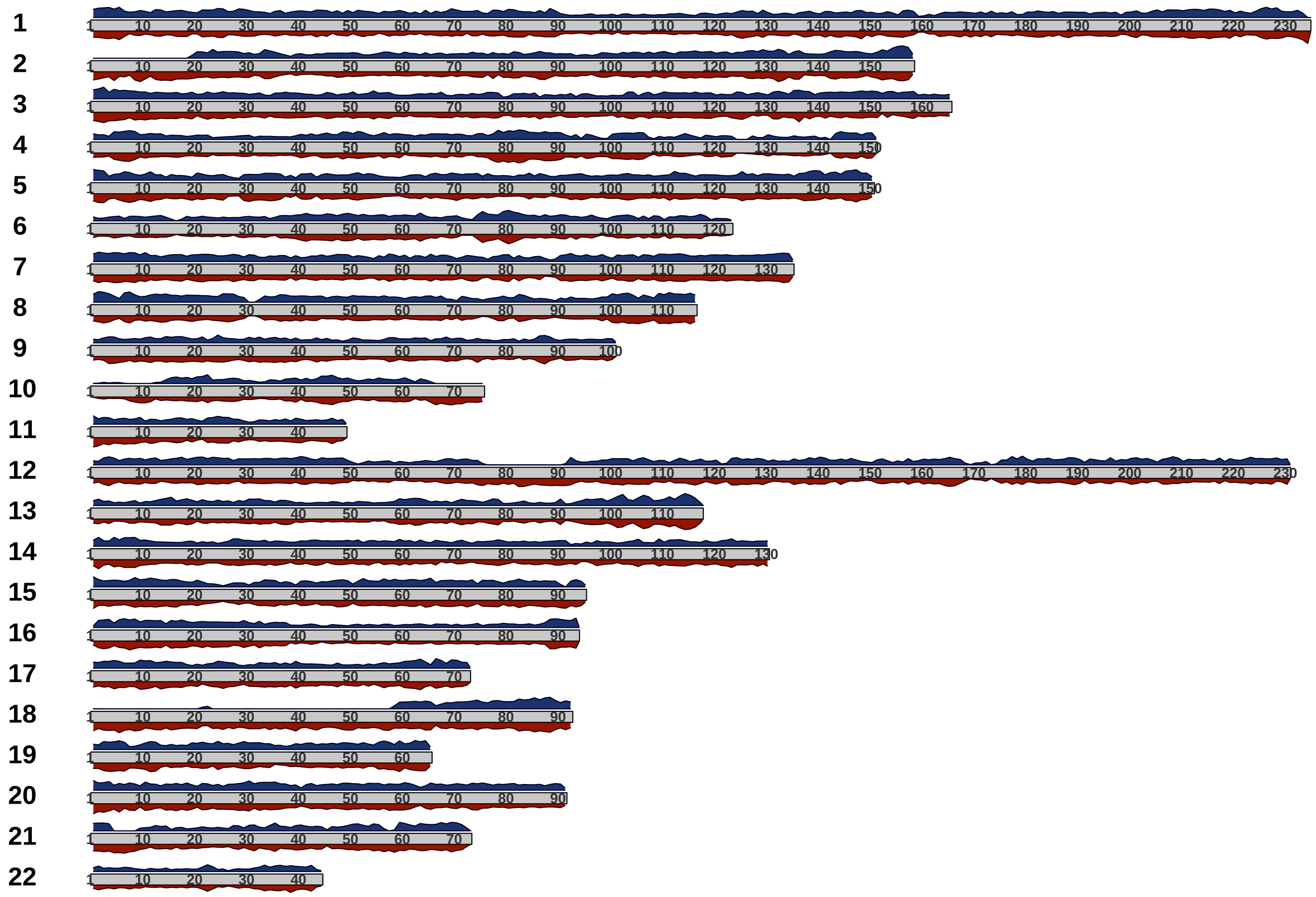


Distribution of phase-informative markers across the autosomes (chromosomes 1-22) in family 13–14, with peak height being proportional to marker density.

**Supplementary Figure 5**.

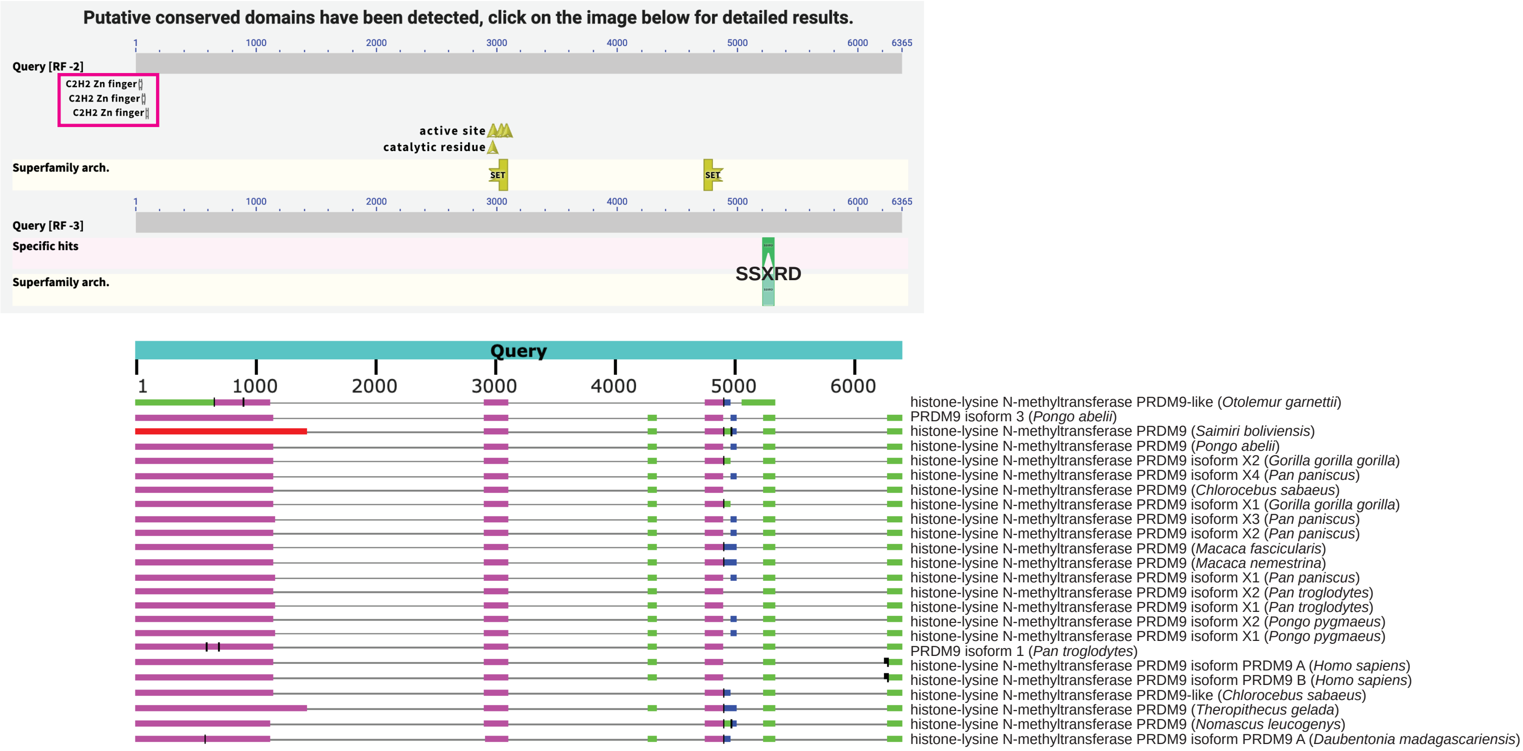


A BLASTX search identified a partial PRDM9 sequence in coppery titi monkey including the SSXRD, PR/SET, and zinc-finger domains (note that no KRAB domain could be identified).

**Supplementary Figure 6**.

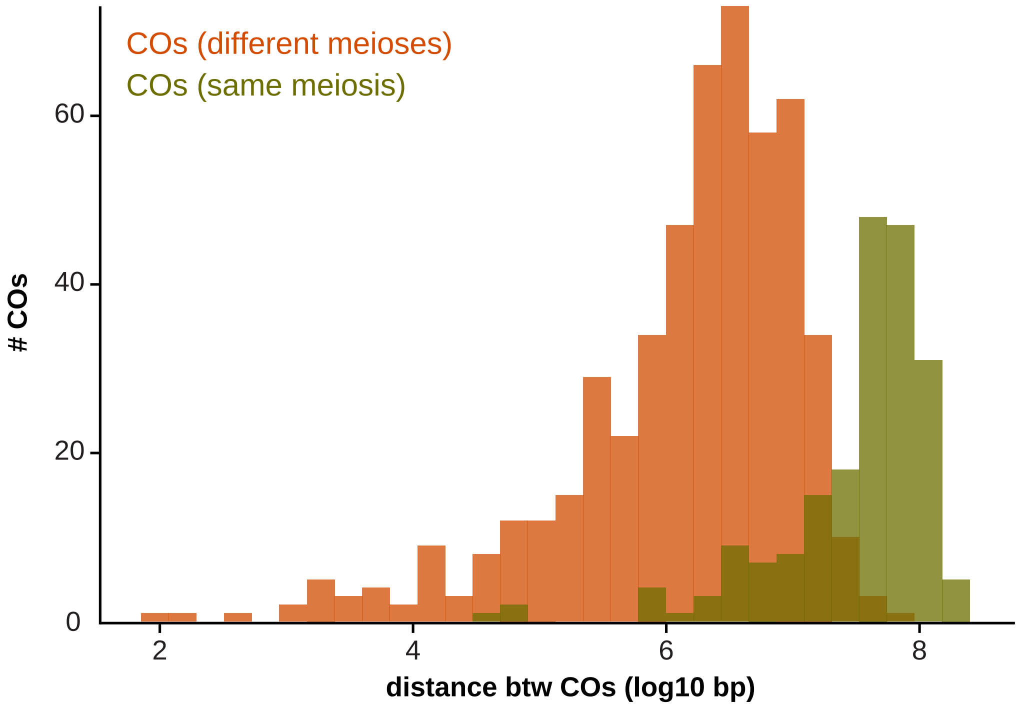


Distribution of the log-scaled distances between crossovers (COs) occurring in the same meiosis and those occurring in different meioses (shown in olive and orange, respectively).

**Supplementary Figure 7**.

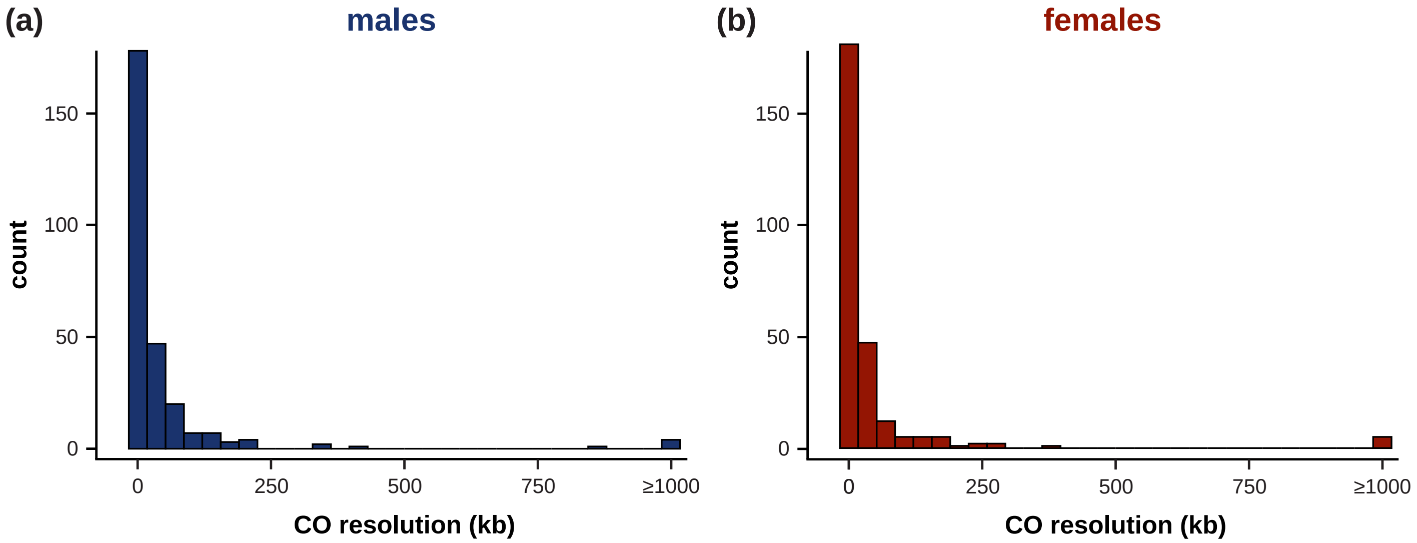


Distribution of the resolution of (a) paternally-inherited COs (shown in blue) and (b) maternally-inherited COs (shown in red).

**Supplementary Figure 8**.

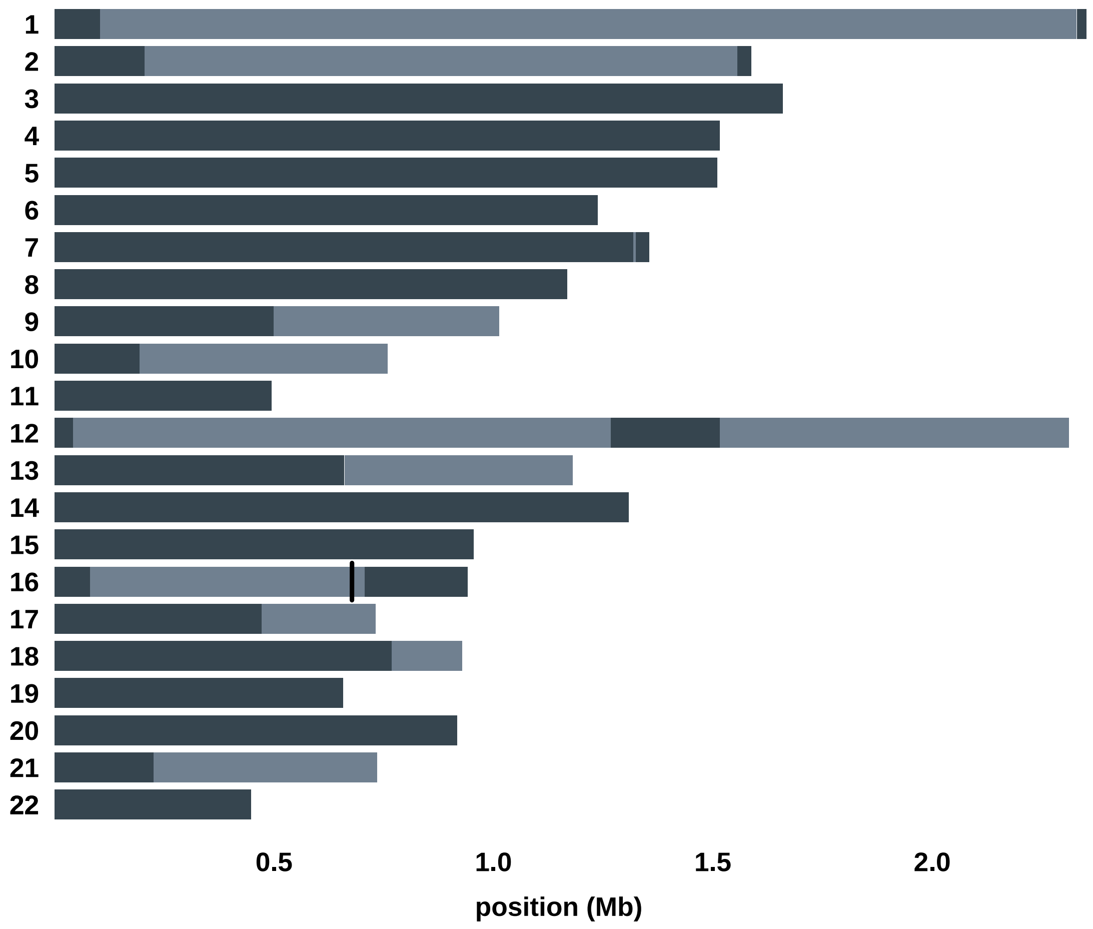


Haplotype blocks observed on the autosomes (chromosomes 1-22) in meiosis 1–3. Haplotype blocks are shown in alternating gray shading (when applicable, gaps are shown in white). NCO events are indicated by vertical black lines.

**Supplementary Figure 9**.

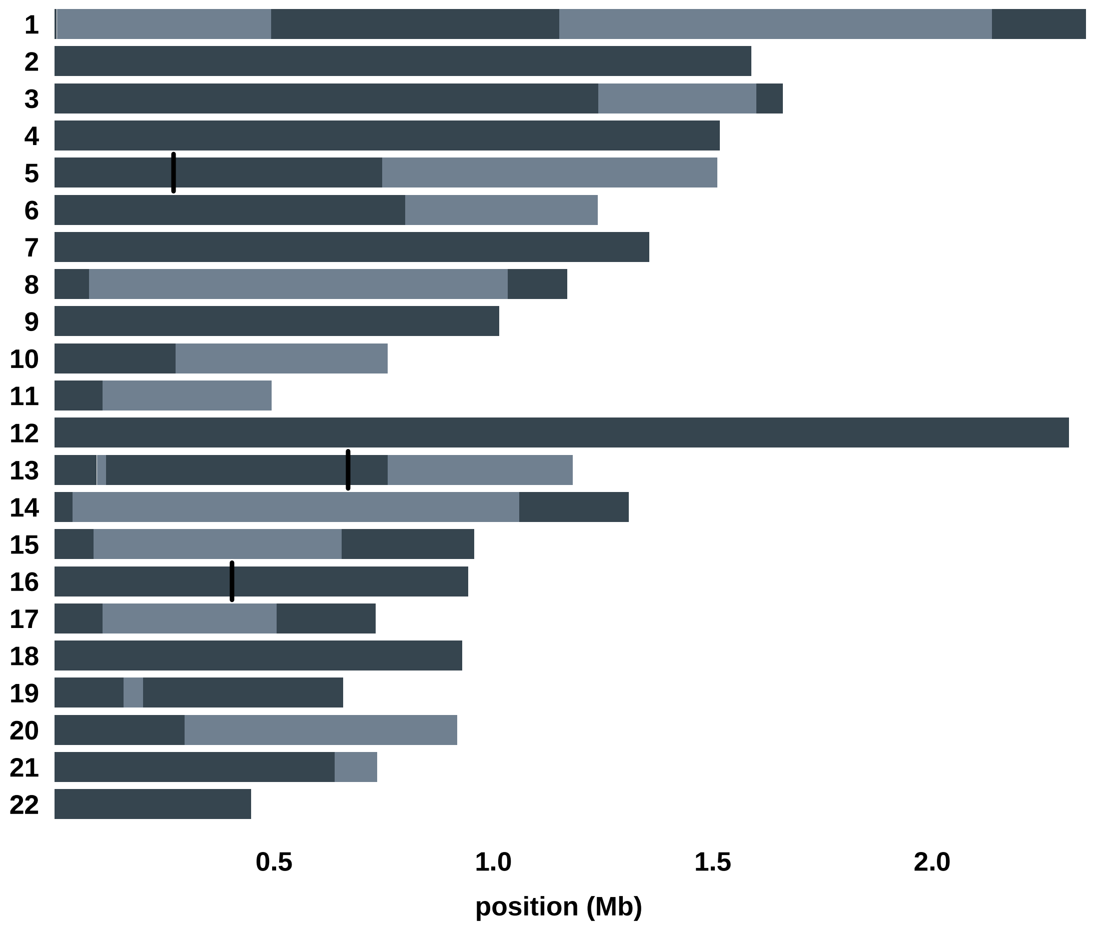


Haplotype blocks observed on the autosomes (chromosomes 1-22) in meiosis 1–4. Haplotype blocks are shown in alternating gray shading (when applicable, gaps are shown in white). NCO events are indicated by vertical black lines.

**Supplementary Figure 10**.

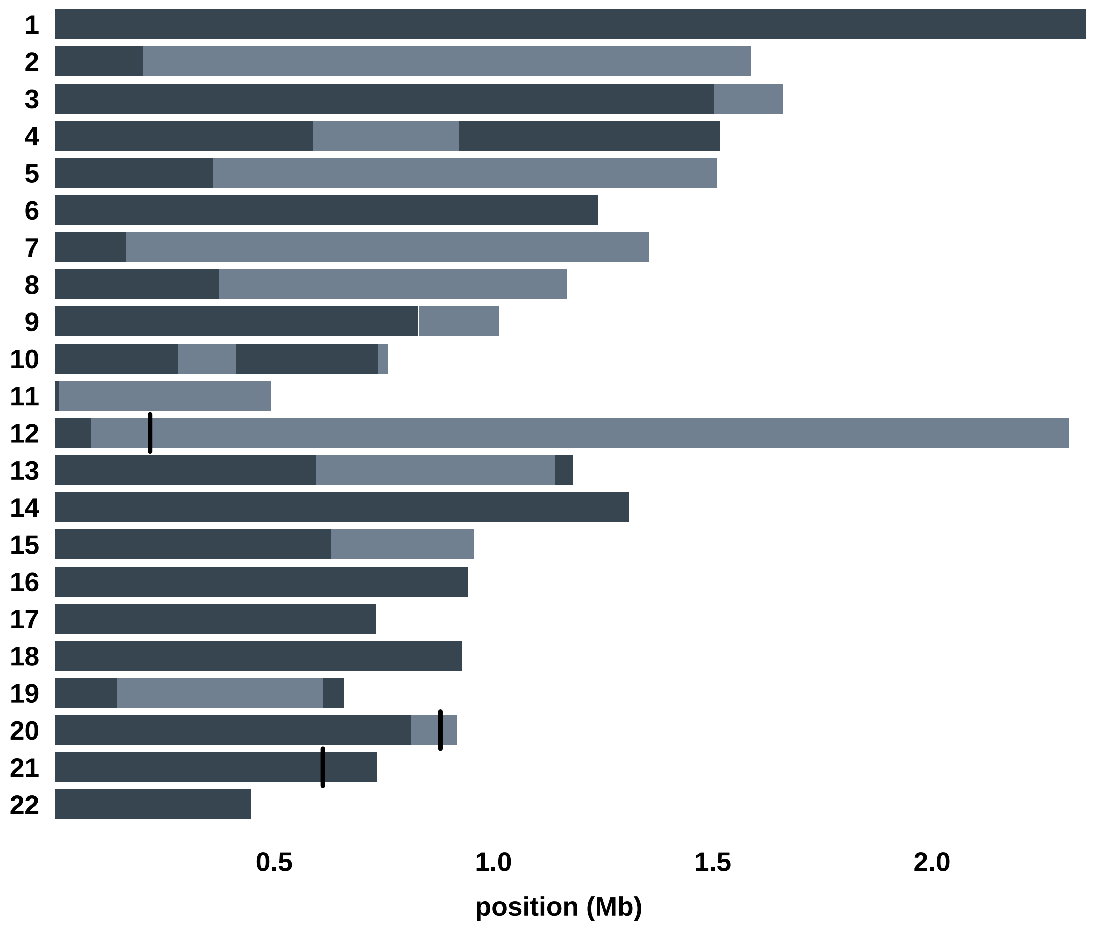


Haplotype blocks observed on the autosomes (chromosomes 1-22) in meiosis 1–5. Haplotype blocks are shown in alternating gray shading (when applicable, gaps are shown in white). NCO events are indicated by vertical black lines.

**Supplementary Figure 11**.

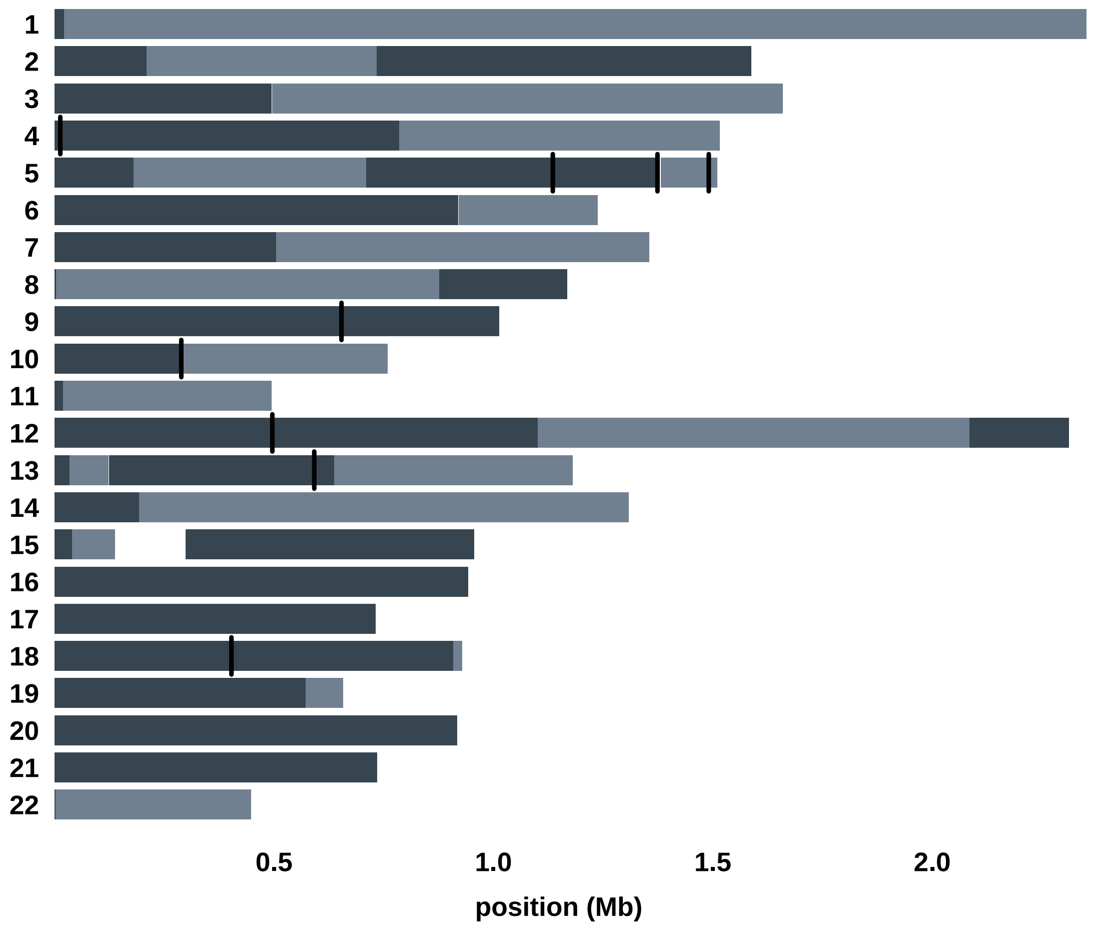


Haplotype blocks observed on the autosomes (chromosomes 1-22) in meiosis 1–6. Haplotype blocks are shown in alternating gray shading (when applicable, gaps are shown in white). NCO events are indicated by vertical black lines.

**Supplementary Figure 12**.

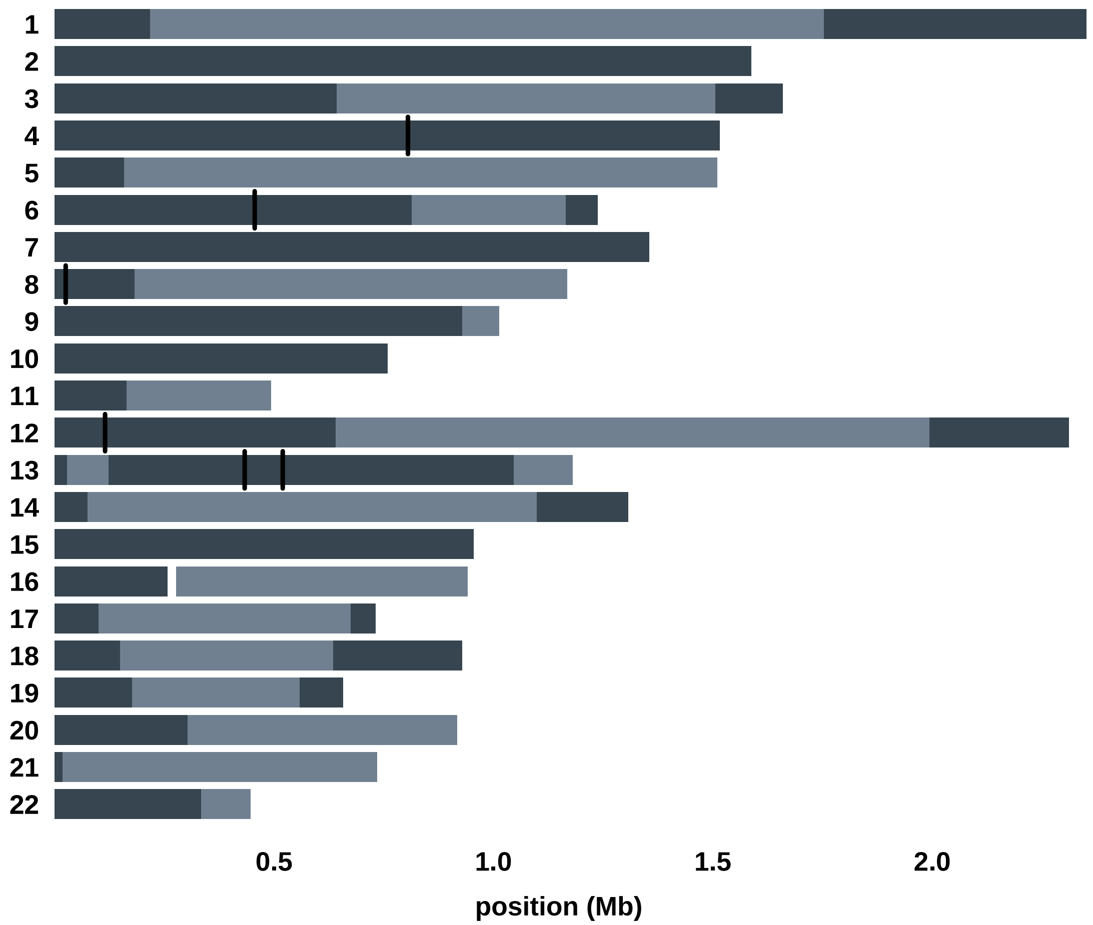


Haplotype blocks observed on the autosomes (chromosomes 1-22) in meiosis 2–3. Haplotype blocks are shown in alternating gray shading (when applicable, gaps are shown in white). NCO events are indicated by vertical black lines.

**Supplementary Figure 13**.

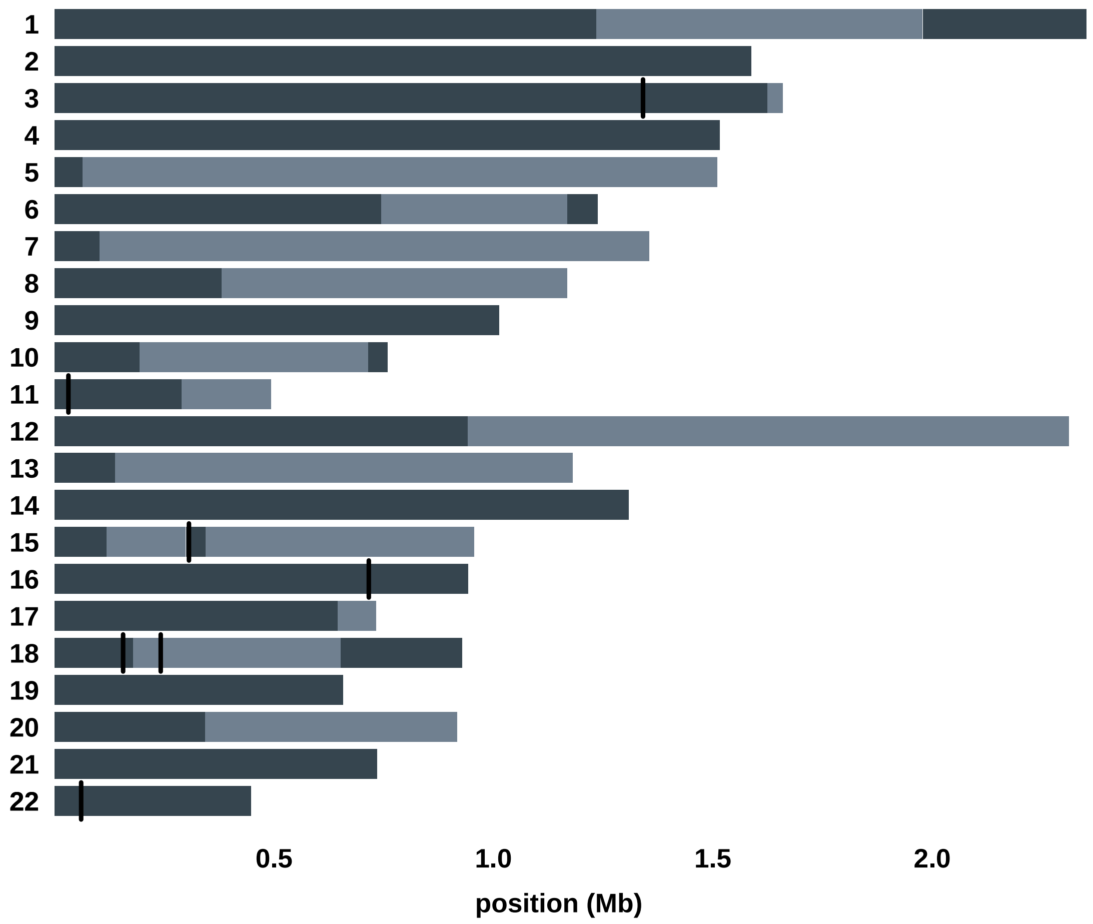


Haplotype blocks observed on the autosomes (chromosomes 1-22) in meiosis 2–4. Haplotype blocks are shown in alternating gray shading (when applicable, gaps are shown in white). NCO events are indicated by vertical black lines.

**Supplementary Figure 14.**

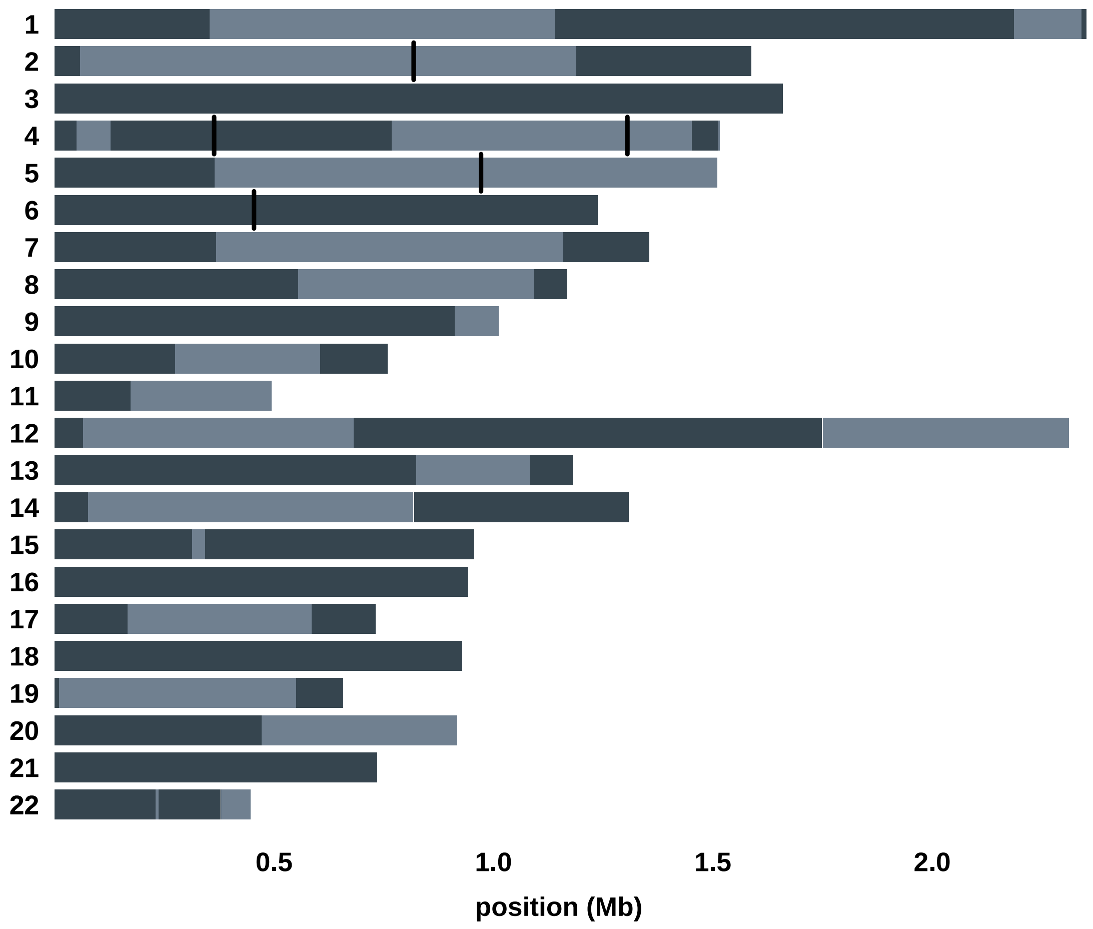


Haplotype blocks observed on the autosomes (chromosomes 1-22) in meiosis 2–5. Haplotype blocks are shown in alternating gray shading (when applicable, gaps are shown in white). NCO events are indicated by vertical black lines.

**Supplementary Figure 15**.

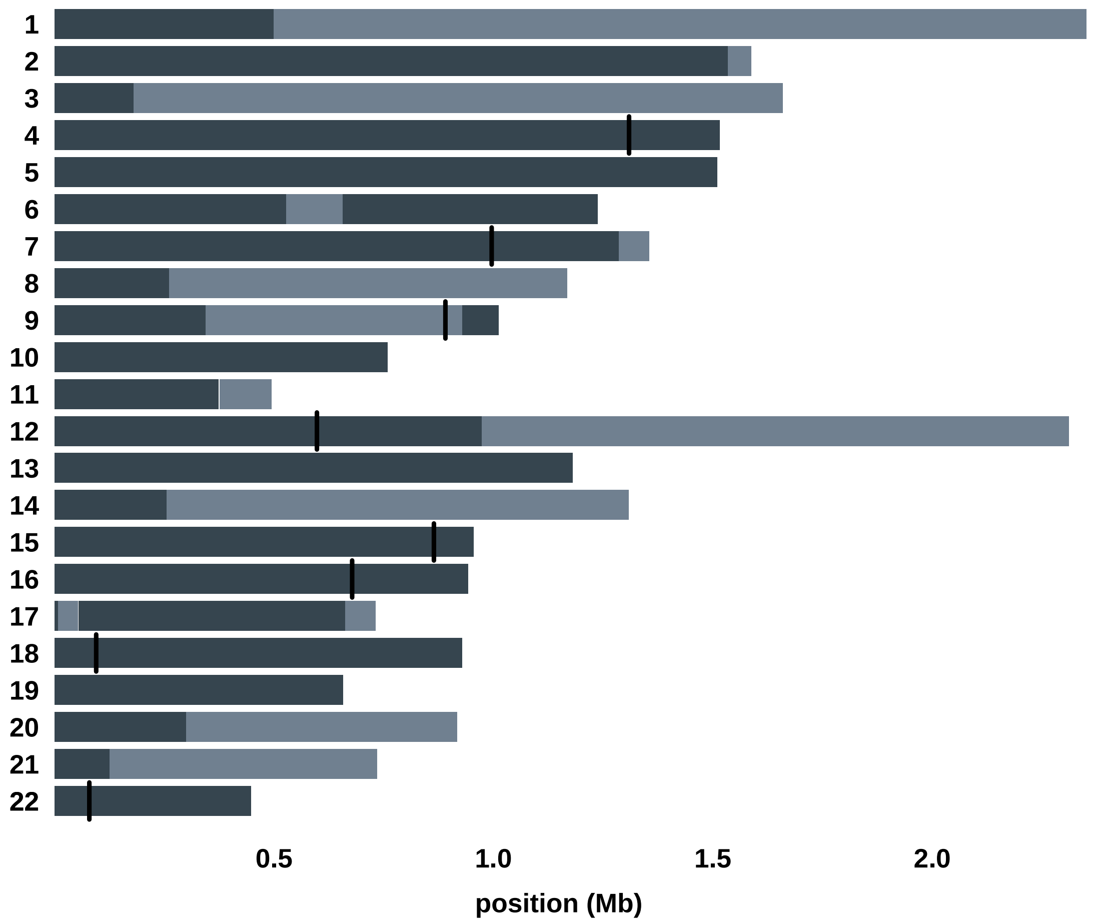


Haplotype blocks observed on the autosomes (chromosomes 1-22) in meiosis 2–6. Haplotype blocks are shown in alternating gray shading (when applicable, gaps are shown in white). NCO events are indicated by vertical black lines.

**Supplementary Figure 16**.

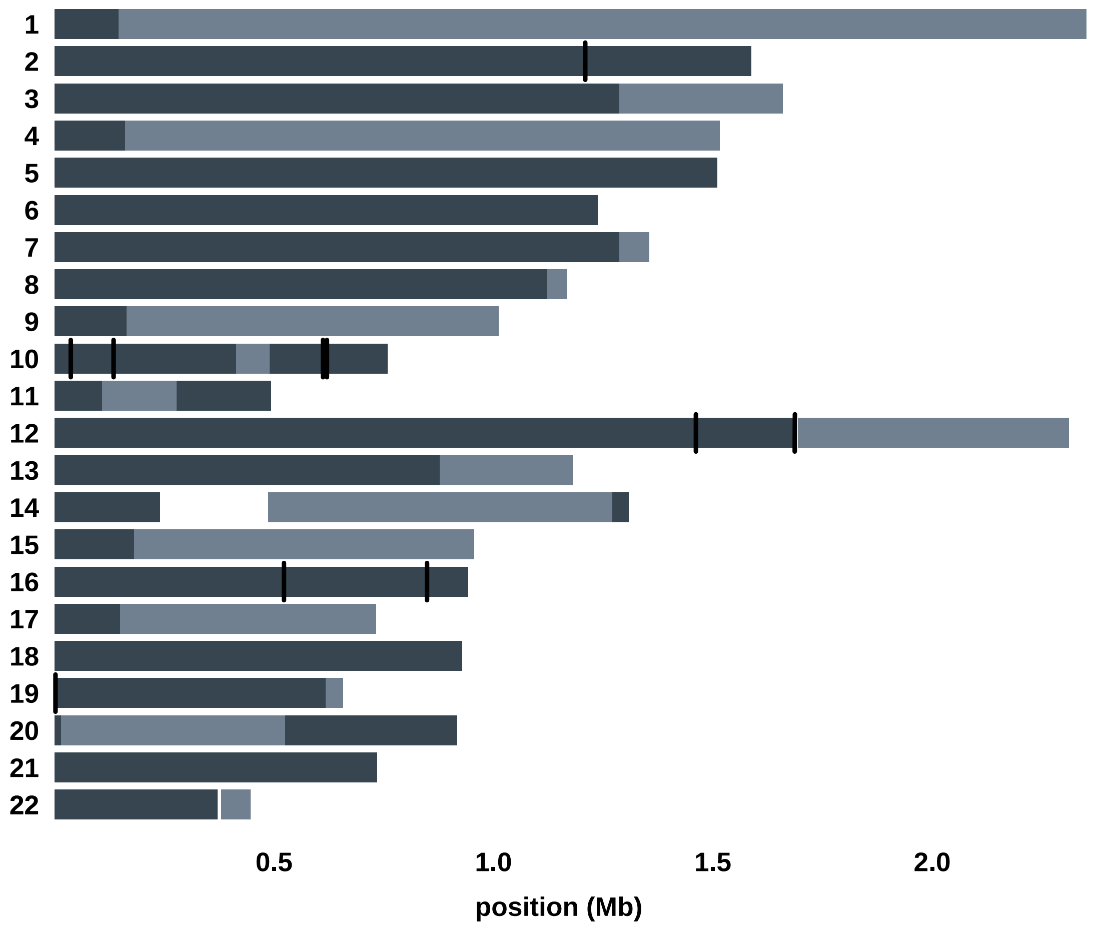


Haplotype blocks observed on the autosomes (chromosomes 1-22) in meiosis 7–9. Haplotype blocks are shown in alternating gray shading (when applicable, gaps are shown in white). NCO events are indicated by vertical black lines.

**Supplementary Figure 17**.

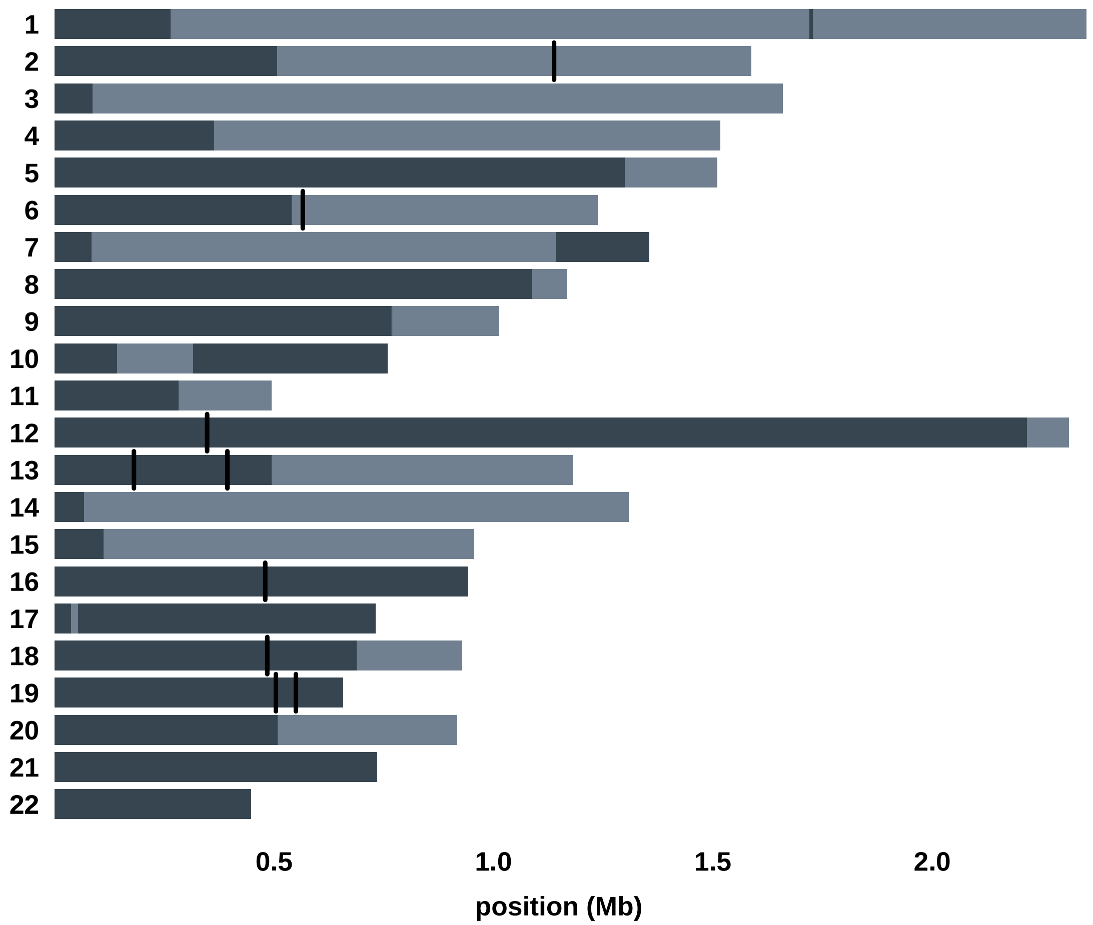


Haplotype blocks observed on the autosomes (chromosomes 1-22) in meiosis 7–10. Haplotype blocks are shown in alternating gray shading (when applicable, gaps are shown in white). NCO events are indicated by vertical black lines.

**Supplementary Figure 18**.

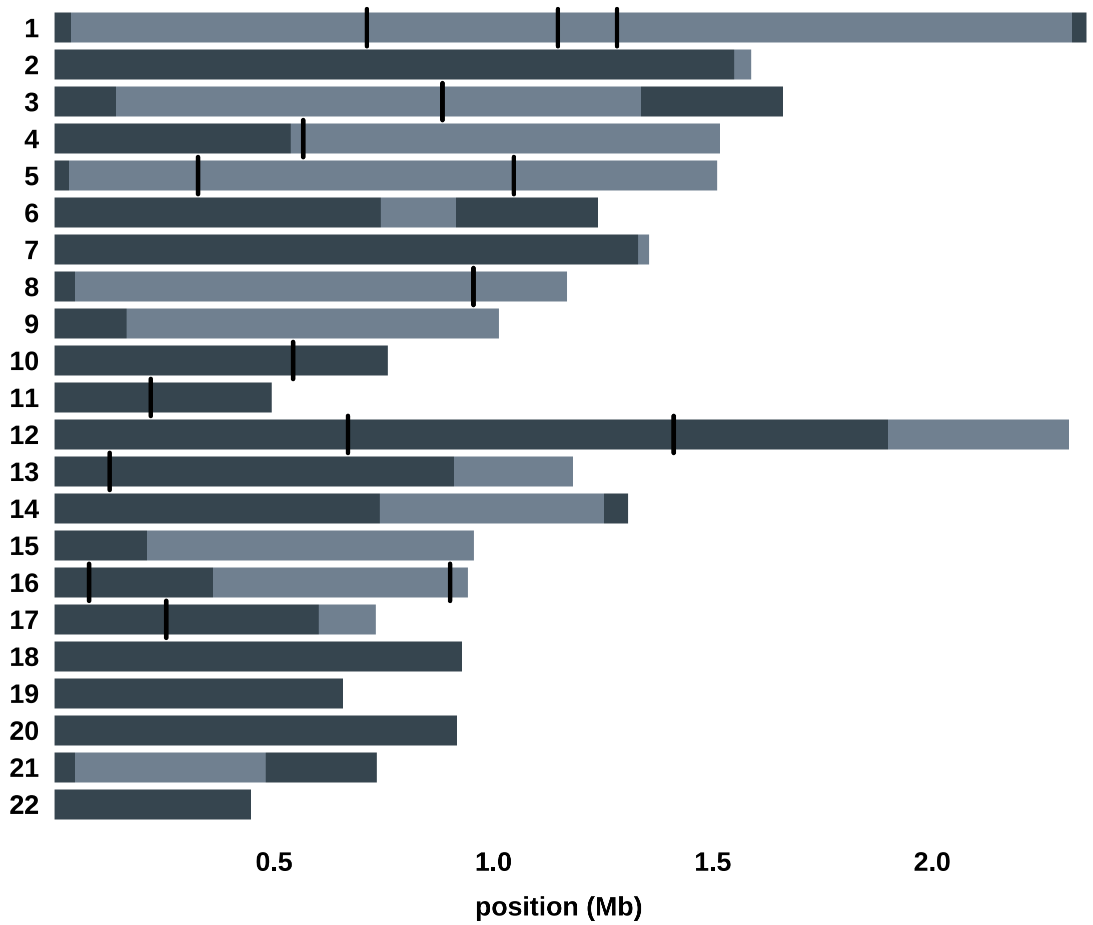


Haplotype blocks observed on the autosomes (chromosomes 1-22) in meiosis 7–11. Haplotype blocks are shown in alternating gray shading (when applicable, gaps are shown in white). NCO events are indicated by vertical black lines.

**Supplementary Figure 19**.

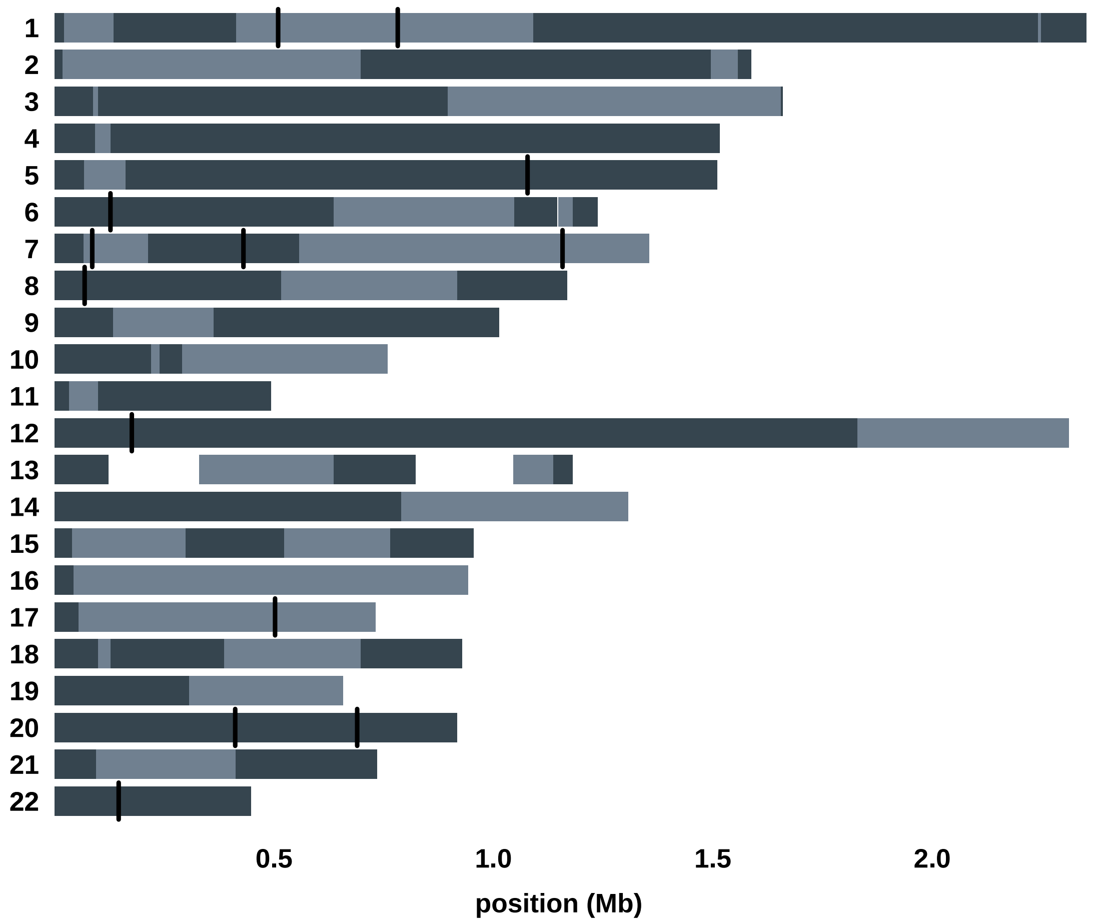


Haplotype blocks observed on the autosomes (chromosomes 1-22) in meiosis 7–12. Haplotype blocks are shown in alternating gray shading (when applicable, gaps are shown in white). NCO events are indicated by vertical black lines.

**Supplementary Figure 20**.

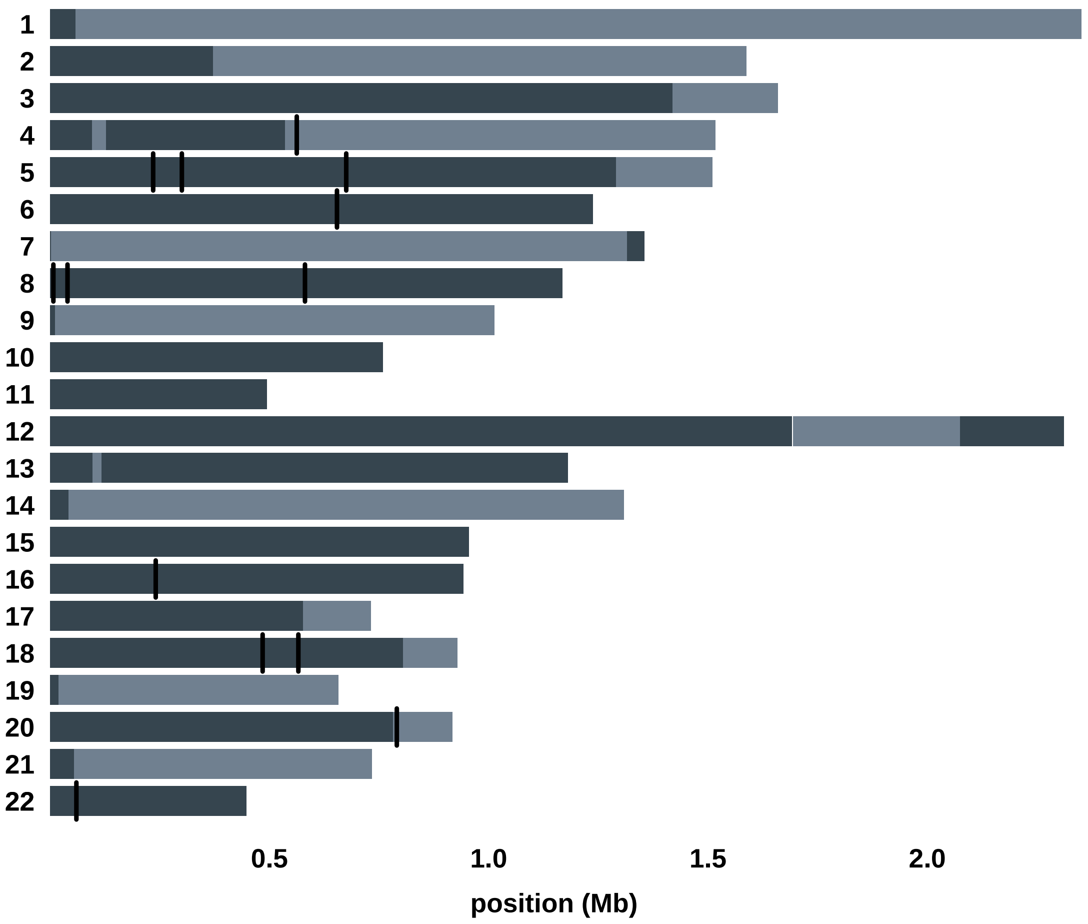


Haplotype blocks observed on the autosomes (chromosomes 1-22) in meiosis 8–9. Haplotype blocks are shown in alternating gray shading (when applicable, gaps are shown in white). NCO events are indicated by vertical black lines.

**Supplementary Figure 21**.

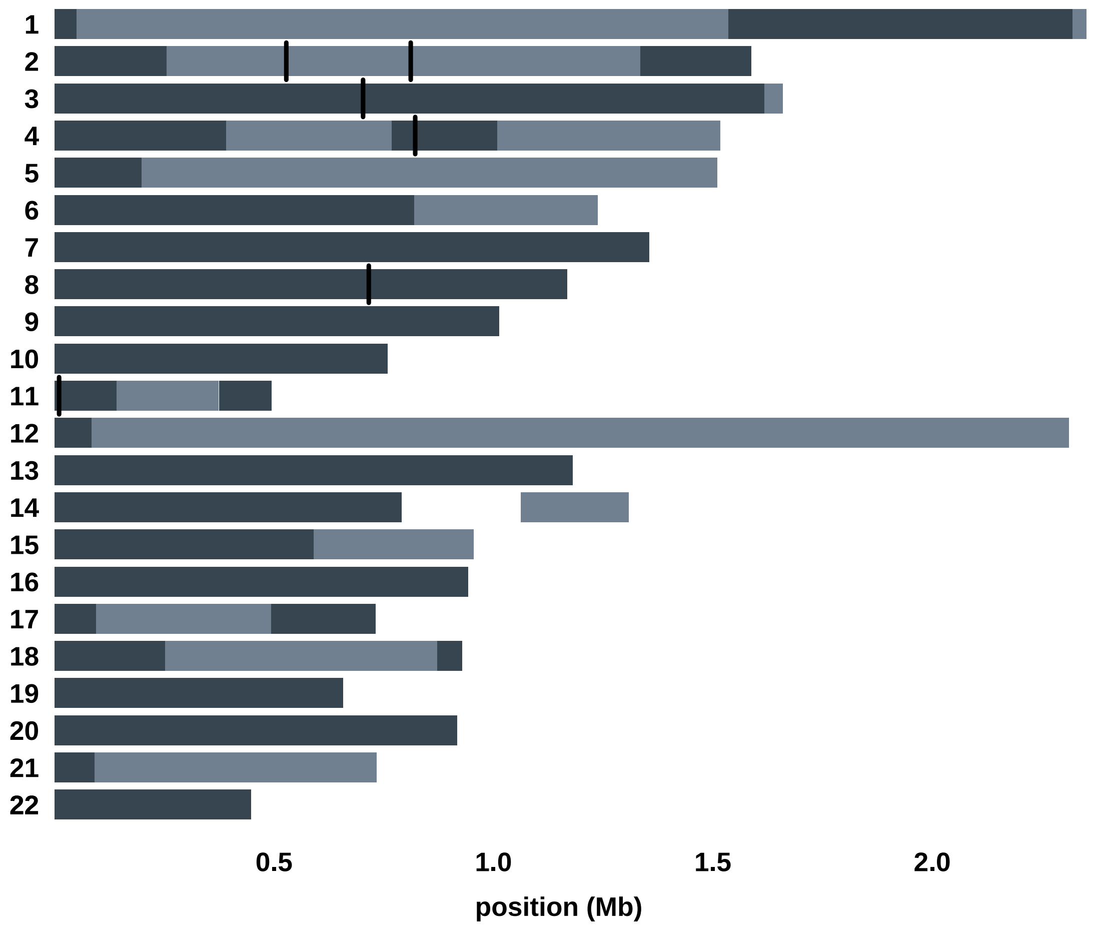


Haplotype blocks observed on the autosomes (chromosomes 1-22) in meiosis 8–10. Haplotype blocks are shown in alternating gray shading (when applicable, gaps are shown in white). NCO events are indicated by vertical black lines.

**Supplementary Figure 22**.

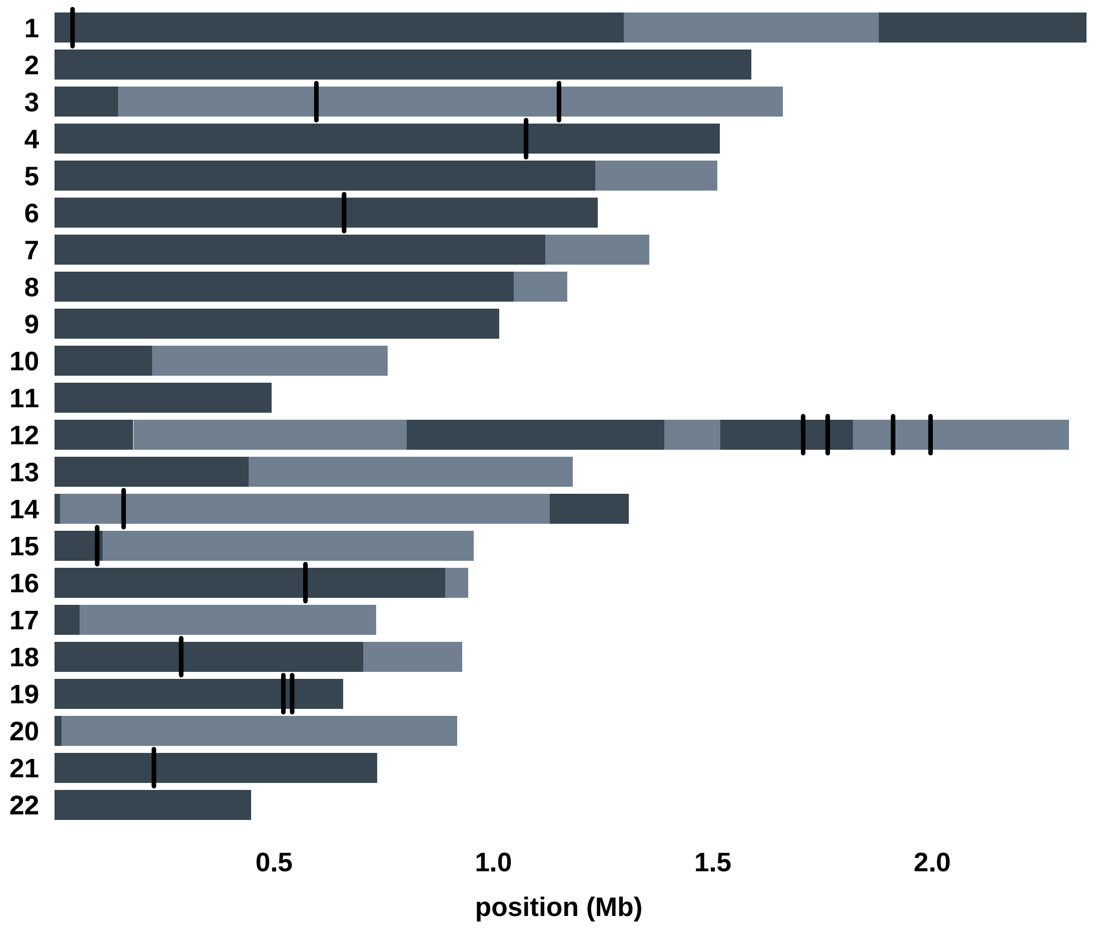


Haplotype blocks observed on the autosomes (chromosomes 1-22) in meiosis 8–11. Haplotype blocks are shown in alternating gray shading (when applicable, gaps are shown in white). NCO events are indicated by vertical black lines.

**Supplementary Figure 23**.

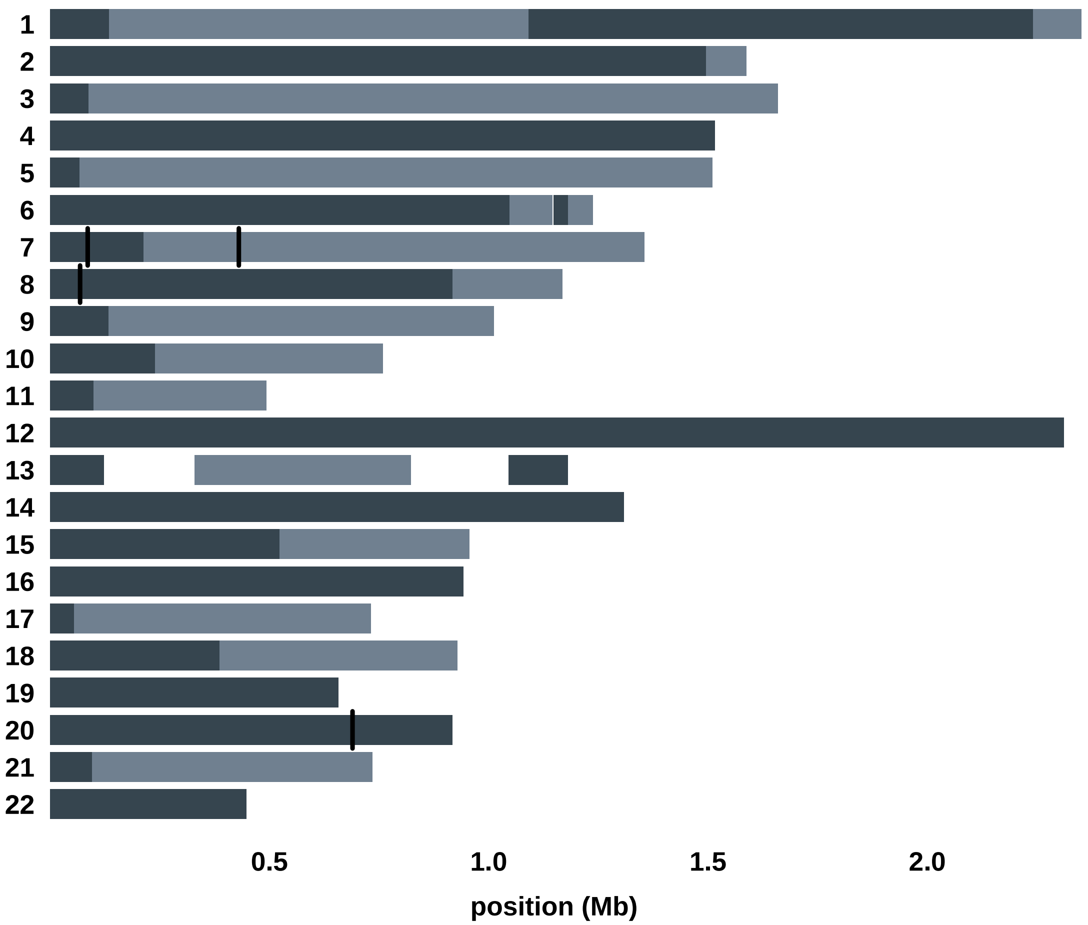


Haplotype blocks observed on the autosomes (chromosomes 1-22) in meiosis 8–12. Haplotype blocks are shown in alternating gray shading (when applicable, gaps are shown in white). NCO events are indicated by vertical black lines.

**Supplementary Figure 24**.

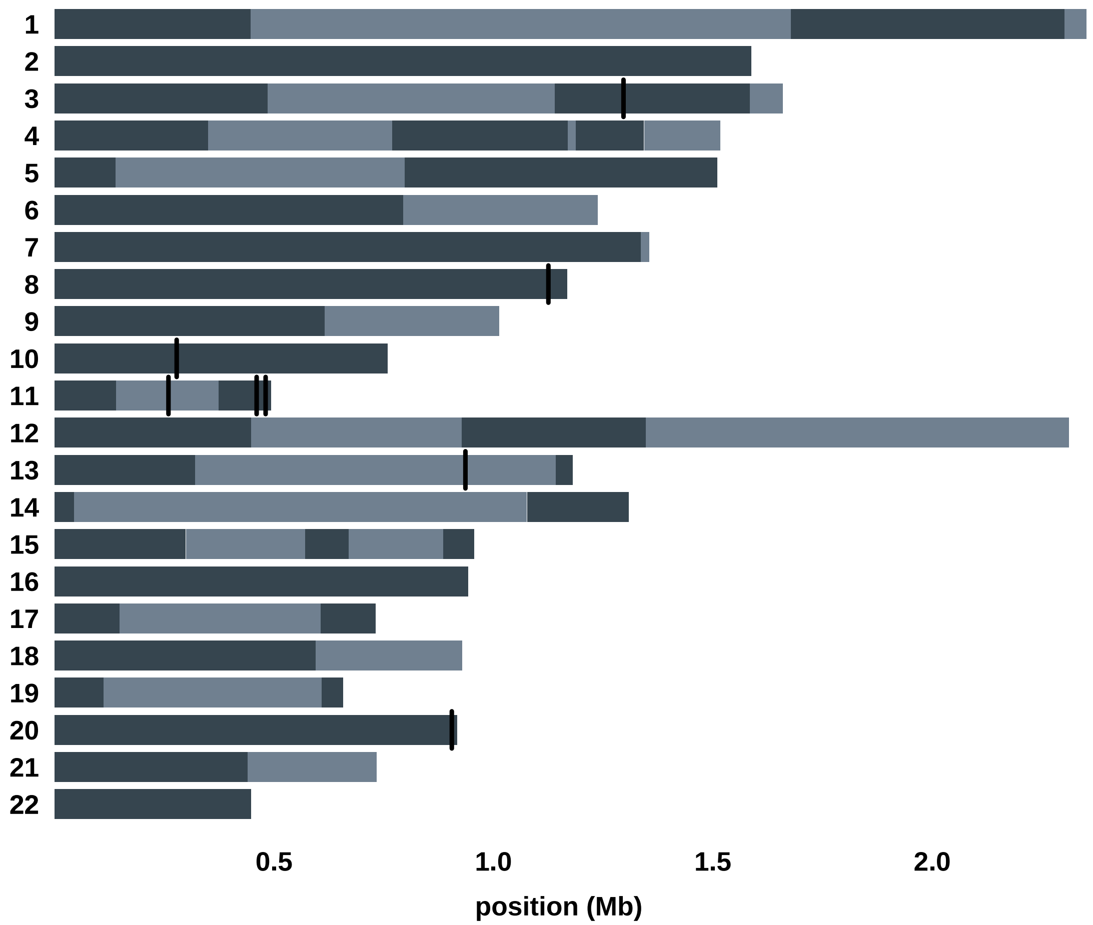


Haplotype blocks observed on the autosomes (chromosomes 1-22) in meiosis 13–15. Haplotype blocks are shown in alternating gray shading (when applicable, gaps are shown in white). NCO events are indicated by vertical black lines.

**Supplementary Figure 25**.

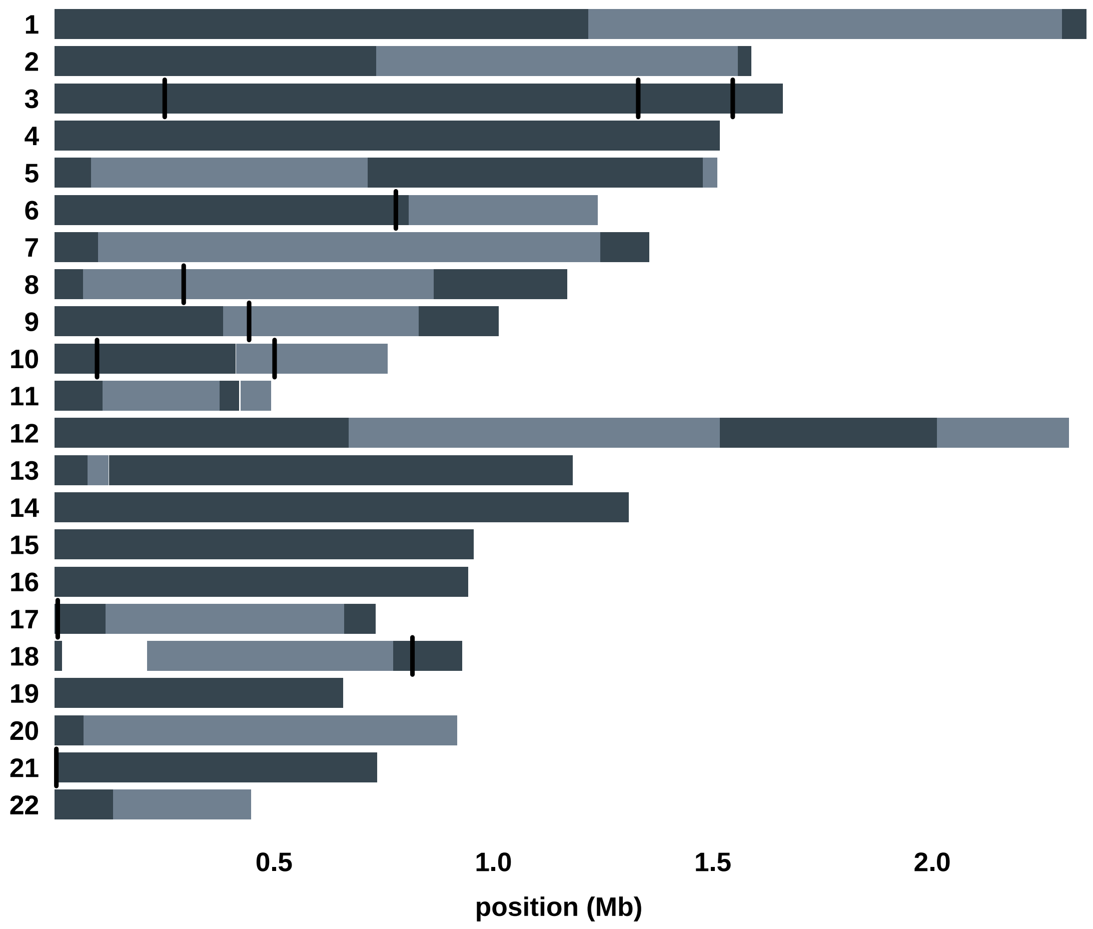


Haplotype blocks observed on the autosomes (chromosomes 1-22) in meiosis 13–16. Haplotype blocks are shown in alternating gray shading (when applicable, gaps are shown in white). NCO events are indicated by vertical black lines.

**Supplementary Figure 26**.

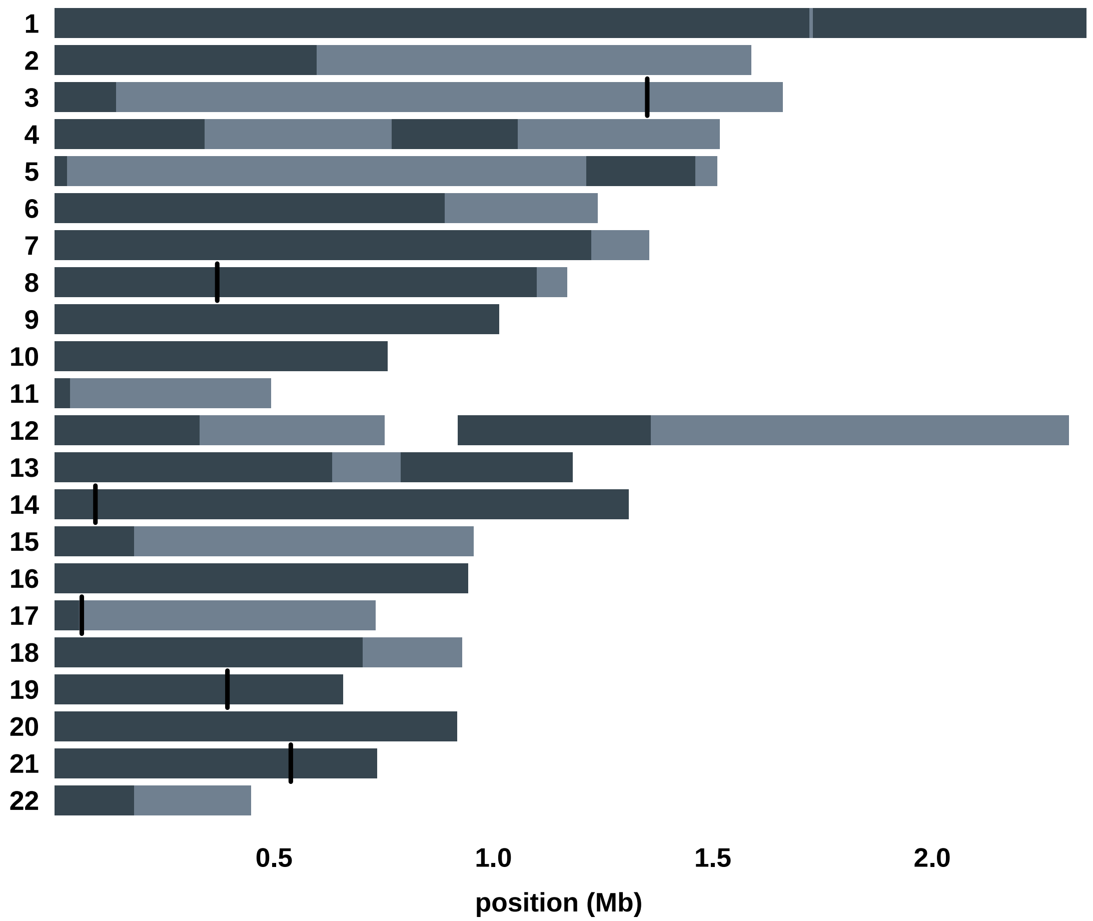


Haplotype blocks observed on the autosomes (chromosomes 1-22) in meiosis 13–17. Haplotype blocks are shown in alternating gray shading (when applicable, gaps are shown in white). NCO events are indicated by vertical black lines.

**Supplementary Figure 27**.

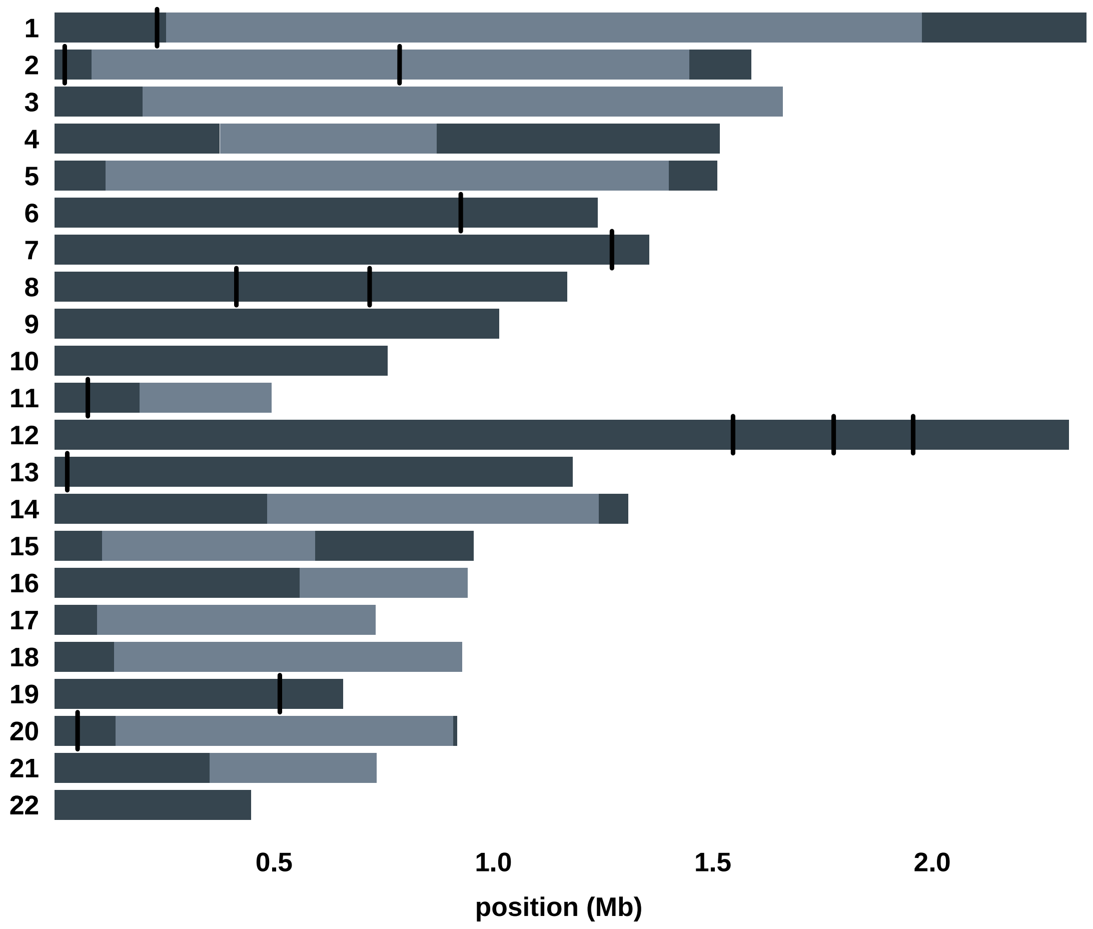


Haplotype blocks observed on the autosomes (chromosomes 1-22) in meiosis 14–15. Haplotype blocks are shown in alternating gray shading (when applicable, gaps are shown in white). NCO events are indicated by vertical black lines.

**Supplementary Figure 28**.

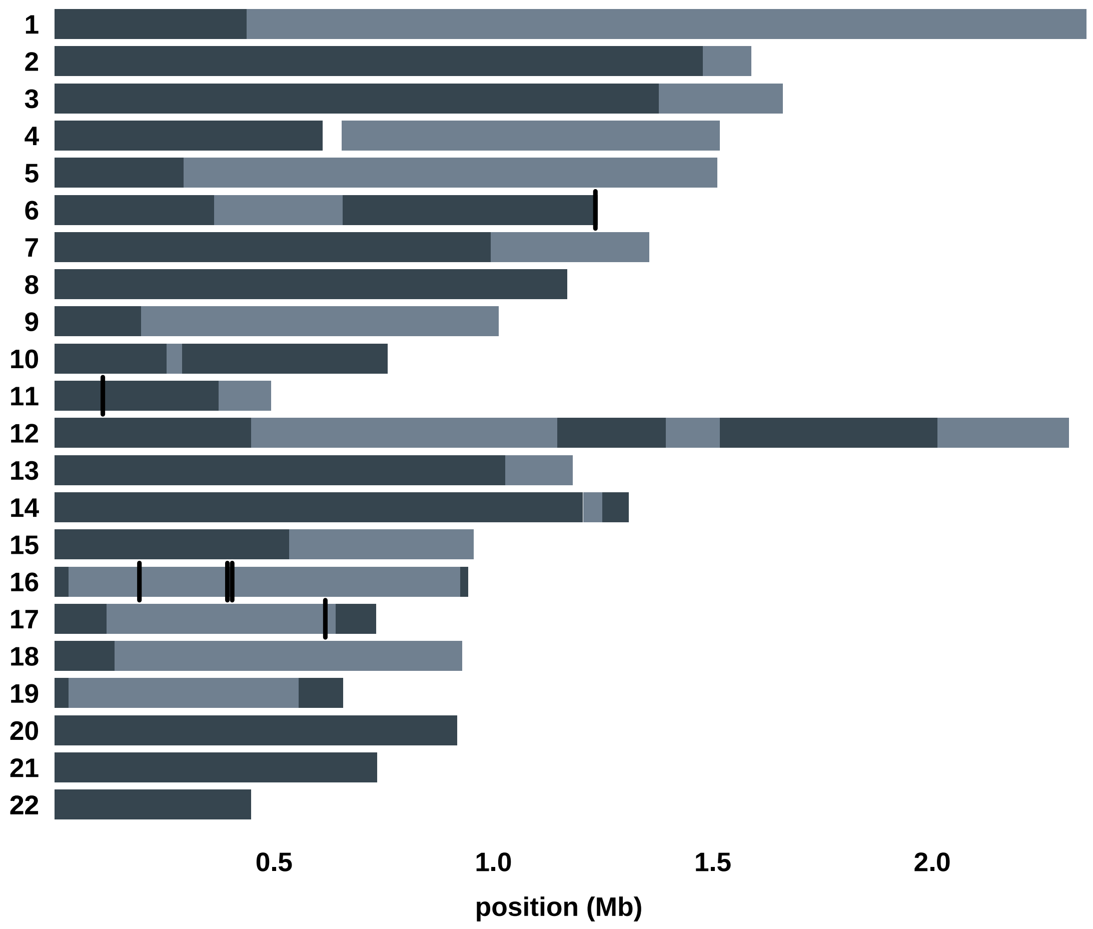


Haplotype blocks observed on the autosomes (chromosomes 1-22) in meiosis 14–16. Haplotype blocks are shown in alternating gray shading (when applicable, gaps are shown in white). NCO events are indicated by vertical black lines.

**Supplementary Figure 29**.

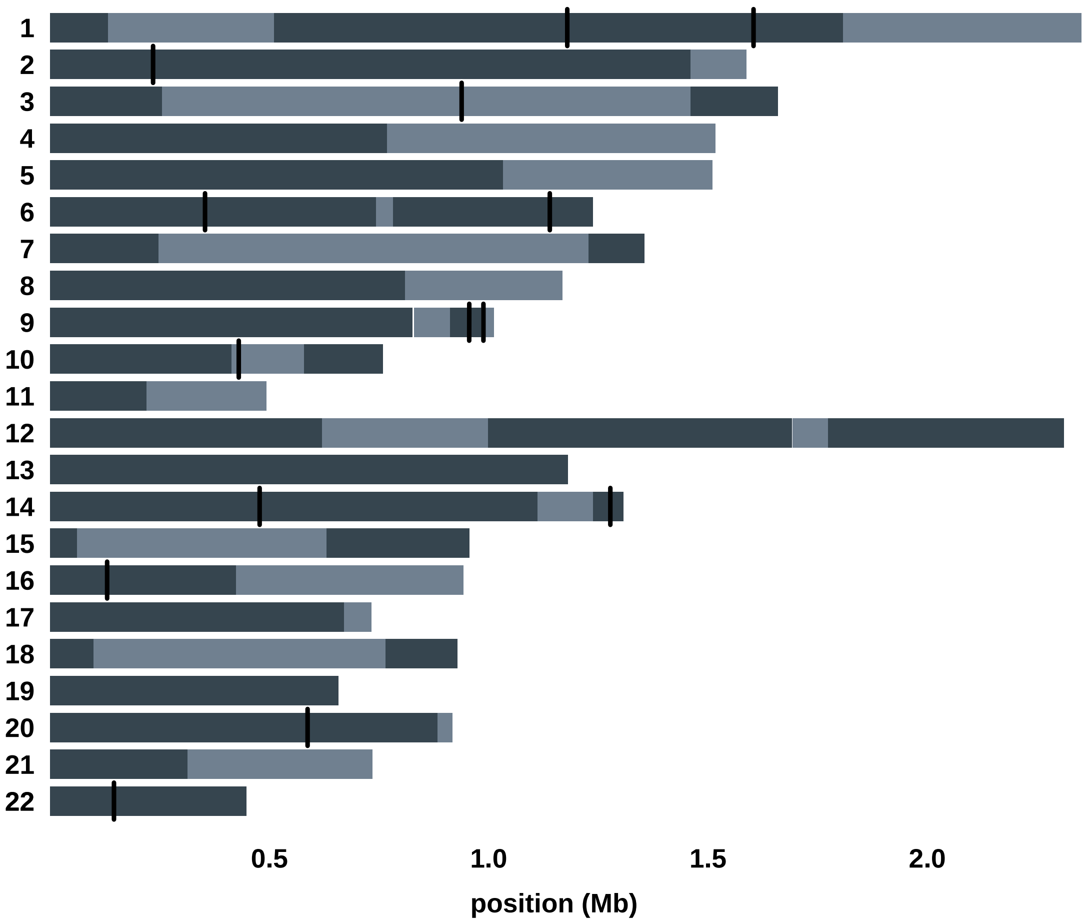


Haplotype blocks observed on the autosomes (chromosomes 1-22) in meiosis 14–17. Haplotype blocks are shown in alternating gray shading (when applicable, gaps are shown in white). NCO events are indicated by vertical black lines.

**Supplementary Figure 30**.

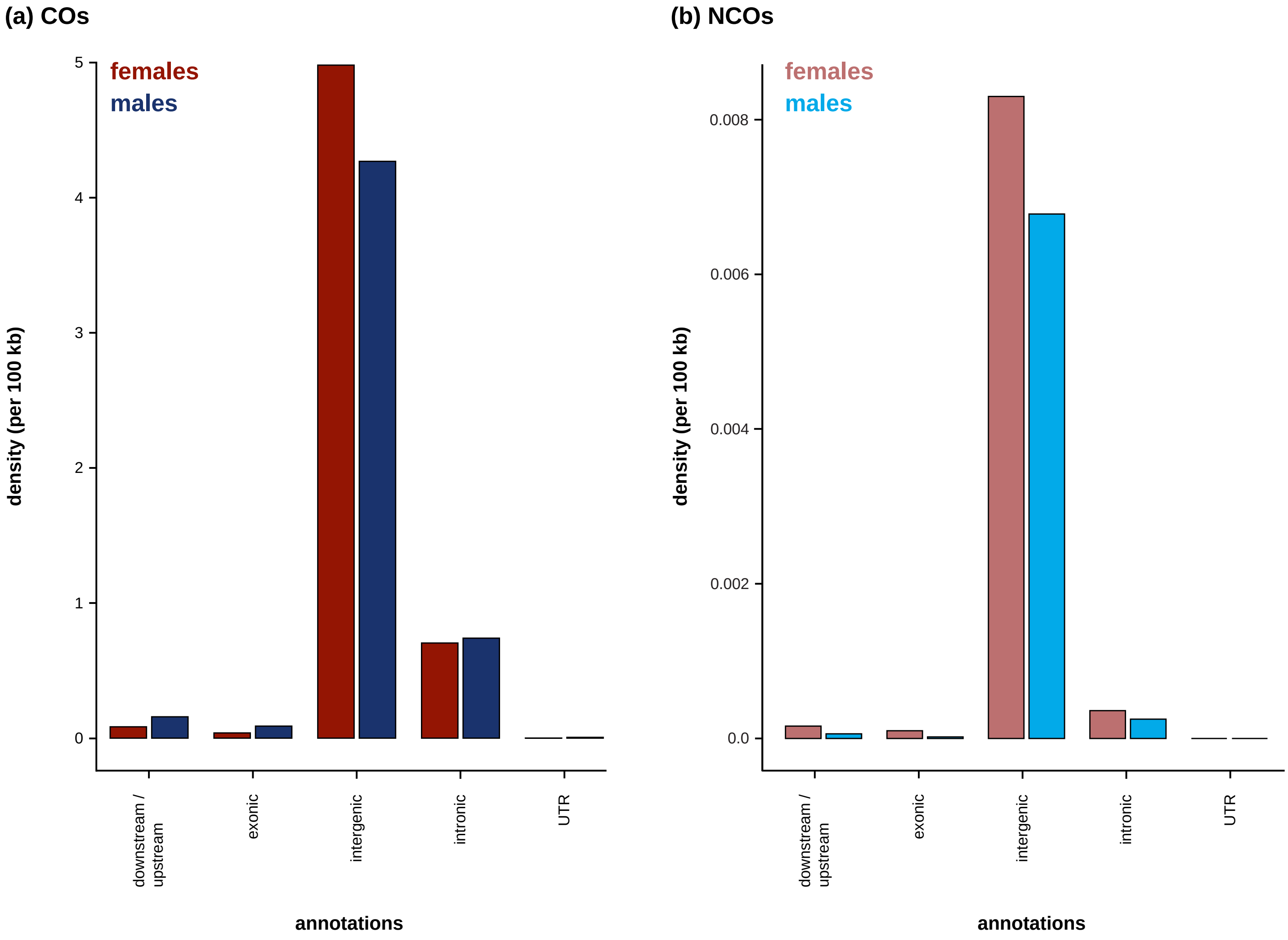


Density of sex-specific (a) COs with a resolution of less than 10 kb and (b) NCOs with a conversion tract length less than 10 kb by genomic region.

**Supplementary Figure 31**.

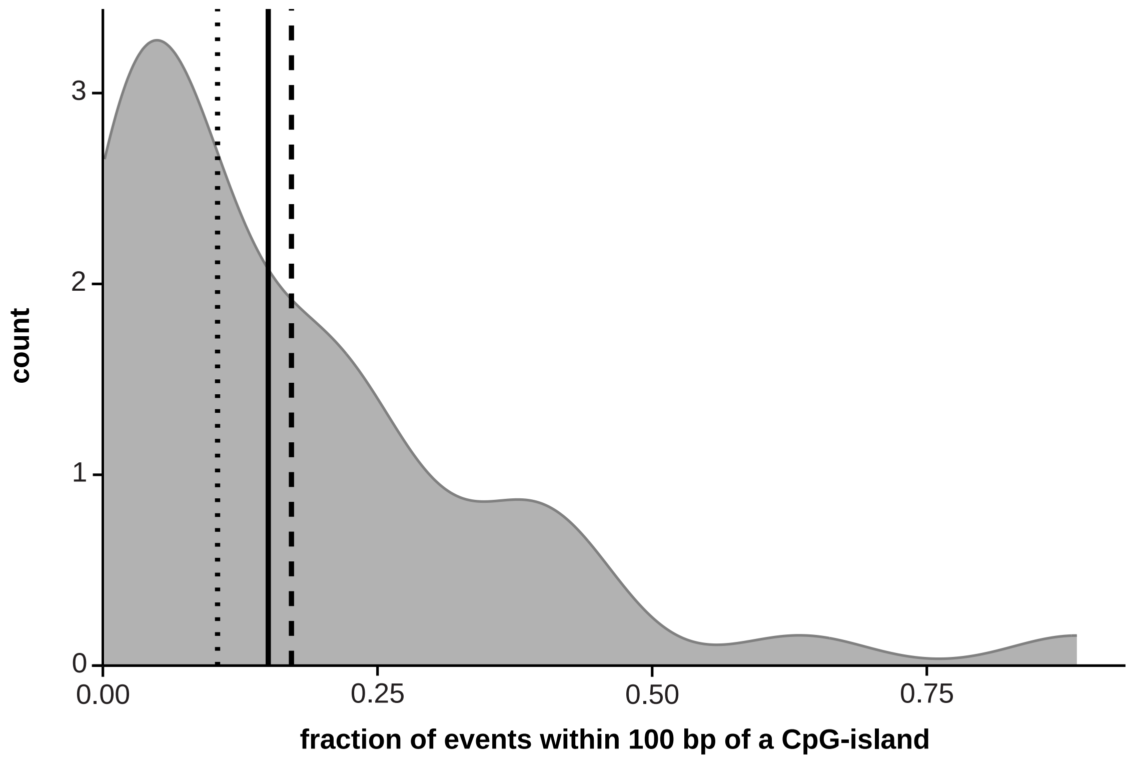


The distribution shown in gray indicates the overlap between CO events and CpG-islands (within 100 bp) that is expected by chance, with the mean and median shown in a black dashed and dotted line, respectively. The empirically observed fraction is shown as a black solid line.

**Supplementary Figure 32**.

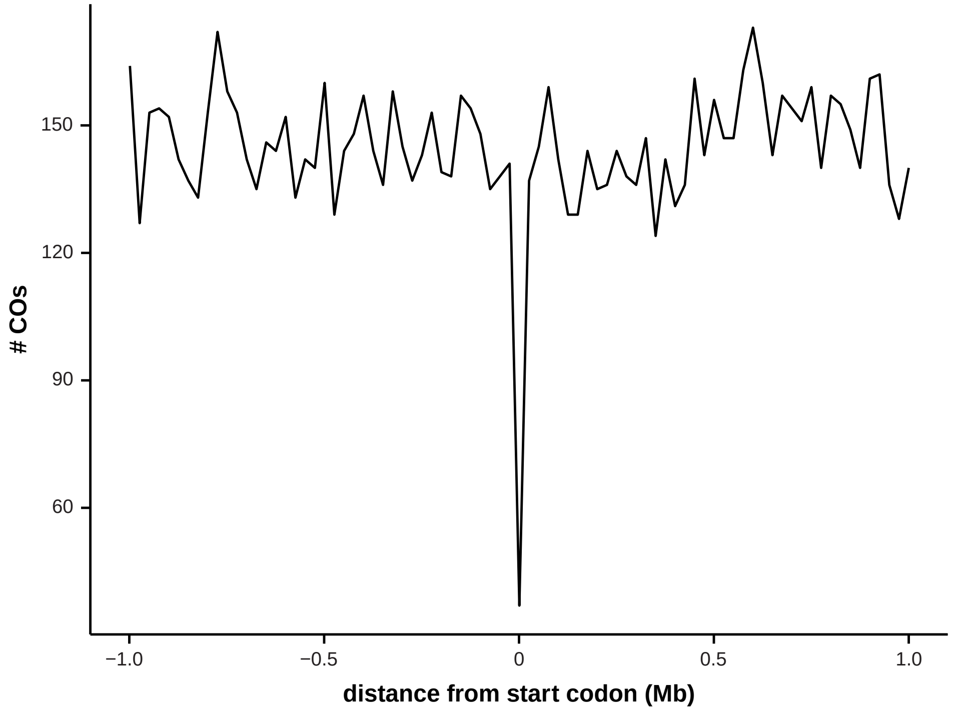


Distribution of the number of COs with a resolution < 10 kb relative to the distance from start codons.

**Supplementary Figure 33**.

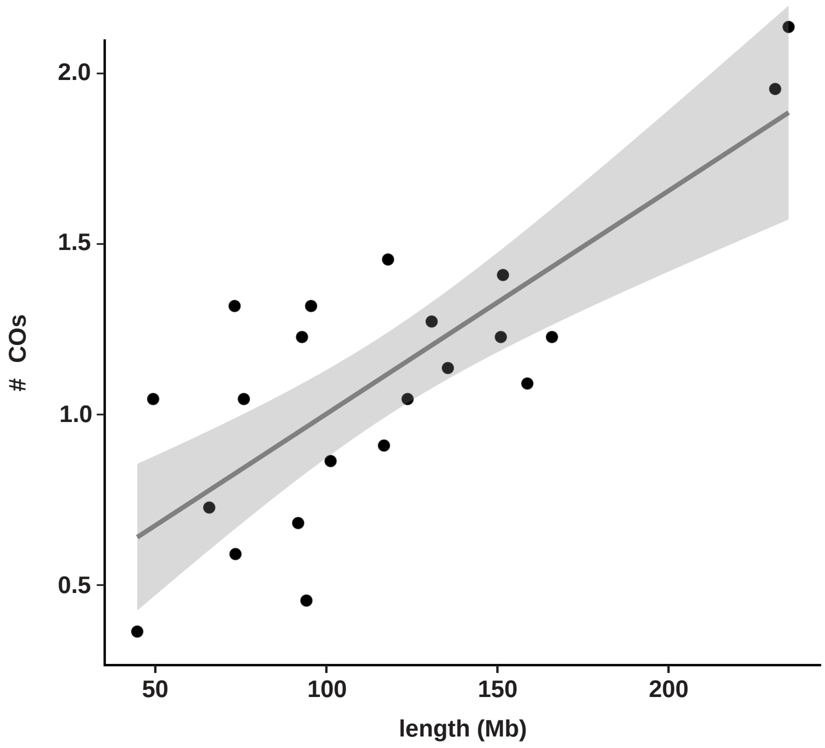


Average number of crossovers (COs) per autosome.

**Supplementary Figure 34**.

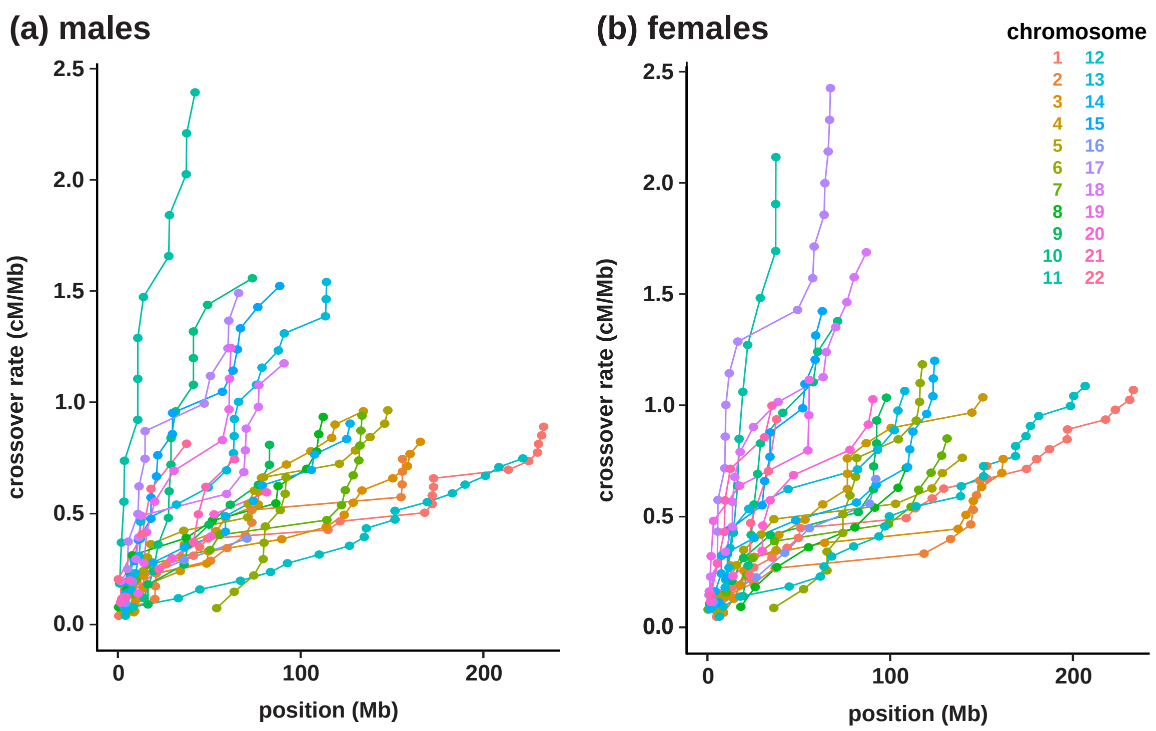


Relationship between the cumulative genetic distance obtained from the distribution of COs and the physical length of each autosome in (a) males and (b) females.

**Supplementary Figure 35**.

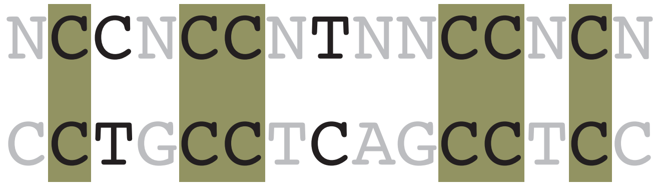


Comparison between the degenerate 13-15-mer motif of the common PRDM9 A allele in humans, (N)CCNCCNTNNCCNC(N) (shown on top), and the identified putative 15-mer PRDM9 binding motif in coppery titi monkeys, CCTGCCTCAGCCTCC (shown at the bottom). Positions shown in gray are degenerate in the human motif while the olive shading highlights the matches of several C/CC nucleotides/dinucleotides.
